# IL-6 prevents Th2 cell polarization by promoting SOCS3-dependent suppression of IL-2 signaling

**DOI:** 10.1101/2022.12.05.519197

**Authors:** Holly Bachus, Erin McLaughlin, Crystal Lewis, Amber M. Papillion, Etty N. Benveniste, Dave Durrell Hill, Alexander F. Rosenberg, André Ballesteros-Tato, Beatriz León

## Abstract

Defective interleukin-6 (IL-6) signaling has been associated with Th2 bias and elevated IgE. However, the underlying mechanism by which IL-6 prevents the development of Th2-driven diseases remains unknown. Using a model of house-dust-mite (HDM)-induced Th2 differentiation and allergic airway inflammation, we show that IL-6 signaling in allergen-specific T cells was required to prevent Th2 development and subsequent IgE response and allergic inflammation. Th2 cell lineage commitment required strong sustained IL-2 signaling. Importantly, we found that IL-6 turned off IL-2 signaling during early T cell activation and thus inhibited Th2 priming. Mechanistically, we found that IL-6-driven inhibition of IL-2 signaling in responding T cells was mediated by upregulation of Suppressor Of Cytokine Signaling 3 (SOCS3). This mechanism could be mimicked by pharmacological Janus Kinase-1 (JAK1) inhibition. Collectively, our results identify an unrecognized mechanism that prevents the development of unwanted Th2 cell responses and associated diseases and outline potential preventive interventions.

## Introduction

Interleukin-6 (IL-6) is a pleiotropic cytokine that plays critical roles in regulating inflammation, hematopoiesis, metabolism, and oncogenesis (1). Classic IL-6 signaling is initiated by the binding of IL-6 to the membrane-bound IL-6-specific receptor α chain (IL-6R). The IL-6/IL-6R complex triggers its association with the signal-transducing subunit, glycoprotein 130 (GP130), leading to the phosphorylation of Janus Kinases (JAK), and Signal Transducers and Activator of Transcription (STAT), which culminate in the nuclear import of phosphorylated STAT dimers, predominantly STAT3, that activate transcription (2). IL-6 signaling is negatively regulated by Suppressors Of Cytokine Signaling (SOCS) proteins, which are generally negative-feedback inhibitors of signaling induced by cytokines that act via the JAK/STAT pathway (3).

Excess IL-6 is central in the pathogenesis of multiple inflammatory conditions such as rheumatoid diseases, cytokine storm, and cytokine release syndrome and targeting of the IL-6 pathway has led to innovative therapeutic approaches for some of these inflammatory diseases (4). On the other hand, loss-of-function mutations that affect IL-6 signaling, including IL6R (5), GP130 (6, 7), and STAT3 (8-11), lead to increased T helper 2 (Th2) bias and manifestation of hallmarks of Th2-mediated immune responses, such as high serum allergen-specific and total IgE concentrations and eosinophilia (12). Consequently, these patients often present with atopy and cutaneous, airway, and systemic manifestations of allergy (13). Although studies of patients have established a role for IL-6 in controlling Th2 bias, the specific contribution of IL-6 and the underlying mechanism remains undefined.

Using a house-dust-mite (HDM)-induced model of Th2-driven allergic airway inflammation, here we show that IL-6 signaling in allergen-specific T cells is required to suppress Th2 cell lineage commitment. Although T-bet has been shown to suppress the Th2 cell-associated program (14, 15), we found that suppression of allergen-specific Th2 cell responses by IL-6 was independent of T-bet. Instead, we found that Th2 cell lineage commitment required strong and prolonged IL-2 signaling, but IL-6 shut down IL-2 signaling during early T cell activation and, thus, inhibited Th2 cell priming. Mechanistically, IL-6 upregulated SOCS3, thus preventing prolonged IL-2 signaling through STAT5 and Th2 cell programming. Furthermore, we found that inhibition of JAK1 with a selective inhibitor could likewise prevent sustained IL-2 signaling and Th2 cell differentiation.

Collectively, our data describe an unrecognized mechanism that prevents the development of harmful Th2 cell responses and allergic disorders. Besides, our data define the specific role of IL-6 signaling in this process. Understanding how IL-6 contributes to allergic sensitization may offer new strategies to prevent atopic disease in patients with deficient IL-6 signaling.

## Results

### IL-6 produced during allergen sensitization is required to suppress allergen-specific Th2 cell responses

To test whether IL-6 influenced Th2 cell response to HDM, we intranasally (i.n.) sensitized and challenged WT and *Il6*^−*/*−^ mice with HDM (**Fig. 1A**) and quantified IL-13/L-5^+^ CD4^+^ Th2 cells in the lungs. We found robust lung accumulation of Th2 cells, but no differences between WT and *Il6*^−*/*−^ mice (**Fig. 1B-D**). In nature, dust allergens are often contaminated with lipopolysaccharide/endotoxin (LPS) at variable levels from ∼10 to 1000 EU/mg (16, 17), with higher levels helping to prevent Th2 cell priming and the development of allergic inflammation (15, 18-21). Our HDM extract contains low endotoxin levels (<30 endotoxin unit (EU)/mg). By adding LPS (1μg LPS per 1mg HDM), we generated high-endotoxin HDM (HDM^LPS^; endotoxin content ∼250 EU/mg) and used it to sensitize mice (**Fig. 1A**). HDM^LPS^ sensitization elevated IL-6 levels in bronchoalveolar lavage fluid (BALF) compared with HDM sensitization (**Fig. S1A**). As expected, sensitization with HDM^LPS^ prevented the accumulation of Th2 cells in the lungs of challenged WT mice (**Fig. 1B-D**). Importantly, however, HDM^LPS^ exposure failed to prevent the increase of Th2 cells in *Il6*^−*/*−^ mice. As a result, Th2 cells largely accumulated in the lung of HDM^LPS^-sensitized *Il6*^−*/*−^ relative to WT mice (**Fig. 1B-D**). These data show that IL-6 is required to prevent the accumulation of allergen-induced Th2 cells. In agreement, while sensitization with HDM^LPS^ prevented Th2-driven eosinophilic airway inflammation (**Fig. 1E-F**) and accumulation of IgE^+^CD138^+^ antibody secreting cells (IgE^+^ ASCs) (**Fig. 1G-I**) in WT challenged mice, it failed to do so in *Il6*^−*/*−^ mice. No differences were observed in lung monocytes or neutrophils (**Fig. 1F and S1B**), although, *Il6*^−*/*−^ mice showed significant less accumulation of Th17 cells in the lung (**Fig. S1C-E**). Our analysis also found no differences in lung regulatory T cells (Tregs) (**Fig. S1F-H**). Similar results were obtained when blocking IL-6 signaling using neutralizing antibodies to IL-6 and IL-6R during sensitization (**Fig. 1J-N and S1I-L**).

**Fig. 1.**
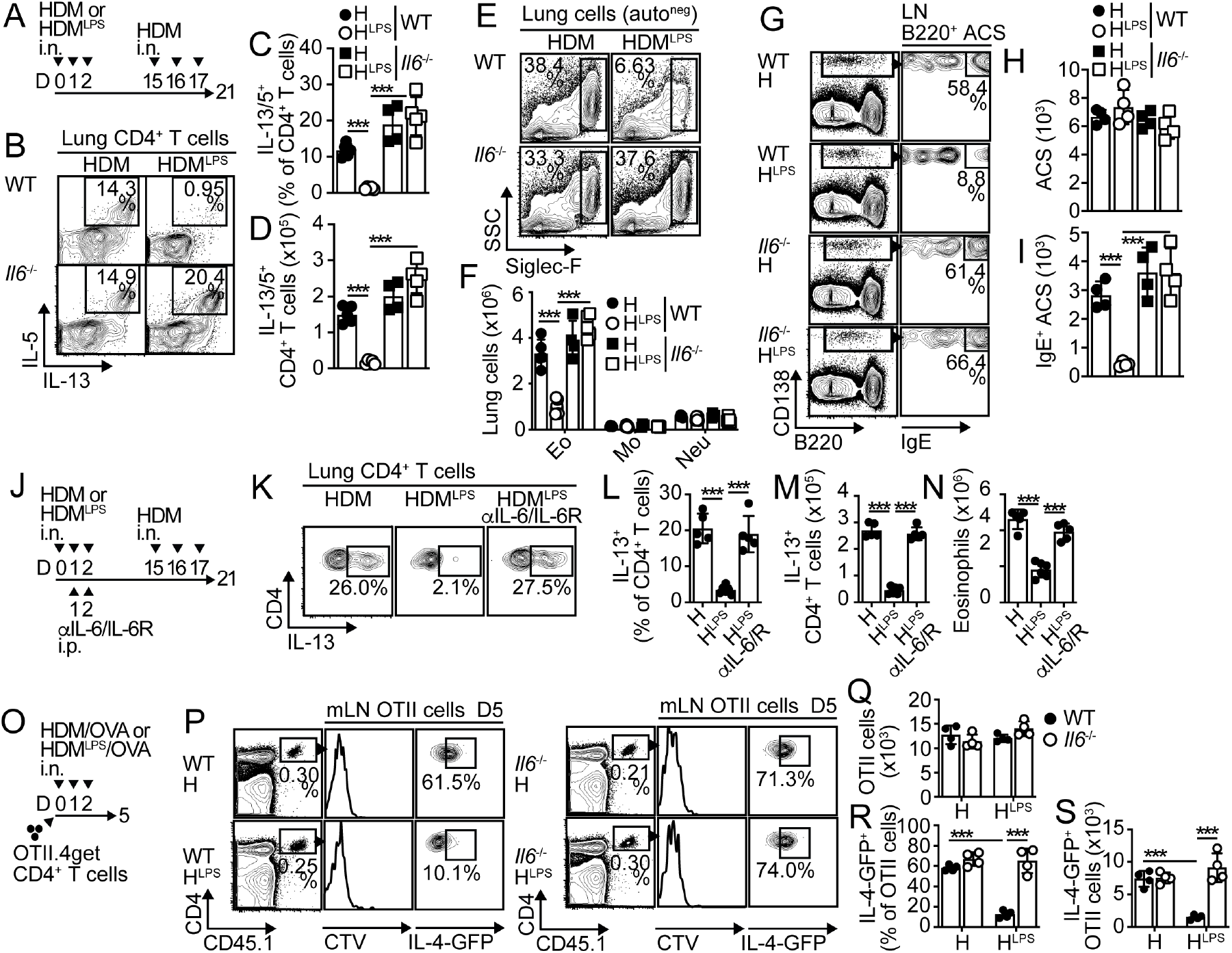
IL-6 signaling during allergen sensitization suppresses allergen-specific Th2 cell-mediated immunity. (**A-I**) B6 (WT) and *Il6*^*-/-*^ mice were i.n. treated with 100μg low-endotoxin HDM (HDM; endotoxin content <30 EU/mg) or high-endotoxin HDM (HDM^LPS^; endotoxin content: 1ug/mg, ∼250 EU/mg) for 3 days. On day 15, mice were i.n challenged with 100μg HDM daily for 3 days and analyzed on day 21 (**A**). Frequencies (**B-C**) and numbers (**D**) of IL-13^+^ IL-5^+^ CD4^+^ T cells in the lungs. Frequencies and numbers of eosinophils, neutrophils and monocytes in the lungs (**E-F**). Frequencies and numbers of total ASCs (**H**) and IgE^+^ ASCs (**G, I**) in the mLN. (**J-N**) B6 mice were i.n. sensitized with HDM or HDM^LPS^. Some mice also received 250μg anti-IL-6 and anti-IL-6R (i.p.). On day 15, mice were i.n challenged with HDM and analyzed on day 21 (**J**). Frequencies (**K-L**) and numbers (**M**) of IL-13^+^ CD4^+^ T cells in the lungs. Numbers of lung eosinophils (**N**). (**O-S**) Mice were transferred with OTII.4get cells, i.n treated with HDM or HDM^LPS^ + 5μg OVA for 3 days and analyzed on day 5 (**O**). Frequencies and numbers of total (**P-Q**) and IL-4-GFP^+^ (**P, R-S**) OTII cells in the mLN. Data are representative of at least three independent experiments (mean±S.D., n=4-5, two-way and one-way Anova). See **Fig. S1**.

The development of allergen-specific Th2 effector cells in the lungs requires initial priming of T cells in the lung-draining, mediastinal LN (mLN) (22). To analyze differences in T-cell priming between WT and *Il6*^−*/*−^ mice, we transferred IL-4-GFP reporter (4get) OTII TCR-transgenic CD4^+^ T cells, followed by sensitization with HDM or HDM^LPS^ in the presence of OVA (**Fig. 1O**). No differences in the expansion of the donor OTII cells in the mLN were observed between groups (**Fig. 1P-Q**); nor in the accumulation of endogenous Tregs (**Fig. S1M-O**). As expected, HDM^LPS^ sensitization prevented the accumulation of IL-4-GFP^+^4get.OTII cells in WT recipients. In contrast, HDM^LPS^ did not prevent the increase of GFP^+^CD4^+^ T cells in *Il6*^−*/*−^ mice **(Fig. 1P** and **1R-S)**. Consequently, GFP^+^4get.OTII cells failed to accumulate in the lungs of HDM^LPS^-sensitized WT mice after challenge, but largely accumulated in the lungs of HDM^LPS^-sensitized *Il6*^−*/*−^ mice (**Fig. S1Q-T)**. Taken together, these data indicate that IL-6, which is produced in response to endotoxin contaminating HDM, is required to prevent the development of specific Th2 cell responses and subsequent allergic inflammation.

### IL-6 signaling in allergen-specific T cells is required to suppress Th2 lineage commitment

We next determined whether IL-6 signaling in T cells was required for suppression of Th2 cell responses. Thus, we sensitized and challenged *Lck*^*cre*^*-Il6r*^*fl/fl*^ and control mice and analyzed cytokine production by T cells in the lungs. IL-13/L-5^+^ Th2 cells failed to accumulate in the lung of control mice that were sensitized with HDM^LPS^. Importantly, however, HDM^LPS^ exposure was unable to prevent the expansion of Th2 cells in *Lck*^*cre*^*-Il6r*^*fl/fl*^ mice (**Fig. 2A-C**). Consequently, eosinophilia (**Fig. 2D-F**) and IgE^+^ASC responses (**Fig. 2G-J**) were suppressed in HDM^LPS^-treated control but not in *Lck*^*cre*^*-Il6r*^*fl/fl*^ mice. Our model of sensitization and challenge did not induce significant numbers of IFNγ^+^CD4^+^ T cells in the lungs, although it induced IL-17 production by T cells (**Fig. 2SA-E**). We found that CD4^+^ T cells from *Lck*^*cre*^*-Il6r*^*fl/fl*^ mice produced significantly less IL-17 (**Fig. S2A-C**), but no differences in lung neutrophilic or monocytic inflammation were observed between *Lck*^*cre*^*-Il6r*^*fl/fl*^ and control mice (**Fig. 2E-F** and **S2F**). IL-6 has been identified as a master regulator of IL-21 in T cells (23, 24). Thus, we studied whether IL-6-driven inhibition of Th2 cell responses was mediated by secondary IL-21 production and signaling. WT and *Il21r*^−*/*−^ mice were sensitized and challenged, and the T cell and inflammatory response were analyzed in the lung. HDM^LPS^ sensitization similarly prevented the accumulation of IL-13/L-5^+^ Th2 cells (**Fig. S2G-I**) and eosinophils (**Fig. S2J-L**) in WT and *Il21r*^−*/*−^ mice. Further, no differences were found in IL-17 production by T cells (**Fig. S2M-O**). Thus, IL-21 signaling is not required to prevent the development of Th2 cell responses to HDM, while, IL-6 signaling is.

**Fig. 2.**
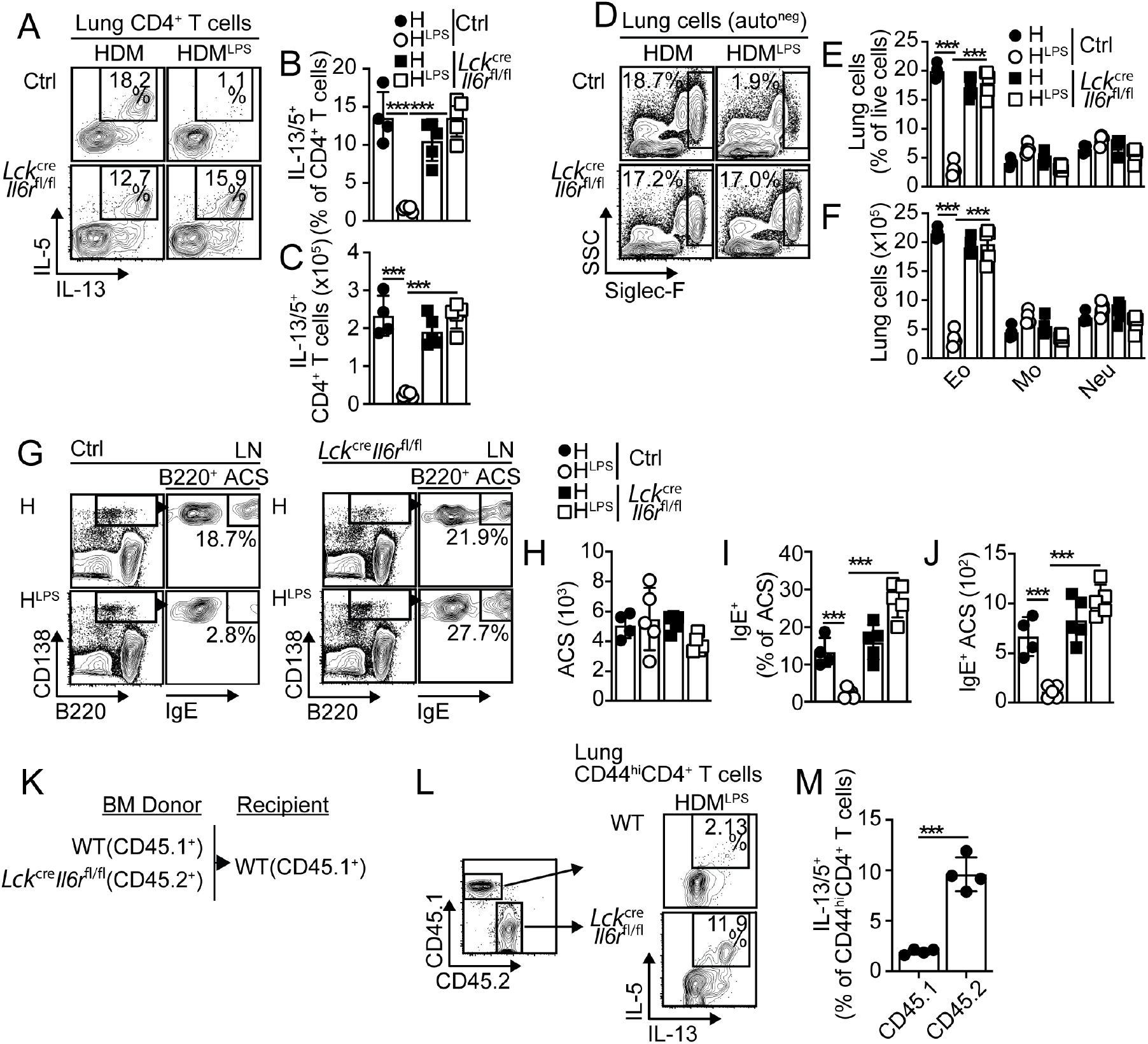
IL-6 signaling in allergen-specific T cells suppresses Th2 cell polarization. (**A-J**) *Lck*^*cre*^*-Il6r*^*fl/fl*^ and control mice were i.n sensitized with HDM or HDM^LPS^ and challenged with HDM. Frequencies (**A-B**) and numbers (**C**) of IL-13^+^ IL-5^+^ CD4^+^ T cells in the lungs. Frequencies and numbers of eosinophils, neutrophils and monocytes in the lungs (**D-F**). Frequencies and numbers of total ASCs (**H**) and IgE^+^ ASCs (**G, I-J**) in the mLN. (**K-M**) Irradiated B6 (CD45.1^+^) mice were reconstituted with 1:1 BM mix of B6 (CD45.1^+^) and *Lck*^*cre*^*-Il6r*^*fl/fl*^ (CD45.2^+^) donors (**K**). Chimeras were i.n sensitized with HDM^LPS^ and challenged with HDM. Frequencies of IL-13^+^ IL-5^+^ WT and *Il6r*^−/−^ CD4^+^ T cells in the lungs (**L-M**). Data are representative of at least three independent experiments (mean±S.D., n=4-5, two-way Anova and unpaired Student’s t test). See **Fig. S2**.

We finally assessed whether the requirement for IL-6 signaling was intrinsically necessary in HDM-responder CD4^+^ T cells. To do this, we sensitized WT:*Lck*^*cre*^*-Il6r*^*fl/fl*^ mixed bone marrow (BM) chimeras (**Fig. 2K**) with HDM^LPS^ and challenged them with HDM. We determined the frequency of IL-13/L-5^+^ Th2 cells within the WT and *Il6r*^−/−^ CD44^hi^CD4^+^ T cell compartments (**Fig. 2L**) and found that Th2 cells largely accumulate in the *Il6r*^−*/*−^ compared to the WT compartment (**Fig. 2L-M**). These results indicated that IL-6 signaling in responder T cells was intrinsically required to suppress the Th2 cell differentiation program.

### Suppression of allergen-specific Th2 cell responses by IL-6 is T-bet independent

T-bet induction by responding T cells is required to suppress the Th2 cell differentiation program to HDM (15). Thus, we tested whether IL-6 signaling suppressed Th2 cell differentiation by up-regulating T-bet. We transferred WT (CD45.1^+^) or Tbx21^−/−^ (CD45.1^+^) 4get.OT-II cells into CD45.2^+^ WT and *Il6*^−*/*−^ recipients, sensitized them and analyzed the progeny from the WT and *Tbx21*^−*/*−^ donors (**Fig. 3A**). OT-II cells expanded similarly in all the groups (**Fig. 3B-C**). As expected, the frequency of GFP^+^ cells within the WT 4get.OT-II from WT recipients decreased in the mLN of HDM^LPS^-relative to HDM-sensitized mice (**Fig. 3B and 3D**). In contrast, the frequency of GFP^+^ cells within the *Tbx21*^−*/*−^ 4get.OT-II was equivalent in both groups (**Fig. 3B and 3D**). Similarly, the frequency of GFP^+^ within the WT 4get.OT-II from *Il6*^−*/*−^ recipients did not decrease after HDM^LPS^ treatment. We observed, however, that the frequency of GFP^+^ cells within the *Tbx21*^−*/*−^ 4get.OT-II from HDM^LPS^-sensitized *Il6*^−*/*−^ recipients increased relative to the same donors primed in a WT environment or to WT donors in *Il6*^−*/*−^ recipients (**Fig. 3B and 3D**). Consequently, a greater accumulation of GFP^+^4get.OTII cells was found when T cells could not up-regulate T-bet and could not receive IL-6 signaling than when these deficiencies occurred separately (**Fig. 3E**). These data suggest that T-bet expression and IL-6 signaling play independent roles in suppressing the Th2 cell differentiation program to HDM.

**Fig. 3.**
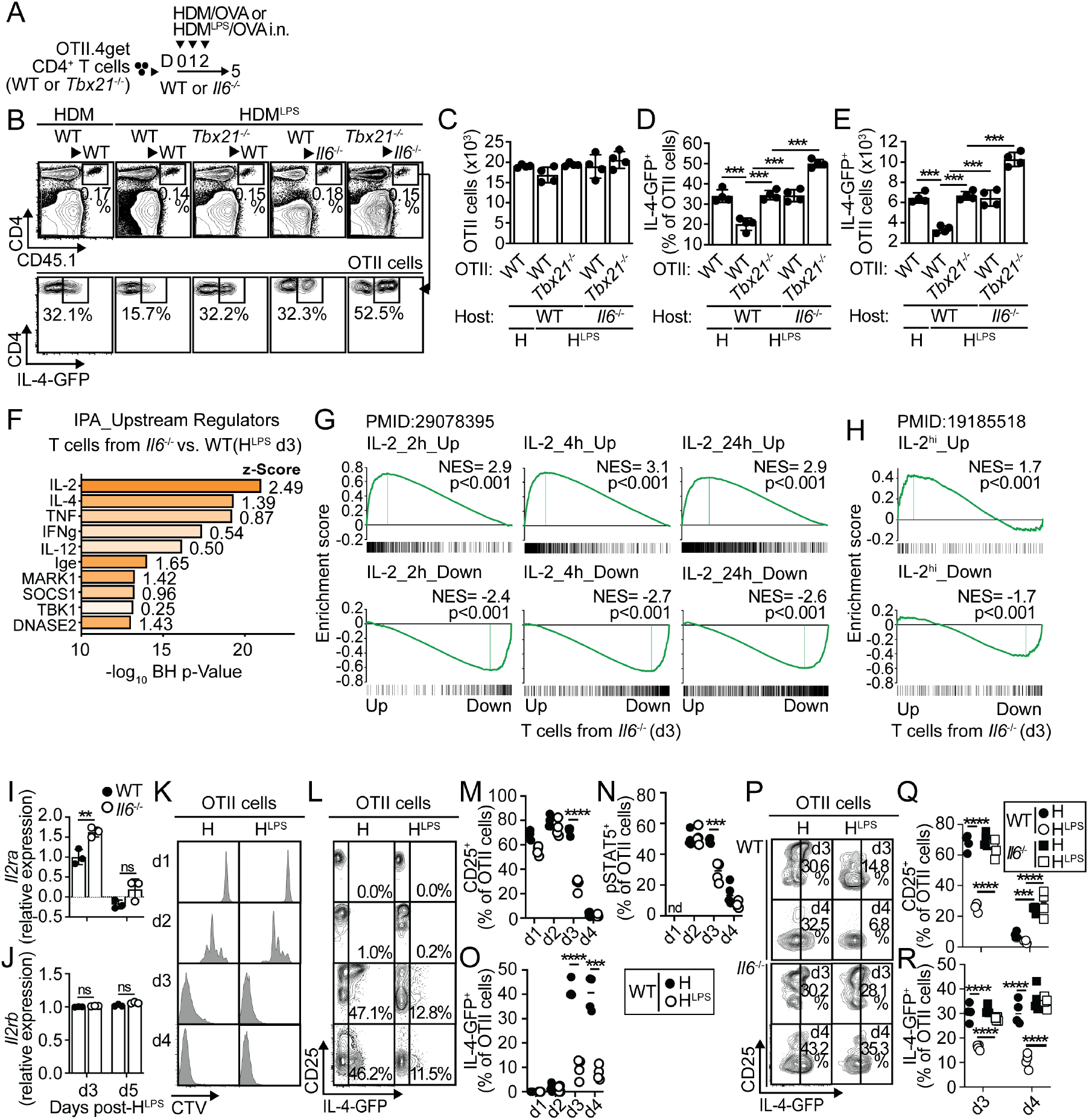
IL-6 signaling in responder T cells prevents prolonged IL-2 responsiveness. (**A-E**) WT and *Il6*^*-/-*^ mice were transferred with WT or *Tbx21*^−/−^ OTII.4get cells, i.n treated with HDM or HDM^LPS^ + OVA for 3 days and analyzed on day 5 (**A**). Frequencies and numbers of total (**B-C**) and IL-4-GFP^+^ (**B, D-E**) OTII cells in the mLN. (**F-H**) WT and *Il6*^*-/-*^ mice were transferred with OTII cells and i.n sensitized with HDM^LPS^+OVA. On day 3, OTII cells were sorted from mLN and RNA-seq was performed (three replicates). 317 differentially expressed genes, with 103 up- and 214 down-regulated in OTII from *Il6*^*-/-*^ mice, were identified (FDR <0.05, −2 FC. See **Table S1**). Activated upstream regulators (positive Z-score) in OTII cells from *Il6*^*-/-*^ versus WT mice based on IPA (**F**). GSEA plots showing the enrichment of genes in OTII cells from *Il6*^*-/-*^ versus WT mice for genes regulated by IL-2 (**G**) or by strong IL-2 signaling (**H**). (**I-J**) Expression of *Il2ra* (**I**) and *Il2rb* in donor OTII cells from HDM^LPS^-treated WT and *Il6*^*-/-*^ mice on days 3 and 5. (**K-R**) WT (**K-R**) and *Il6*^*-/-*^ (**P-R**) mice were transferred with CTV-labeled OTII.4get cells and i.n sensitized with HDM or HDM^LPS^ + OVA. CTV profiles (**K**) and frequencies of CD25 (**M, Q**) and IL-4-GFP (**L, O, P, R**) expression in donor OTII cells from mLNs on different days. Cells from mLNs were stimulated with 1ug/ml rIL-2 for 15 min and STAT5 phosphorylation in OTII cells was determined (**N**). Data are representative of two independent experiments (mean±S.D., n=3-4, one-way and two-way Anova and unpaired Student’s t test). See **Fig. S3 and S4**.

Although IL-6 signaling was found to be required to suppress Th2 cell responses to HDM^LPS^ (LPS content: 1μg/mg HDM), we found that IL-6 (**Fig. S3A-F**) and IL-6 signaling (**Fig. S3G-L**) were dispensable for inhibiting lung Th2 cell accumulation and allergic eosinophilic inflammation when contaminant LPS levels reached 0.4mg LPS per 1mg HDM (HDM^LPS400^). Further, we found that IL-6 was not required to inhibit the lung accumulation of HDM-induced endogenous (**Fig. S3M-N**) or OTII donor (**Fig. S3P-S**) Th2 cells when T cells were primed in an IL-12-rich environment. Thus, IL-6 signaling is particularly necessary to suppress the Th2 cell differentiation program under reduced Th1-polarizing conditions.

### IL-6 suppresses IL-2 signaling early after T cell activation

To understand the role of IL-6 in suppressing Th2 cell commitment, we performed RNA-seq comparing transcriptomes of donor OT-II cells primed in WT and *Il6*^−*/*−^ recipients after sensitization with HDM^LPS^ + OVA. 317 genes were differentially expressed between OTII primed in *Il6*^−*/*−^ and WT recipients on day 3 with FDR < 0.05 and at least a 2-fold change (**Table S1**). Ingenuity Pathway Analysis (IPA) predicted that IL-2 was the regulator with the highest significant enrichment and with the largest activation z-Score (-log (p-value) = 22, Z-score = 2.5) (**Fig. 3F** and **Table S2**), suggesting a role for IL-2 in the expression changes observed in T cells activated in the absence of IL-6. Furthermore, earlier identified kinetic cluster of genes regulated by IL-2 (25) as well as IL-2 highly inducible genes (26) in CD4^+^ T cells were highly enriched in OTII cells primed in *Il6*^−*/*−^ mice (**Fig. 3G-H**), suggesting dominant regulation by IL-2 in the absence of IL-6 signaling. The highest values of normalized enrichment scores (NES) were found on T cells from day 3 compared with day 5 (**Fig. 3G-H** and **S4A-B**), suggesting a most prominent IL-2-dependent regulation early after T cell activation. In agreement, we found that the gene encoding IL-2 receptor α chain (*Il2ra*), which is the top-ranked gene regulated by IL-2 (25), was highly expressed in OTII cells from HDM^LPS^-sensitized *Il6*^−*/*−^ recipients, particularly at day 3, but not day 5 (**Fig. 3I**). No differences were found in the expression of the IL-2 receptor β chain (*Il2rb*) (**Fig. 3J**). These data suggested that IL-6 inhibited IL-2 signaling early after T cell activation and thus prevented IL-2-dependent gene regulation.

To test the prediction that IL-6-dependent inhibition of IL-2 signaling in T cells may be required to suppress the Th2 cell differentiation program to HDM, we first analyzed the kinetics of OTII cell activation in WT recipients following sensitization with HDM or HDM^LPS^ in the presence of OVA. On day 1 after treatment, OTII cells did not yet proliferate (**Fig. 3K**) but strongly upregulated IL-2Rα/CD25 expression (**Fig. 3L-M**), suggesting robust IL-2 signaling. However, no differences were found between sensitization with HDM or HDM^LPS^. On day 2, OTII from HDM- and HDM^LPS^-sensitized mice began proliferating as shown by CellTrace Violet (CTV) dilution (**Fig. 3K**), still maintaining strong responsiveness to IL-2 as indicated by elevated expression of IL-2-driven pSTAT5 (**Fig. 3N**) and CD25 (**Fig. 3L-M**). On day 3, OTII from HDM-sensitized mice began to increase IL-4 expression (**Fig. 3L** and **3O**). OTII cells still retained strong responsiveness to IL-2 (**Fig.. 3L-N**), and indeed, it was the OTII cells expressing the highest levels of CD25 that upregulated IL-4 expression (**Fig. 3L**). In contrast, OTII cells sensitized in the presence of HDM^LPS^ did not prominently upregulated IL-4 (**Fig. 3L** and **3O**), coinciding with the early downregulation of CD25 expression from day 3 (**Fig. 3L-M**) and poor responsiveness to IL-2 (**Fig. 3N**). These results showed that HDM^LPS^ sensitization limited IL-2/STAT5 responsiveness shortly after T cell activation and before Th2 cell commitment. To test whether limited IL-2/STAT5 responsiveness in T cells after HDM^LPS^ sensitization was dependent on IL-6 presence, we performed similar experiments in *Il6*^−*/*−^ recipients. OTII proliferated similarly in WT and *Il6*^−*/*−^ recipients (**Fig. S4C**). However, we found that the early downregulation of CD25 expression (**Fig. 3P-Q**) and the premature attenuation of IL-2-driven pSTAT5 (**Fig. S4D**) observed in OTII cells from WT mice on day 3 post-HDM^LPS^ sensitization did not occur in *Il6*^−*/*−^ recipients. Coincident with this, OTII cells in HDM^LPS^-sensitized *Il6*^−*/*−^ mice did not suppress IL-4 expression (**Fig. 3R**). Our analyses did not find differences in IL-2-driven pSTAT5 response in endogenous Treg cells (**Fig. S4E**). Collectively, these results show that IL-6 limits IL-2 signaling in antigen-specific T cells and suggest that restricting IL-2 signaling is a mechanism to suppress the Th2 cell differentiation program to HDM.

### IL-6 prevents IL-2 responsiveness required for the polarization of naïve T cells to the Th2 cell phenotype

IL-2 plays a critical role in the polarization of Th2 cells (27-29). Consequently, we found that *in vitro* neutralization of IL-2 reduced IL-4-producing cells in CD3/CD28-stimulated 4get.CD4^+^ T cells without altering cell division (**Fig. 4A**). In control conditions, IL-4-GFP was produced by CD25^hi^ CD4^+^ T cells, but anti-IL-2 reduced expression of pSTAT5 and CD25 concurrently with the reduction in IL-4-GFP production (**Fig. 4A-B**). IL-2 neutralization similarly inhibited IL-4-GFP expression by WT (**Fig. 4A-B**) and *Tbx21*^−*/*−^ (**Fig. 4A** and **4C**) CD4^+^ T cells. Comparable results were observed when the strength of IL-2 signaling in T cells was inhibited by blocking CD25 (**Fig. 4D-F**). Thus, strong IL-2 signaling in activated CD4^+^ T cells is required to become IL-4 producers.

**Fig. 4.**
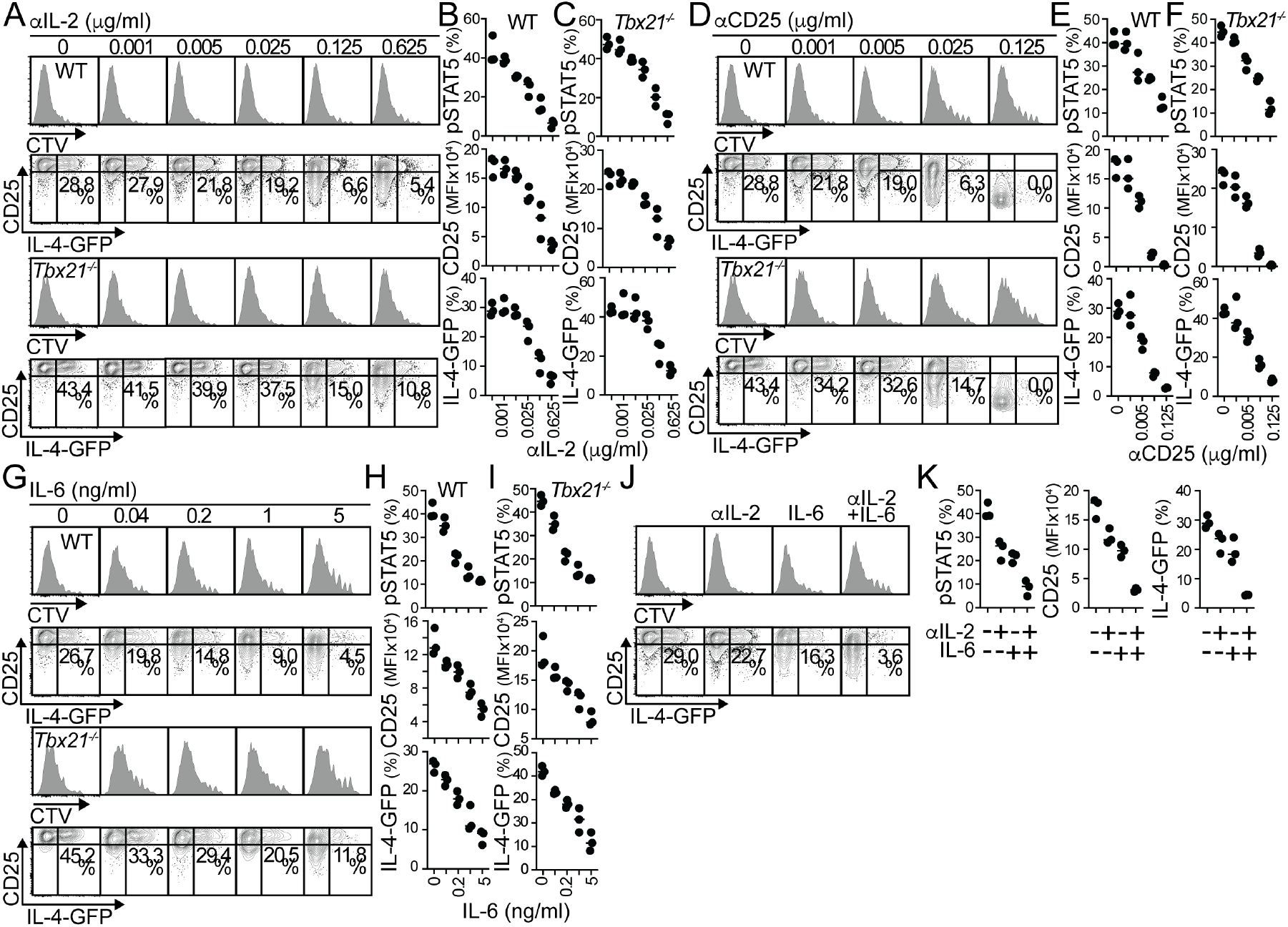
Th2 cell polarization requires strong IL-2 signaling, which is prevented by IL-6. (**A-G**) CTV-labeled WT.4get (CD45.1^+^) and *Tbx21*^−/−^.4get (CD45.2^+^) CD4^+^ T cells from naïve mice were mixed 1:1 and stimulated *in vitro* with plate-bound anti-CD3 and soluble anti-CD28 for 48h. Anti-IL-2 Abs (JES6-1A12 and S4B6), anti-CD25 (PC-61.5.3), or rIL-6 were added at the indicated concentrations for additional 72h. CTV profiles and frequencies of pSTAT5, CD25, and IL-4-GFP expression in WT (**A-B, D-E, G-H**) and *Tbx21*^−/−^ (**A, C, D, F, G, I**) CD4^+^ T cells. (**J-K**) CTV-labeled WT.4get CD4^+^ T cells from naïve mice were activated *in vitro* with plate-bound anti-CD3 and soluble anti-CD28 for 48h. Anti-IL-2 Abs (0.025μg/ml), rIL-6 (0.2ng/ml), or both were added for additional 72h. CTV profiles (**J**) and frequencies of pSTAT5, CD25, and IL-4-GFP expression (**J-K**). Data are representative of four independent experiments. All values were obtained in triplicate, and the data are shown as mean±S.D.

To test whether the presence of IL-6 affected IL-2 responsiveness and IL-4 expression, CD3/CD28-stimulated 4.get.CD4^+^ T cells were cultured in the presence of increasing concentrations of rIL-6. Although CD4^+^ T cells proliferated at a similar rate under all conditions, we found that the addition of IL-6 reduced the expression of pSTAT5, CD25, and IL-4-GFP in WT (**Fig. 4G-H**) and *Tbx21*^−*/*−^ (**Fig. 4G** and **4I**) CD4^+^ T cells. These results indicated that IL-6 signaling prevented IL-2 responsiveness and thus early IL-4 production by activated and proliferating CD4^+^ T cells. These findings also indicated that these IL-6 dependent functions were independent of T-bet expression. To confirm these conclusions, we tested whether the presence of IL-6 affected the threshold of anti-IL-2 required to prevent IL-4 production. The addition of low concentrations of anti-IL-2 or rIL-6 to stimulated 4.get.CD4^+^ T cells moderately decreased the expression of pSTAT5, CD25, and IL-4-GFP **Fig. 4J-K**). However, both conditions together greatly reduced IL-2 responsiveness in T cells and largely suppressed IL-4-GFP expression (**Fig. 4J-K**). Thus, in the presence of IL-6, low concentrations of anti-IL-2 can successfully block response to IL-2 and Th2 cell priming, reinforcing the conclusion that IL-6 controls IL-2 signaling in T cells and, as such, suppresses the priming for Th2 cell differentiation.

### *In vivo* blockade of IL-2 signaling suppresses allergen-specific Th2 cell responses occurring in the absence of IL-6 signaling

We next studied whether *in vivo* inhibition of IL-2 signaling can suppress Th2 cell responses to HDM in low-IL-6 environments. Thus, we blocked the binding of IL-2 to its receptor by administering anti-CD25 mAb to WT:*Lck*^*cre*^*-Il6r*^*fl/fl*^ (*Il6r*^*-/-*^) mixed BM chimeras following initial sensitization (**Fig. 5A-B**). Mice were then analyzed after HDM challenge (**Fig. 5B**). We found similar frequencies and numbers of WT and *Il6r*^*-/-*^ CD44^hi^CD4^+^ T cells in the lungs under all the treatments (**Fig. 5C-D**). As expected, HDM sensitization and challenge induced robust lung accumulation of Th2 cells from both WT and *Il6r*^*-/-*^ donors. However, anti-CD25 treatment prevented this Th2 cell accumulation (**Fig. 5E**), indicating that effector Th2 cell responses to HDM depended on strong IL-2 signaling. Our previous data support that one mechanism by which HDM^LPS^ sensitization suppresses Th2 cell development is by inducing IL-6 and thus suppressing IL-2 responsiveness in allergen-specific T cells. As such, *Il6r*^*-/-*^ T cells cannot suppress Th2 cell differentiation. We found, however, that anti-CD25 treatment could effectively prevent Th2 cell development in *Il6r*^*-/-*^ T cells form HDM^LPS^-sensitized mice (**Fig. 5E**). Treatment with anti-CD25 did not affect Treg numbers (**Fig. S5A**). It also did not affect IL-17 production by activated CD4^+^ T cells, although, as expected, *Il6r*^*-/-*^ T cells produced less IL-17 (**Fig. 5F**). These data show that blocking IL-2 signaling can prevent Th2 cell development in low-IL-6 environments. We confirmed that the effect of blocking IL-2 signaling was on suppressing the priming of antigen-specific Th2 cells by analyzing donor 4get.OTII cells in the mLN of WT and *Il6*^-/-^ recipients after sensitization (**Fig. 5G**). Consistent with our previous results, anti-CD25 mAb administered 24h after sensitization did not affect the expansion of the donor OTII cells (**Fig. 5H-I**) but suppressed their IL-4-GFP^+^ expression and thus the accumulation of IL-4-GFP^+^4get.OTII cells in HDM^LPS^-sensitized *Il6*^-/-^ recipients (**Fig. 5H** and **5J-K**).

**Fig. 5.**
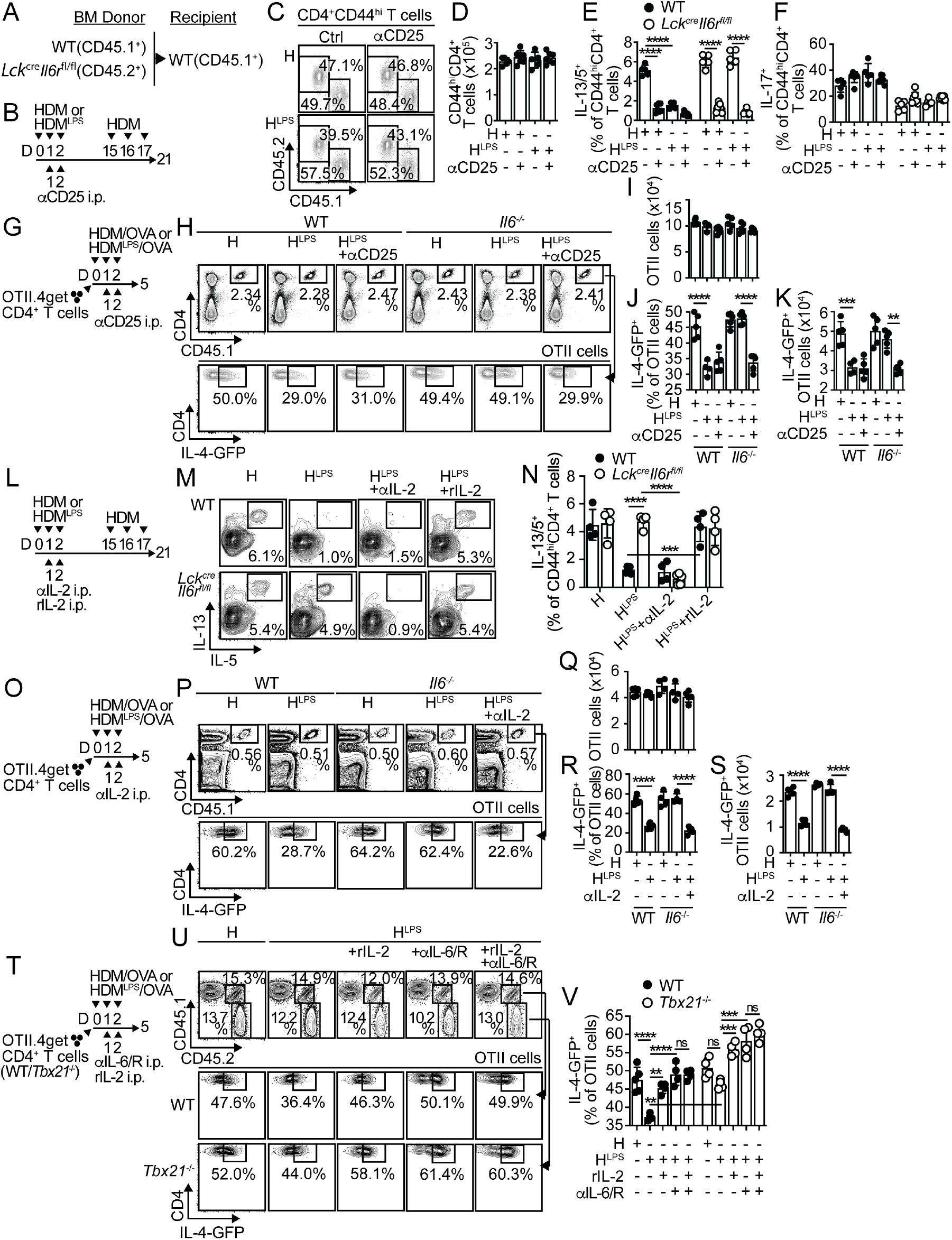
IL-6 and IL-2 signaling on allergen-specific T cells oppositely regulate polarization toward a Th2 profile. (**A-F**) Irradiated B6 (CD45.1^+^) mice were reconstituted with 1:1 BM mix of B6 (CD45.1^+^) and *Lck*^*cre*^*-Il6r*^*fl/fl*^ (CD45.2^+^) donors (**A**). Chimeras were i.n sensitized with HDM or HDM^LPS^. Some mice also received 250μg anti-CD25 (i.p.). Mice were then challenged with HDM (**B**). Frequencies of CD45.1^+^ and CD45.2^+^ cells in CD44^hi^CD4^+^ T cells from the lungs (**C**). Numbers of CD44^hi^CD4^+^ T cells in the lung (**D**). Frequencies of IL-13^+^ IL-5^+^ (**E**) and IL-17^+^ (**F**) cells within the WT and *Lck*^*cre*^*-Il6r*^*fl/fl*^ CD44^hi^CD4^+^ T cell compartments in the lungs. (**G-J**) WT and *Il6*^*-/-*^ mice were transferred with OTII.4get cells, i.n treated with HDM or HDM^LPS^ + OVA, and i.p treated with anti-CD25 or PBS (**G**). Frequencies and numbers of total (**H-I**) and IL-4-GFP^+^ (**H, J-K**) OTII cells in the mLN. (**L-N**) WT:*Lck*^*cre*^*-Il6r*^*fl/fl*^ chimeras were i.n sensitized with HDM or HDM^LPS^, i.p treated with anti-IL-2 Abs (JES6-1A12 and S4B6; 250μg each), rIL-2 (60,000U) or PBS, and i.n challenged with HDM (**L**). Frequencies of IL-13^+^ IL-5^+^ cells within the WT and *Lck*^*cre*^*-Il6r*^*fl/fl*^ CD44^hi^CD4^+^ T cell compartments in the lungs (**M-N**). (**O-S**) WT and *Il6*^*-/-*^ mice were transferred with OTII.4get cells, i.n treated with HDM or HDM^LPS^ + OVA, and i.p treated with anti-IL-2 Abs or PBS (**O**). Frequencies and numbers of total (**P-Q**) and IL-4-GFP^+^ (**P, R-S**) OTII cells in the mLN. (**T-V**) WT (CD45.1^+^) mice were co-transferred with WT (CD45.1^+^CD45.2^+^) and *Tbx21*^−/−^ (CD45.2^+^) OTII.4get cells, i.n treated with HDM or HDM^LPS^ + OVA, and i.p treated with anti-IL-6 and anti-IL-6R Abs, rIL-2 or PBS (**T**). Frequencies of total (**U**) and IL-4-GFP^+^ (**U-V**) WT and *Tbx21*^−/−^ OTII cells in the mLN. Data are representative of two independent experiments (mean±S.D., n=4-6, one-way Anova). See **Fig. S5**.

In addition, we also examined the ability of CD4^+^ T cells that cannot receive normal IL-2 signals to produce cytokines in low- and high-IL-6 environments. Thus, WT:*Il2ra*^*-/-*^ mixed BM chimeras were sensitized, treated with anti-IL-6/IL-6R or control mAb, and analyzed after challenge (**Fig. S5B-C**). We found less pulmonary accumulation of CD44^hi^CD4^+^ T cells from *Il2ra*^*-/-*^ than WT donors but a similar proportion of naïve CD44^lo^CD4^+^ T cells (**Fig. S5D-E**), indicating a known role for IL-2 signaling in T cell expansion. Since our data showed that IL-2Rα/CD25 neutralization 24 hours after initial sensitization had no significant impact on the number of T cells activated in the mLN (**Fig. 5G-I**) and recruited to the lung (**Fig. 5B-D**), this could indicate that IL-2 signaling within the first few hours after T cell activation is most crucial for expansion. We analyzed cytokine production by activated CD44^hi^CD4^+^ T cells in the lung and found that *Il2ra*^*-/-*^ T cells had impaired capacity to produce Th2 cell cytokines even in low-IL-6 environments (**Fig. S5F-G**). However, they were perfectly able to produce IL-17 at levels comparable to WT T cells (**Fig. S5H-I**). Thus, inhibition of IL-2 signaling in allergen-stimulated T cells can prevent the Th2 cell fate determination favored in low-IL-6 environments or in T cells that do not respond to IL-6.

### The balance between IL-6 and IL-2 signals regulates the development of allergen-specific Th2 cell responses

We next decided to change the availability of IL-2 during the sensitization phase in CD4^+^ T cells that did or did not receive IL-6 signaling. Thus, we treated WT:*Lck*^*cre*^*-Il6r*^*fl/fl*^ mixed BM chimeras with anti-IL-2 mAb or rIL-2 during sensitization to decrease or increase the availability of IL-2, respectively (**Fig. 5L**). Mice were then challenged, and cytokine production by WT and *Il6r*^*-/-*^ T cells were analyzed in the lung. In HDM^LPS^-sensitized mice, treatment with anti-IL-2 mAb prevented Th2 cell cytokine production by *Il6r*^*-/-*^ CD44^hi^CD4^+^ T cells. On the contrary, treatment with rIL-2 promoted Th2 cell cytokine production by WT CD44^hi^CD4^+^ T cells ^+^ T cells (**Fig. 5M-N**). Administration of anti-IL-2 mAb or rIL-2 did not affect the capacity of the T cells to produce IL-17 (**Fig. S5J-K**). These data showed that lung effector Th2 cell responses to HDM promoted by lack of IL-6 signaling could be prevented by neutralizing IL-2. In contrast, excess IL-2 signaling promoted effector Th2 cell responses to HDM in the lung even in the presence of intact IL-6 signaling.

We next evaluated whether the opposite effect of IL-2 and IL-6 on lung Th2 cell responses resulted from different priming of antigen-specific CD4^+^ T cells after sensitization. First, we evaluated the effect of anti-IL-2 mAb in donor 4get.OTII cell expansion and polarization (**Fig. 5O**). Anti-IL-2 treatment did not affect the expansion of the donor OTII cells (**Fig. 5P-Q**) but downregulated CD25 (**Fig. S5L-N**) and suppressed IL-4-GFP^+^ expression (**Fig. 5P** and **5R-S**) in OTII cells from *Il6*^-/-^ recipients treated with HDM^LPS^. Thus, inhibition of IL-2 signaling by IL-2 neutralization prevented the Th2 cell polarization of antigen-specific T cells arising in the absence of IL-6. Next, we evaluated the effect of administering rIL-2 on the activation and polarization of antigen-specific CD4^+^ T cells. These analyses were done in combination with blockade of IL-6 signaling and T-bet deficiency, all of which contribute to enhanced Th2 cell responses. Thus, we co-transferred WT (CD45.1^+^ CD45.2^+^) and Tbx21^−/−^ (CD45.2^+^) 4get.OT-II cells into CD45.1^+^ WT recipients, sensitized and treated them with anti-IL-6/IL-6R or rIL-2, and analyzed the progeny from the WT and *Tbx21*^−*/*−^ donors (**Fig. 5T**). WT and *Tbx21*^−*/*−^ OTII cells expanded similarly in all the conditions (**Fig. 5U**). In WT OTII cells form HDM^LPS^-treated mice, rIL-2 and anti-IL-6/IL-6R treatments enhanced the frequency of IL-4-GFP^+^ cells. The combination of both treatments did not have additive effects. Similar results were observed within *Tbx21*^−*/*−^ OTII from HDM^LPS^ -sensitized mice, although, in this case, the absence of T-bet expression in the OTII cells had an additive effect with rIL-2 or anti-IL-6/IL-6R treatments. Overall, our data show that sustained IL-2 signaling after initial T cell activation, which is particularly favored when T cells receive suboptimal IL-6 signaling, supports Th2 cell polarization of antigen-specific T cells by a mechanism independent of T-bet. Therefore, maintaining a proper balance between IL-6 and IL-2 appears to be crucial in controlling Th2 cell immunity to allergens.

### IL-6 prevents IL-2 responsiveness by inducing SOCS3 expression

To better define the mechanism by which IL-6 controls IL-2 responsiveness in recently activated CD4^+^ T cells, we examined gene expression differences between OTII cells activated in sufficient and deficient IL-6 environment (**Table S1**). As expected, on day 3 after HDM^LPS^ sensitization OTII cells primed in *Il6*^−*/*−^ mice overexpressed IL-2 highly inducible genes (25), such as *Il2ra* and *Cish* (**Fig. 6A**). In contrast, one of the genes most significantly downregulated was *Socs3* (**Fig. 6A**). In addition, earlier identified genes regulated in *Socs3*-deficient livers (30) (**Fig. 6B**) and macrophages (31) (**Fig. 6C**) were highly enriched in OTII cells from *Il6*^−*/*−^ mice, suggesting dominant regulation by IL-6-induced SOCS3 in early T cell activation to HDM^LPS^.

**Fig 6.**
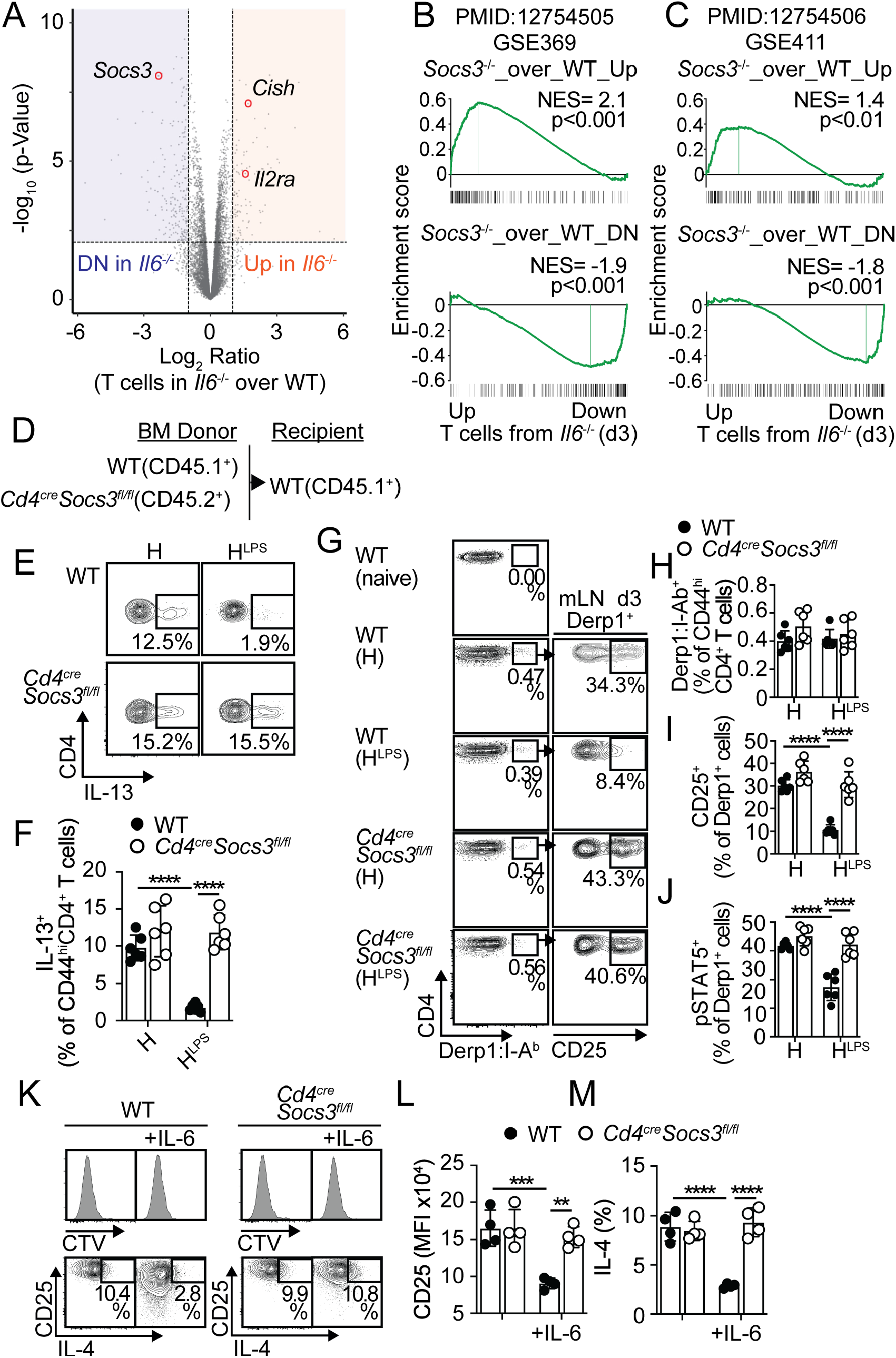
IL-6 suppresses allergen-specific Th2 cell responses by promoting SOCS3-mediated inhibition of IL-2 responsiveness. (**A-C**) WT and *Il6*^*-/-*^ mice were transferred with OTII cells and i.n sensitized with HDM^LPS^+OVA. On day 3, OTII cells were sorted from mLN and RNA-seq was performed (three replicates). Volcano plot highlighting the IL-2-driven genes *Il2ra* and *Cish*, and the IL-6-driven gene *Socs3*, upregulated and downregulated in OTII cells from *Il6*^*-/-*^ mice, respectively (FDR <0.05, −2 FC. See **Table S1**) (**A**). GSEA plots showing the enrichment of genes in OTII cells from *Il6*^*-/-*^ versus WT mice for genes regulated by IL-6-driven SOCS3 in liver cells (**B**) and macrophages (**C**). (**D-F**) WT:*CD4*^*cre*^*-Socs3*^*fl/fl*^ chimeras (**D**) were i.n sensitized with HDM or HDM^LPS^ and i.n challenged with HDM. Frequencies of IL-13^+^ cells within the WT and *CD4*^*cre*^*-Socs3*^*fl/fl*^ CD44^hi^CD4^+^ T cell compartments in the lungs (n=6) (**E-F**). (**G-J**) WT:*CD4*^*cre*^*-Socs3*^*fl/fl*^ chimeras (**D**) were i.n sensitized with HDM or HDM^LPS^ and analyzed on day 3. Frequencies of total (**G-H**), CD25^+^ (**G, I**), and pSTAT5^+^ (**J**) Derp1-specific CD44^hi^CD4^+^ T cells within the WT and *CD4*^*cre*^*-Socs3*^*fl/fl*^ CD44^hi^CD4^+^ T cell compartments in the mLN (n=6). (**K-M**) CTV-labeled WT (CD45.1^+^) and *CD4*^*cre*^*-Socs3*^*fl/fl*^ (CD45.2^+^) CD4^+^ T cells from naïve mice were mixed 1:1 and stimulated *in vitro* with plate-bound anti-CD3 and soluble anti-CD28 for 48h. 1ng/ml rIL-6 or PBS were added for additional 72h. CTV profiles and CD25 (**K-L**) and IL-4 expression in WT (**K, M**) and *CD4*^*cre*^*-Socs3*^*fl/fl*^ CD4^+^ T cells (values in quadruplicate). Data are representative of at least two independent experiments (mean±S.D., two-way Anova).

SOCS3 is strongly induced by IL-6 (32) and has been shown to prevent activation of STAT3 by IL-6 (30, 31, 33). However, SOCS3 can also inhibit signaling by many different cytokines (34), including IL-2 (35). To test whether SOCS3 influenced IL-2 responsiveness of CD4^+^ T cells and Th2 cell polarization to HDM, we first sensitized WT:*CD4*^*cre*^*-Socs3*^*fl/fl*^ mixed BM chimeras (**Fig. 6D**) with HDM or HDM^LPS^ and challenged them with HDM. We determined the frequency of IL-13^+^ Th2 cells within the WT and *Socs3*^−/−^ CD44^hi^CD4^+^ T cell compartments and found that whereas HDM^LPS^ sensitization prevented the accumulation of WT Th2 cells in the lungs, it failed to prevent the increase of *Socs3*^−/−^ Th2 cells (**Fig. 6E-F**). These results indicated that SOCS3 expression in responding T cells was intrinsically required to suppress the Th2 cell differentiation program to HDM. Since Th2 cell commitment to HDM required strong IL-2 signaling, we next tested whether SOCS3 expression influenced the IL-2 responsiveness of HDM-responsive T cells. Thus, we analyzed HDM/Derp1-specifc T cells (36) in the mLNs of WT:*CD4*^*cre*^*-Socs3*^*fl/fl*^ mixed BM chimeras on day 3 after HDM or HDM^LPS^ sensitization. The frequencies of Derp1-specific T cells within the WT and *Socs3*^−/−^ CD4^+^ T cell compartments were similar after HDM or HDM^LPS^ sensitization (**Fig. 6G-H**). As expected, CD25 expression and IL-2-driven pSTAT5 was downregulated in WT Derp1-specific T cells from HDM^LPS^-treated compared with HDM-treated mice (**Fig. 6G, I-J**). In contrast, *Socs3*-deficient Derp1-specific T cells maintained elevated expression of CD25 and IL-2-driven pSTAT5 in HDM^LPS^-treated mice (**Fig. 6G, I-J**). These results showed that SOCS3 expression in HDM-specific T cells is required to limit IL-2/STAT5 responsiveness shortly after T cell activation. Since this effect was also dependent on IL-6 signaling, our results further suggest that IL-6 controls SOCS3 expression in T cells to constraint IL-2 responsiveness and Th2 cell commitment. To confirm this, CD3/CD28-stimulated WT and *Socs3*-deficient CD4^+^ T cells were co-cultured in the presence of rIL-6. WT and *Socs3*-deficient CD4^+^ T cells proliferated at a similar rate, either in the presence or absence of IL-6 (**Fig. 6K**). As expected, IL-6 reduced the expression of CD25 and IL-4 in WT CD4^+^ T cells, but it failed to have similar effect in *Socs3*-deficient CD4^+^ T cells (**Fig. 6K-M**). Overall, our findings suggest that IL-6-driven SOCS3 acts to inhibit IL-2 responses in recently activated T cells and subsequent Th2 cell lineage commitment.

### JAK1 inhibition blocks IL-2 signaling and prevents Th2 bias in the absence of IL-6

JAK family members are key in initiating cytokine receptor signaling. IL-2 activates JAK1 and JAK3, with JAK1 playing a dominant role (37, 38). Thus, we tested whether treatment of *Il6*^*-*/-^ mice with the JAK1 selective inhibitor, Upadacitinib, blocked IL-2 signaling in recently activated T cells and hence their polarization to Th2 cells. Upadacitinib administered on days 1 and 2 post-sensitization (**Fig. 7A**) did not affect the expansion of the donor OTII cells (**Fig. 7C**) but induced downregulation of CD25 expression (**Fig. 7B** and **7D**) and IL-2-driven pSTAT5 (**Fig. 7B** and **7E**) in donor OTII cells from HDM^LPS^-treated *Il6*^*-*/-^ mice. Furthermore, this course of Upadacitinib treatment suppressed IL-4-GFP^+^ expression in donor OTII cells (**Fig. 7G**) and thus the accumulation of IL-4-GFP^+^4get.OTII cells (**Fig. 7I**) in HDM^LPS^-sensitized *Il6*^-/-^ recipients without affecting total OTII cell expansion (**Fig. 7H**). Nonetheless, a later Upadacitinib administration on days 3 and 4 post-sensitization did not affect IL-4-GFP^+^ expression in donor OTII cells (**Fig. 7F-I**). We further analyzed the effect of Upadacitinib treatment on lung effector Th2 cell responses developed in the absence of IL-6 signaling. Thus, we treated WT:*Lck*^*cre*^*-Il6r*^*fl/fl*^ mixed BM chimeras (**Fig. 7J**) with Upadacitinib at different times during sensitization (i.e., 1-2 vs. 3-4 days post-sensitization) (**Fig. 7K**). Mice were then challenged, and cytokine production by WT and *Il6r*^*-/-*^ T cells were analyzed in the lung. In HDM^LPS^-sensitized mice, treatment with Upadacitinib on days 1-2 prevented Th2 cell cytokine production by *Il6r*^*-/-*^ CD44^hi^CD4^+^ T cells, while treatment with Upadacitinib on days 3-4 had no effect on Th2 cell cytokine production by *Il6r*^*-/-*^ CD44^hi^CD4^+^ T cells (**Fig. 7L-M**). These data suggest that JAK1 inhibition in recently activated T cells can effectively suppress IL-2 signaling and subsequent Th2 cell lineage commitment.

**Fig. 7.**
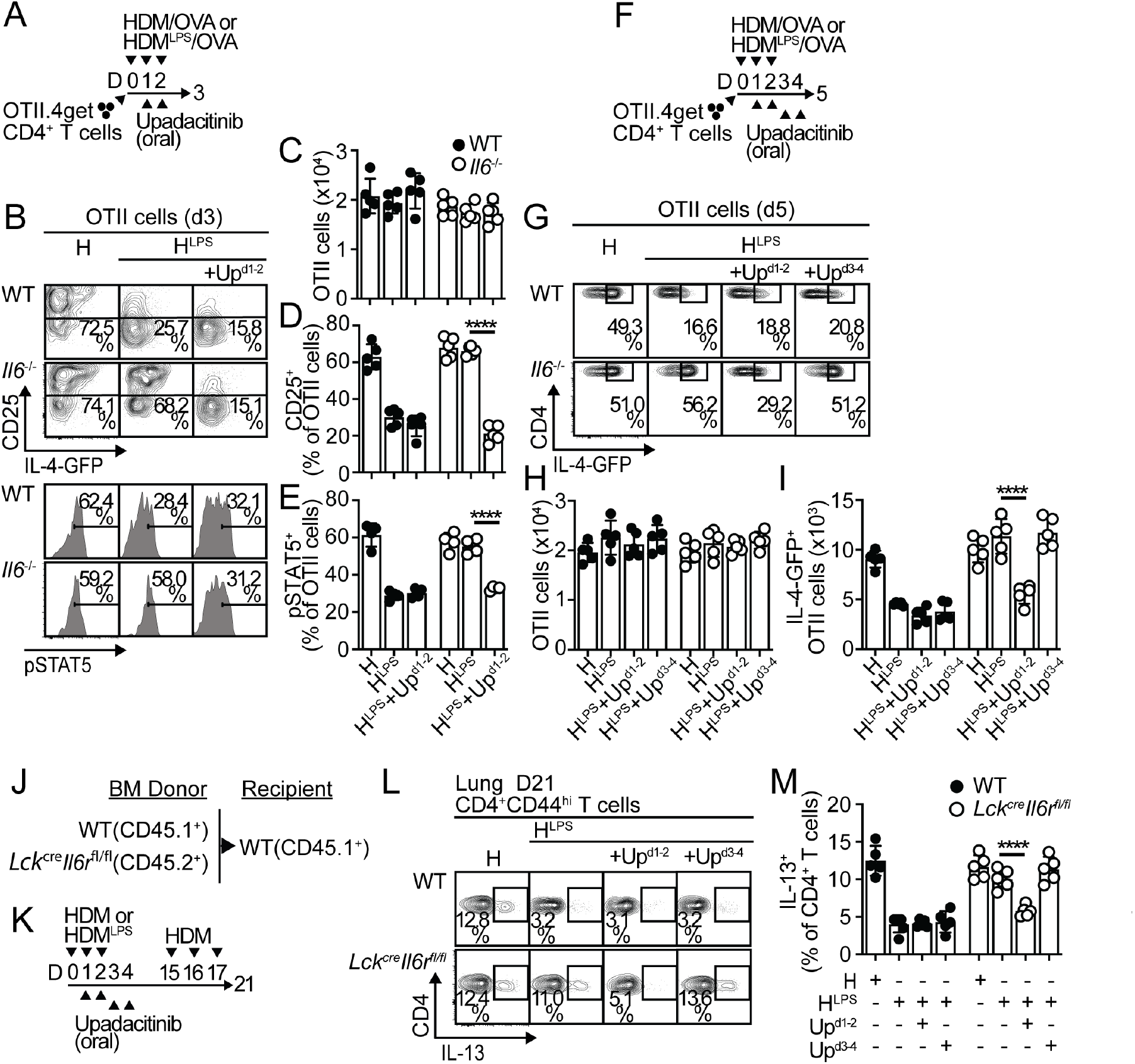
JAK1 inhibition prevents excess IL-2 signaling and Th2 cell polarization in the absence of IL-6 signaling. (**A-F**) WT and *Il6*^*-/-*^ mice were transferred with OTII.4get cells, i.n treated with HDM or HDM^LPS^ + OVA, orally treated with 20mg/Kg/day JAK1 inhibitor Upadacitinib or vehicle at the indicated time points and analyzed on day 3 (**A-E**) and day 5 (**F-I**). Frequencies and numbers of total (**C, H**), CD25^+^ (**B, D**), pSTAT5^+^ (**B, E**), and IL-4-GFP^+^ (**G, I**) OTII cells in the mLN. (**J-M**) WT:*Lck*^*cre*^*-Il6r*^*fl/fl*^ chimeras (**J**) were i.n sensitized with HDM or HDM^LPS^, orally treated with Upadacitinib or vehicle, and i.n challenged with HDM (**K**). Frequencies of IL-13^+^ cells within the WT and *Lck*^*cre*^*-Il6r*^*fl/fl*^ CD44^hi^CD4^+^ T cell compartments in the lungs (**L-M**). Data are representative of two independent experiments (mean±S.D., n=5, one-way Anova).

Collectively, our data indicate that IL-6 signaling in allergen-specific T cells is required to prevent bias towards the Th2 cell subset and subsequent allergic inflammation. Our work supports a model in which IL-6 upregulates SOCS3 expression, which limits IL-2 signaling in antigen-specific T cells and, thereby, restricts the acquisition of a Th2 cell differentiation program. Furthermore, we found that Upadacitinib, a JAK1 inhibitor, can suppress IL-2 signaling in recently activated T cells and their subsequent commitment to the Th2 cell lineage, suggesting that blocking the JAK1 pathway could be used to prevent new allergen sensitizations in environments with reduced IL-6 signaling.

## Discussion

Defective IL-6 signaling caused by mutations in the IL-6 receptor complex (5-7) or downstream mediators such as STAT3 (8-11) leads to pathological Th2-biased immune responses. Despite the negative role of aberrant Th2-biased responses in patients harboring these mutations, little is known about how Th2 immunity is favored under defective IL-6 signaling. Our data demonstrate that defective IL-6 signaling in CD4^+^ T cells responding to allergens enables prolonged IL-2 signaling that favors Th2 cell differentiation. Thus, we show that IL-6 is key in limiting IL-2 responsiveness in activated CD4^+^ T cells and that by restricting IL-2 signaling, IL-6 suppresses the Th2 cell differentiation program in these cells.

Our findings support a model in which IL-6 is produced in response to detection of pathogen-associated molecular patterns (PAMPs) contained in allergens. Several studies have indicated that high levels of exposure to microbial products, particularly bacterial endotoxins or LPS, inversely correlate with the development of allergen-induced, Th2-driven diseases such as allergic asthma and atopy (19-21). LPS detection by inflammatory monocyte-derived dendritic cells (mo-DCs) has been linked to suppression of Th2 cell immune responses to allergens (18). In response to LPS, mo-DCs produce cytokines, such as tumor necrosis factor alpha (TNFα), that instruct type-2 conventional dendritic cells (cDC2s) for IL-12 production (15, 18). As a consequence, CD4^+^ T cells interacting with cDC2s upregulate T-bet, which preclude Th2 cell differentiation (14, 15, 39, 40) and subsequent pathogenic allergic responses (15). However, our current data demonstrate that LPS detection can also prevent the development of allergic Th2 cell responses by an IL-12/T-bet-independent mechanism. Here, we found that LPS-driven IL-6 and subsequent signaling in allergen-activated CD4^+^ T cells contributed to preventing the Th2 cell differentiation program by restricting IL-2 signaling that would otherwise favor Th2 polarization. Therefore, IL-12/T-bet-dependent and IL-6-dependent pathways are independent mechanisms that work together to ensure the avoidance of abnormal Th2-biased immune reactions. Although both pathways could be thought of as having overlapping roles in preventing Th2 allergic responses, the lack of one or the other seems to lead to a lack of effective suppression of Th2 responses and the development of allergic diseases. In particular, a defective IL-12/T-bet-dependent pathway operating during infancy could be responsible for the higher susceptibility to Th2-bias and the increased tendency to develop allergic airway inflammation observed in children (15). On the other hand, as stated above, defective IL-6-dependent pathway also leads to pathological over-activation of Th2 immunity (5-11). Thus, it appears that both pathways must be active to ensure adequate prevention of Th2 cell responses. However, our data support that when LPS levels or IL-12 production are high, allowing for strong T-bet-dependent signals, the IL-6-dependent pathway is no longer necessary.

IL-2 is a well-known potent driver of T cell expansion (41). However, IL-2 has other immunomodulatory functions in antigen receptor-activated T cells. Particularly, IL-2 can promote Th1 (42) and Th2 (27-29) fate decisions in proliferating CD4^+^ T cells, while suppressing the differentiation of Th17 (42-44) and T follicular helper (Tfh) cells (45). Th2 cells depend on IL-4 signals and the transcriptional activities of STAT6 and GATA3 (46). Thus, Th2 cells depend on expression of the IL-4 receptor, which consists of IL4Rα and γ_c_. IL4Rα is not expressed by naïve CD4^+^ T cells and thus must be induced following antigen encounter to allow IL-4 responsiveness and potent Th2 cell differentiation. Studies indicate that IL-2 promotes Th2 fate decisions by inducing IL4Rα (29). In addition, IL-2 stabilizes the accessibility of the *Il4* locus (27-29), allowing for early production of IL-4 (27, 28) that serves as a positive feedback loop that preserves the Th2 cell phenotype. IL-2-driven induction of IL-4Rα and chromatin accessibility at the *Il4* locus required STAT5 (28, 29). Thus, supporting data indicate a key role for IL-2-driven STAT5 activation in the priming of Th2 cell differentiation and helping maintain this phenotype. On the other hand, our data support that IL-6 is a key negative regulator of Th2 cell differentiation by counteracting IL-2 signaling.

IL-2 binds with high affinity to a cell surface receptor complex consisting of IL-2Rα/CD25, IL-2Rβ/CD122 and γ_c_. IL-2 can also bind with low affinity to the IL-2Rβ/CD122 and γ_c_ dimer. IL-2 induces expression of IL-2Rα/CD25 (25) serving as a positive feedback loop. We found that IL-6 interrupted this positive feedback loop, causing cessation of IL-2Rα/CD25 induced expression, thus rendering activated CD4^+^ T cells unable to respond to IL-2 with high affinity. The tyrosine-protein kinases JAK1 and JAK3, which are associated with the IL-2Rβ/CD122 and γ_c_ subunits, respectively, are activated after IL-2 binding. This creates docking sites for STAT5 that facilitate STAT5 activation and subsequent nuclear translocation for transcription of target genes (47). SOCS family proteins are negative feedback inhibitors of cytokine-driven signaling that act through the JAK/STAT pathway (3). In particular, studies have identified SOCS1 (48) and SOCS3 (35) as inhibitors of IL-2 signaling, both of which function by inhibiting the kinase activity of JAK1. We demonstrate that SOCS3 is strongly induced by IL-6 in antigen receptor-activated CD4^+^ T cells and potently inhibits IL-2-induced STAT5 activation and Th2 cell differentiation to HDM allergens. Mimicking this mechanism, we have further shown that a selective JAK1 inhibitor can also inhibit IL-2-driven STAT5 activation and polarization toward a Th2 phenotype, particularly in IL-6-deficient environments that promote reduced SOCS3 activity in activated CD4^+^ T cells and presumably high JAK1 activation. JAK1 has also been implicated as a transducer in IL-4-signaling (49). Thus, JAK1 inhibition could directly inhibit Th2 differentiation by interfering with an IL-4-driven positive feedback loop. However, our data show that at doses that inhibited IL-2-driven STAT5 activation, the JAK1 inhibitor did not markedly alter the IL-4-signaling system. In brief, our data reveal a major role for JAK1 in IL-2-mediated signaling and Th2 bias that could have therapeutic implications.

Our results show that IL-6 interferes with high-affinity IL-2 signaling, which is critically required for Th2 fate commitment. However, IL-6 signaling can also inhibit the up-regulation of IL-2Rβ/CD122, thereby inhibiting low-affinity IL-2 signaling (50). This mechanism has been shown to control a state of persistent IL-2 hyporesponsiveness upon late and sustained antigen receptor triggering that enables the generation of Tfh cells in the germinal center (GC) (50). Without this mechanism of IL-6-driven inhibition of IL-2 signaling, Tfh cells receiving cognate interactions from GC B cells would initiate an IL-2 signaling cycle that would inhibit the maintenance of their Tfh phenotype. Our results, together with the aforementioned data, demonstrate that IL-6 can control IL-2 signaling by different mechanisms and at various T cell activation and differentiation stages, which determine particular immunomodulatory functions. However, since IL-2 signaling is primarily initiated by T-cell antigen receptor activation, it stands to reason that the role of IL-6 in counteracting IL-2 signaling is more prominent upon ongoing antigen presentation, such as at the onset of the immune response or in the GC. Conventional dendritic cells and B cells act as antigen-presenting cells in these situations, so future experiments should determine whether these cells are a source of IL-6 and if so, the specific pathways that determine IL-6 production.

Taken together, our data demonstrate that IL-6 signaling in allergen-specific T cells is essential for preventing Th2 development by counteracting IL-2-driven pro-Th2 signals. This action of IL-6 on allergen-responding T cells was mediated by upregulation of SOCS3 and could be mimicked by pharmacological inhibition of JAK1. Our data provide insights for understanding the immunological processes behind skewed Th2 responses in patients with defective IL-6 signaling and have the potential to open new therapeutic options for these patients.

## Materials and Methods

### Mouse strains

The mouse strains used in these experiments include: C57BL/6J (B6), B6.SJL-Ptprca Pepcb/BoyJ (CD45.1^+^ B6 congenic), C57BL/6-Tg(TcraTcrb)425Cbn/J (OTII), B6.129-Il4tm1Lky/J (B6.4get IL-4 reporter mice), B6.129S6-Tbx21tm1Glm/J (*Tbx21*^−*/*−^), B6.129S2-IL-6tm1Kopf/J (*Il6*^−*/*−^), B6.129S4-Il2ratm1Dw/J (*Cd25*^−*/*−^), B6.Cg-Tg(Lck-cre)3779Nik/J (*Lck-cre*), B6;SJL-Il6ratm1.1Drew/J (*Il6ra*^*fl/fl*^), B6N.129-Il21rtm1Kopf/J (*IL-21r*^−*/*−^), B6.Cg-Tg(Cd4-cre)1Cwi/BfluJ (*Cd4-cre*), and B6;129S4-Socs3tm1Ayos/J (*Socs3*^*fl/fl*^). B6.4get mice were originally obtained from Dr. M. Mohrs (Trudeau Institute). All other mice were originally obtained from Jackson Laboratory and were bred and housed in the University of Alabama at Birmingham animal facility under specific pathogen–free conditions. Experiments were equally performed with male and female mice. The University of Alabama at Birmingham Institutional Animal Care and Use Committee approved all procedures involving animals.

### Immunizations

HDM (Dermatophagoides pteronyssinus and D. farina) extract was obtained from Greer Laboratories (<30 EU/mg endotoxin). Endotoxin quantification was determined using the Pierce Chromogenic Endotoxin Quant Kit (A39552S; ThermoFisher Scientific). Mice were intranasally administered (i.n.) with 100μg of HDM extract, +/- 5μg of LPS-free EndoFit OVA (<0.1 EU/mg endotoxin; InvivoGen) +/- LPS from Escherichia coli 0111:B4 (Sigma-Aldrich) +/- 100 ng of rIL-6 (PeproTech) daily for 3 days and challenged (i.n.) with 100μg of HDM +/- 5μg of LPS-free EndoFit OVA for 3 days. i.n. administrations were given in 100μl of PBS. In some experiments, mice were intraperitoneally administered (i.p.) with 250 μg of a mix of anti–IL-6/R (15A7; Bio X Cell) and anti-IL-6 (MP5-20F3; Bio X Cell) neutralizing Abs, 250μg of anti-CD25 neutralizing Ab (PC-61.5.3; BioXCell), 250 μg of a mix of anti-IL-2 neutralizing Abs (JES6-1A12 and S4B6-1; Bio X Cell), or 60,000U rIL-2 (National Cancer Institute) at the indicated time points. In some experiments, mice were orally treated with JAK1 inhibitor Upadacitinib (ABT-494; Selleck Chemicals) at a dose of 20mg/Kg per day in palm oil at the indicated time points.

### BM chimeras

Recipient mice were irradiated with 950 Rads from a high-energy X-rays source delivered in a split dose and reconstituted with 10^7^ total BM cells. Mice were allowed to reconstitute for at least 8-12 weeks before HDM treatment.

### Cell preparation and flow cytometry

Lungs were isolated, cut into small fragments and digested for 45 min at 37°C with 0.6 mg/ml collagenase A (Sigma) and 30 μg/ml DNAse I (Sigma) in RPMI-1640 medium (GIBCO). Digested lungs, mLN or spleens were mechanically disrupted by passage through a wire mesh. Red blood cells were lysed with 150 mM NH4Cl, 10 mM KHCO3 and 0.1 mM EDTA. Fc receptors were blocked with anti-mouse CD16/32 (5 μg/ml; BioXCell), followed by staining with fluorochrome-conjugated Ab. Fluorochrome-labeled anti-B220 (RA3-6B2), anti-CD3 (17A2), anti-CD4 (GK1.5), anti-CD11b (M1/70), anti-CD11c (HL3), anti-CD25 (PC61), anti-CD44 (IM7), anti-CD45.1 (A20), anti-CD45.2 (104), anti-CD138 (281-2), anti-IgE (R35-72y) anti-Ly6C (AL-21), anti-Ly6G (IA8), and anti-Siglec-F (E50-2440) were from BD Biosciences. The I-A^b^ HDM Derp1_217-227_ MHC class II tetramer was obtained from the NIH Tetramer Core Facility. Dead cell exclusion was performed using 7-AAD (Calbiochem). For intracellular cytokine staining of T cells, cell suspensions were stimulated with PMA (20 ng/ml) plus Calcimycin (1μg/ml) in the presence of BD GolgiPlug for 5h. Restimulated cells were surface stained, fixed and permeabilized with BD Cytofix/Cytoperm Plus Kit and, stained with antibodies against IL-13 (13A; eBiosciences), IL-5 (TRFK5; BioLegend), IL-4 (11B11; BD Biosciences), IFNγ (XMG1.2; BD Biosciences), and IL-17 (TC11-18H10.1; BioLegend). For pSTAT5 staining of *ex vivo* CD4^+^ T cell populations, cell suspensions were stimulated with 1μg/ml rIL-2 for 15 min and then fixed and permeabilized with BD Cytofix/cytoperm buffer (BD) and the Phosflow Perm Buffer III (BD) followed by Transcription Factor Phospho Perm Wash Buffer (BD) and incubation with anti-pSTAT5 pY694 (47; BD Biosciences). For pSTAT5 staining of *in vitro* activated CD4^+^ T cell, cells were fixed, permeabilized, and stained without stimulation with rIL-2. Foxp3 intracellular staining was performed using the Mouse regulatory T cell staining kit (eBioscience) and Abs against Foxp3 (FJK-16s; ebioscience), Flow cytometry was performed on Attune NxT instrument.

### Cell purifications, sorting, cell transfers and *in vitro* cultures

CD4^+^ T cells were isolated by MACs (Miltenyi Biotec) from the spleens of naïve mice. Equivalent numbers (5 × 10^4^-5 × 10^5^) of naïve OTII cells were transferred (i.v.) into naïve congenic recipients. For some experiments, transferred donor OTII cells were sorted from recipient mice, after staining with fluorochrome-conjugated B220, CD4, CD44, and CD45.1. All sorting experiments were performed using a FACSAria (BD Biosciences) sorter in the University of Alabama at Birmingham Flow Cytometry core. Sorted cells were more than 98% pure as determined by flow cytometry. For *in vitro* cultures, CD4^+^ T cells were labeled for 10 min at 37°C with CellTrace™ CTV (Molecular Probes, ThermoFischer Scientific) and then activated with plate-bound anti-CD3 (clone 145-2C11; 1.5 μg/ml) and soluble anti-CD28 (clone 37.51; 1.5 μg/ml) Abs for 48h at 37°C in flat-bottom 96-well plates. The indicated concentration of anti-IL-2 neutralizing Ab (JES6-1A12 and S4B6-1; Bio X Cell) anti-CD25 neutralizing Ab (PC-61.5.3; Bio X Cell) and rIL-6 (PeproTech) were added for additional 72h. Complete medium included RPMI 1640 supplemented with sodium pyruvate, HEPES (pH range, 7.2 to 7.6), non-essential amino acids, penicillin, streptomycin, 2-mercaptoethanol and 10% heat-inactivated FBS (all from Gibco).

### BAL collection and measurement of cytokines

BAL was collected using 0.5-1 ml sterilized PBS per mouse. The BAL fluids were centrifuged at 5,000×g for 10 min and the supernatants were frozen at − 80°C. IL-6 measurement was performed using IL-6 Mouse Uncoated ELISA Kit (88-7064-22; Invitrogen).

### RNA-sequencing (RNA-seq)

#### Primary Analysis

Library preparation and RNA-seq was conducted through Genewiz. Libraries were sequenced using a 1 × 50–base pair single-end rapid run on the HiSeq 2500 platform. The quality of raw sequence fastq-formatted files was assessed using fastQC (http://www.bioinformatics.babraham.ac.uk/projects/fastqc). Sequences were trimmed using Trim Galore (version 0.4.4) with phred33 scores, paired-end reads and the Nextera adapter options (http://www.bioinformatics.babraham.ac.uk/projects/trim_galore). Trimmed sequences were aligned using STAR (version 2.5.2a) with mouse GRCm39 and default options (51). Aligned reads were counted with HTseq-count (version 0.6.1p1) set for unstranded and using the GRCm39 annotation file (52).

#### Downstream Analysis

The R package edgeR (53) was used to assess differential expression between groups and to generate gene-by-sample matrices. Volcano plot was generated using R. Comparison of our data to other published data sets was accomplished using Gene Set Enrichment Analysis (GSEA) (54). Upstream regulator analysis was performed using Ingenuity Pathway Analysis (IPA; Qiagen Digital Insights, Redwood City CA, USA) (55).

### Quantification and Statistical analysis

All plots and histograms were plotted in FlowJo v.9 and v.10 software (Treestar). GraphPad Prism (Version 9) was used for data analysis. The statistical significance of differences in mean values was determined using two-tailed Student’s t test or one-way/two-way ANOVA with post-hoc Turkey’s multiple comparison test. P values of less than 0.05 were considered statistically significant. \**P* < 0.05 \*\**P* < 0.01 \*\*\**P* < 0.001 \*\*\*\**P* < 0.0001.

## Data availability

Raw data fastq files, processed data count files and a counts per million table for RNA-Seq analyses reported in this paper have been deposited to GEO database under the accession code: GSE212158. This paper does not report original code

## Acknowledgements

We thank Becca Burnham, Uma Mudunuru and Thomas S Simpler for animal husbandry. The work was supported by UAB and NIH grants 2R01AI116584 to B.L., and R01AI150664 and R01AI162698 to A.B.T. The X-RAD 320 unit was purchased using a Research Facility Improvement Grant, 1 G20RR022807-01, from the NCRR, NIH. Support for the UAB flow cytometry core was provided by grants P30 AR048311 and P30 AI027767.

## Author information

### Contributions

B.L. designed the study and wrote the manuscript. H.B., E.M., B.L., C.L., and A.M.P. performed the experiments. E.M. and B.L. analyzed the data. A.F.R. and D.D.H. analyzed the RNAseq data. E.N.B. provided resources. A.B.T. contributed to data interpretation and discussion. All authors reviewed the manuscript before submission.

