## Supplementary figures S1-S5 for "IL-6 prevents Th2 cell polarization by promoting SOCS3-dependent suppression of IL-2 signaling"

### Supplementary Information

Fig. S1. IL-6 signaling during allergen sensitization oppositely regulates Th2 and Th17 cell responses to HDM.

Fig. S2. IL-21 signaling does not affect allergen-specific Th2 cell-mediated immunity.

Fig. S3. IL-6 signaling is not required to suppress Th2 cell differentiation in the presence of high LPS or IL-12.

Fig. S4. IL-6 signaling in responder T cells prevents prolonged IL-2 responsiveness.

Fig. S5. IL-2 signaling on allergen-specific T cells does not regulate polarization toward a Th17 profile.

Table S1. Raw data file (Excel)

Table S2. Raw data file (Excel)

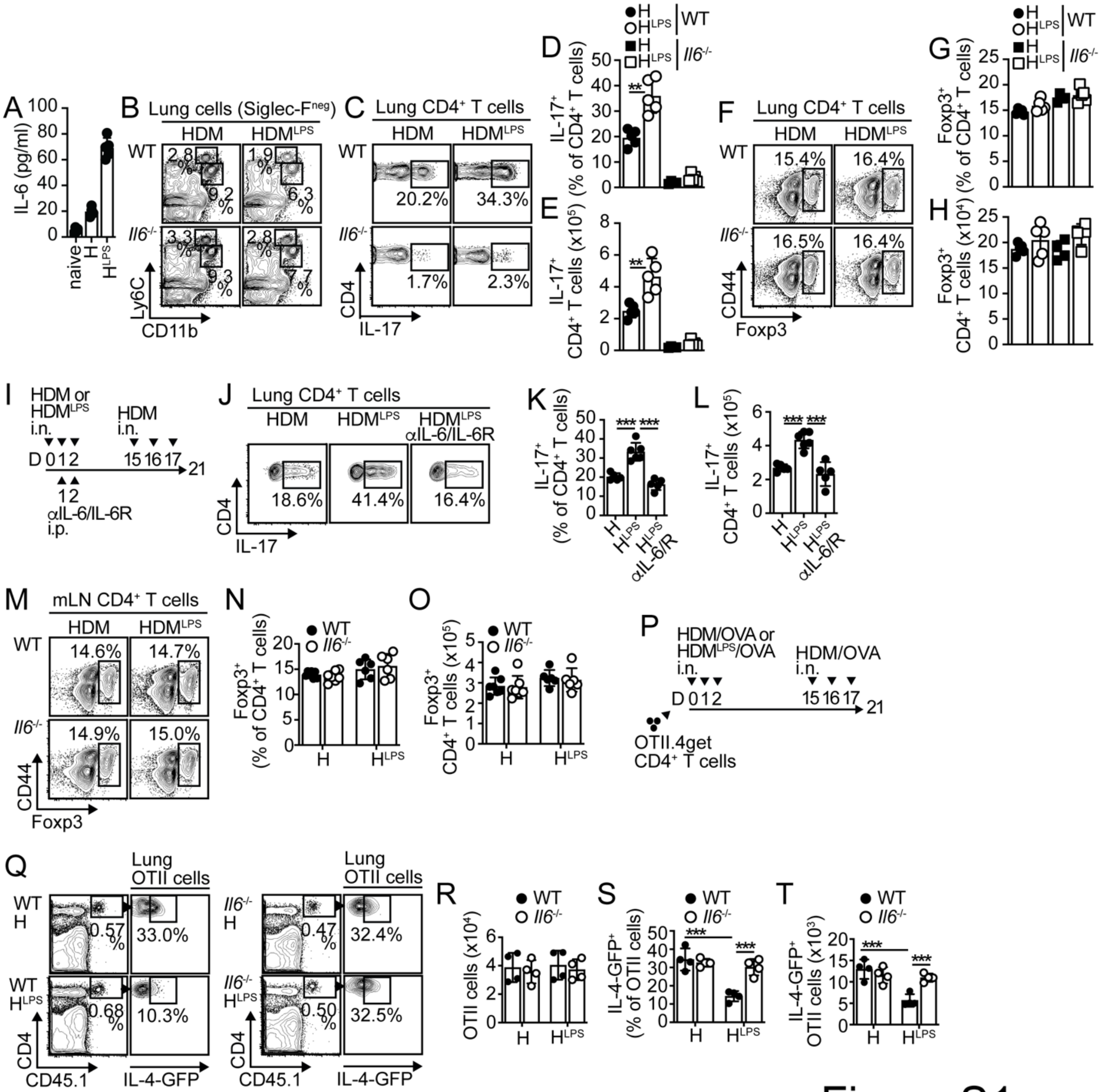


Figure S1

**Fig. S1. IL-6 signaling during allergen sensitization oppositely regulates Th2 and Th17 cell responses to HDM.**

(**A**) IL-6 in BAL from untreated or HDM- or HDM$^{LPS}$-treated B6 mice analyzed on day 1. (**B-H**) B6 (WT) and *Il6*$^{-/-}$ mice were i.n. sensitized with HDM or HDM$^{LPS}$ and challenged with HDM. Frequencies of $CD11b^{int}Ly6C^{hi}$ monocytes and $CD11b^{hi}Ly6C^{int}$ neutrophils in the lungs (**B**). Frequencies (**C-D**) and numbers (**E**) of IL-$17^+$ $CD4^+$ T cells in the lungs. Frequencies (**F-G**) and numbers (**H**) of $Foxp3^+$ $CD4^+$ T cells in the lungs. (**I-L**) B6 mice were i.n. sensitized with HDM or $HDM^{LPS}$. Some mice also received 250μg anti-IL-6 and anti-IL-6R (i.p.). On day 15, mice were i.n challenged with HDM and analyzed on day 21 (**I**). Frequencies (**J-K**) and numbers (**L**) of IL-$17^+$ $CD4^+$ T cells in the lungs. (**M-O**) Frequencies (**M-N**) and numbers (**O**) of $Foxp3^+$ $CD4^+$ T cells in the mLNs on day 5 after sensitization. (**P-T**) Mice were transferred with OTII.4get cells, i.n sensitized with HDM or $HDM^{LPS}$ + OVA, and challenged with HDM+OVA (**P**). Frequencies and numbers of total (**Q-R**) and IL-4-$GFP^+$ (**Q, S-T**) OTII cells in the lung. Data are representative of at least three independent experiments (mean±S.D., n=4-6, two-way and one-way Anova).

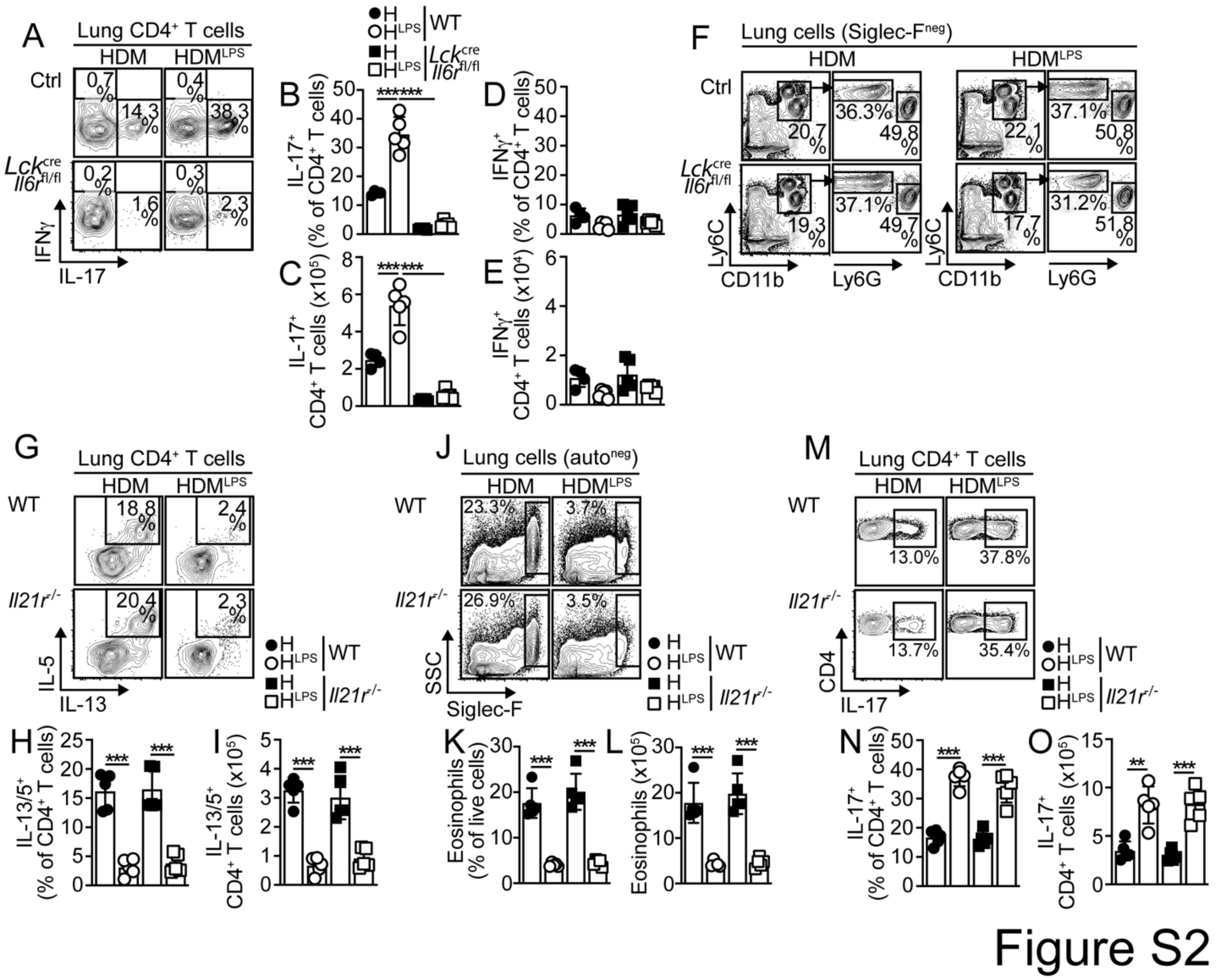

**Fig. S2. IL-21 signaling does not affect allergen-specific Th2 cell-mediated immunity.**

(**A-F**) *Lck^cre^-Il6r^fl/fl^* and control mice were i.n. sensitized with HDM or $HDM^{LPS}$ and challenged with HDM. Frequencies and numbers of $IL\text{-}17^{+}$ (**A-C**) and $IFN\gamma^{+}$ (**A, D-E**) $CD4^{+}$ T cells in the lung. Frequencies of $CD11b^{int}Ly6C^{hi}Ly6G^{lo}$ monocytes and $CD11b^{hi}Ly6C^{int}Ly6G^{hi}$ neutrophils in the lungs (**F**). (**G-O**) B6 (WT) and *Il21r^-/-^* mice were i.n. sensitized with HDM or $HDM^{LPS}$ and challenged with HDM. Frequencies (**G-H**) and numbers (**I**) of $IL\text{-}13^{+}$ $IL\text{-}5^{+}$ $CD4^{+}$ T cells in the lungs. Frequencies (**J-K**) and numbers (**L**) of eosinophils in the lungs. Frequencies (**M-N**) and numbers (**O**) of $IL\text{-}17^{+}$ $CD4^{+}$ T cells in the lungs. Data are representative of two independent experiments (mean±S.D., n=4-5, two-way Anova).

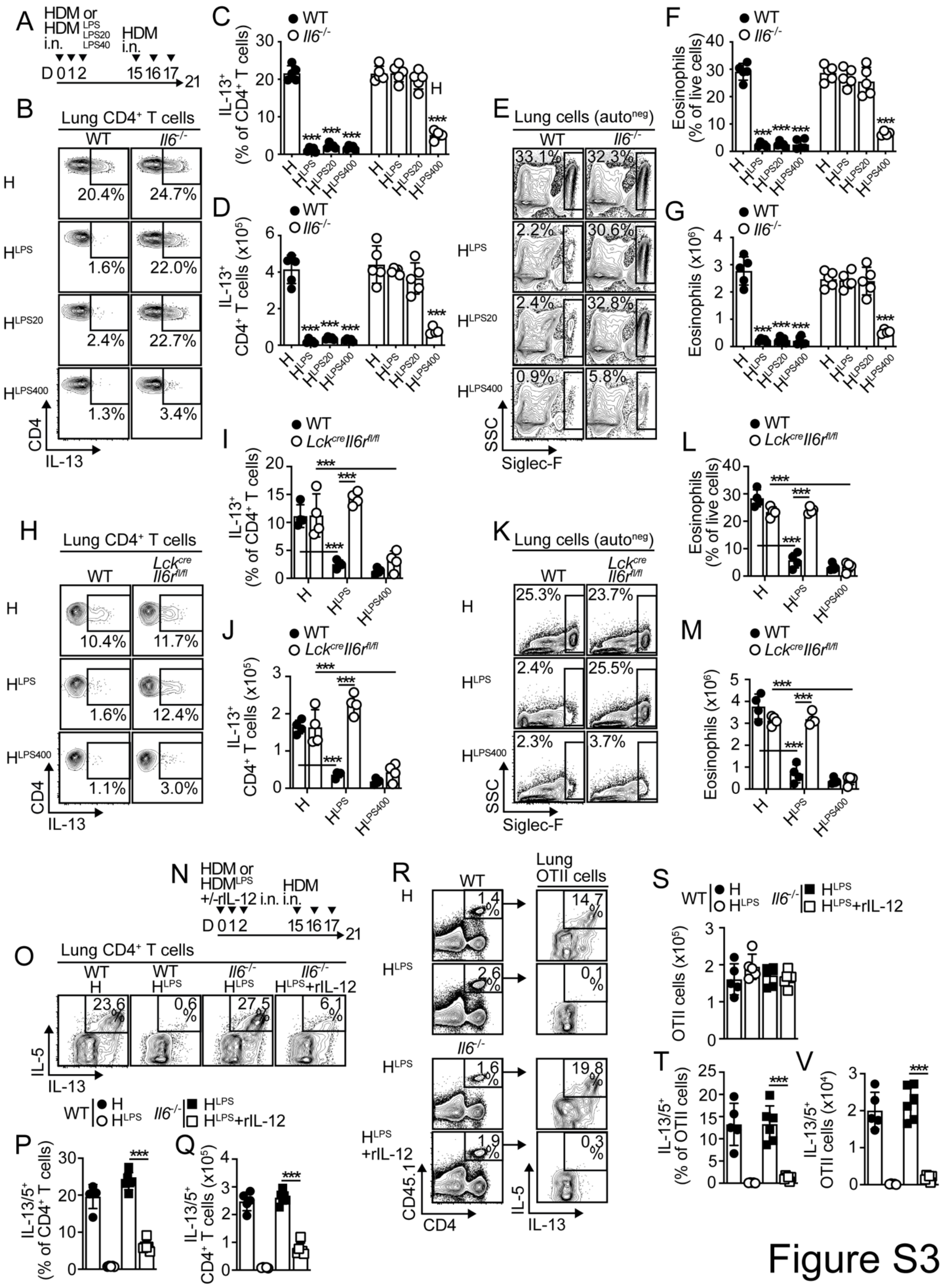

A
HDM or HDM LPS LPS20 LPS40 i.n.
HDM i.n.
D 0 1 2 15 16 17 21
B
Lung CD4+ T cells
WT
Il6-/-
H
20.4%
24.7%
HLPS
1.6%
22.0%
HLPS20
2.4%
22.7%
HLPS400
1.3%
3.4%
CD4
IL-13
C
WT
Il6-/-
IL-13+ (% of CD4+ T cells)
D
IL-13+ CD4+ T cells (x10^5)
E
Lung cells (autoneg)
33.1%
32.3%
2.2%
30.6%
2.4%
32.8%
0.9%
5.8%
SSC
Siglec-F
F
Eosinophils (% of live cells)
G
Eosinophils (x10^6)
H
Lung CD4+ T cells
WT
Lckcre Il6rfl/fl
10.4%
11.7%
1.6%
12.4%
1.1%
3.0%
I
J
K
Lung cells (autoneg)
25.3%
23.7%
2.4%
25.5%
2.3%
3.7%
L
M
N
HDM or HDMLPS +/-rIL-12 i.n.
HDM i.n.
D 0 1 2 15 16 17 21
O
Lung CD4+ T cells
WT H 23.6%
WT HLPS 0.6%
Il6-/- HLPS 27.5%
Il6-/- HLPS+rIL-12 6.1%
IL-5
IL-13
P
IL-13/5+ (% of CD4+ T cells)
Q
IL-13/5+ CD4+ T cells (x10^5)
R
Lung OTII cells
1.4%
14.7%
2.6%
0.1%
1.6%
19.8%
1.9%
0.3%
HLPS +rIL-12
CD45.1
CD4
S
OTII cells (x10^5)
T
IL-13/5+ (% of OTII cells)
V
IL-13/5+ OTII cells (x10^4)


Figure S3

**Fig. S3. IL-6 signaling is not required to suppress Th2 cell differentiation in the presence of high LPS or IL-12.**

(**A-G**) WT and *Il6*$^{-/-}$ mice were i.n sensitized with 100μg HDM containing different amount of LPS (HDM$^{LPS}$: 1μg LPS/mg, HDM$^{LPS20}$: 20μg LPS/mg, HDM$^{LPS400}$: 400μg LPS/mg). Mice were then challenged with HDM (**A**). Frequencies (**B-C**) and numbers (**D**) of IL-13$^{+}$ CD4$^{+}$ T cells in the lungs. Frequencies (**E-F**) and numbers (**G**) of eosinophils in the lungs. (**H-M**) *Lck*$^{cre}$*-Il6r*$^{fl/fl}$ and control mice were i.n. sensitized with containing different amount of LPS (HDM$^{LPS}$: 1μg LPS/mg, HDM$^{LPS400}$: 400μg LPS/mg) and challenged with HDM (**A**). Frequencies (**H-I**) and numbers (**J**) of IL-13$^{+}$ CD4$^{+}$ T cells in the lungs. Frequencies (**K-L**) and numbers (**M**) of eosinophils in the lungs. (**N-Q**) WT and *Il6*$^{-/-}$ mice were i.n sensitized with HDM or HDM$^{LPS}$+/-150ng rIL-12 and challenged with HDM (**N**). Frequencies (**O-P**) and numbers (**Q**) of IL-13$^{+}$ IL-5$^{+}$ CD4$^{+}$ T cells in the lungs. (**R-V**) WT and *Il6*$^{-/-}$ mice were transferred with OTII.4get cells, i.n sensitized with HDM+OVA or HDM$^{LPS}$+OVA+/-150ng rIL-12, and i.n. challenged with HDM+OVA. Frequencies and numbers of total (**R-S**) and IL-13$^{+}$ IL-5$^{+}$ (**R, T-V**) OTII cells in the lung. Data are representative of two independent experiments (mean±S.D., n=4-5, one-way Anova).

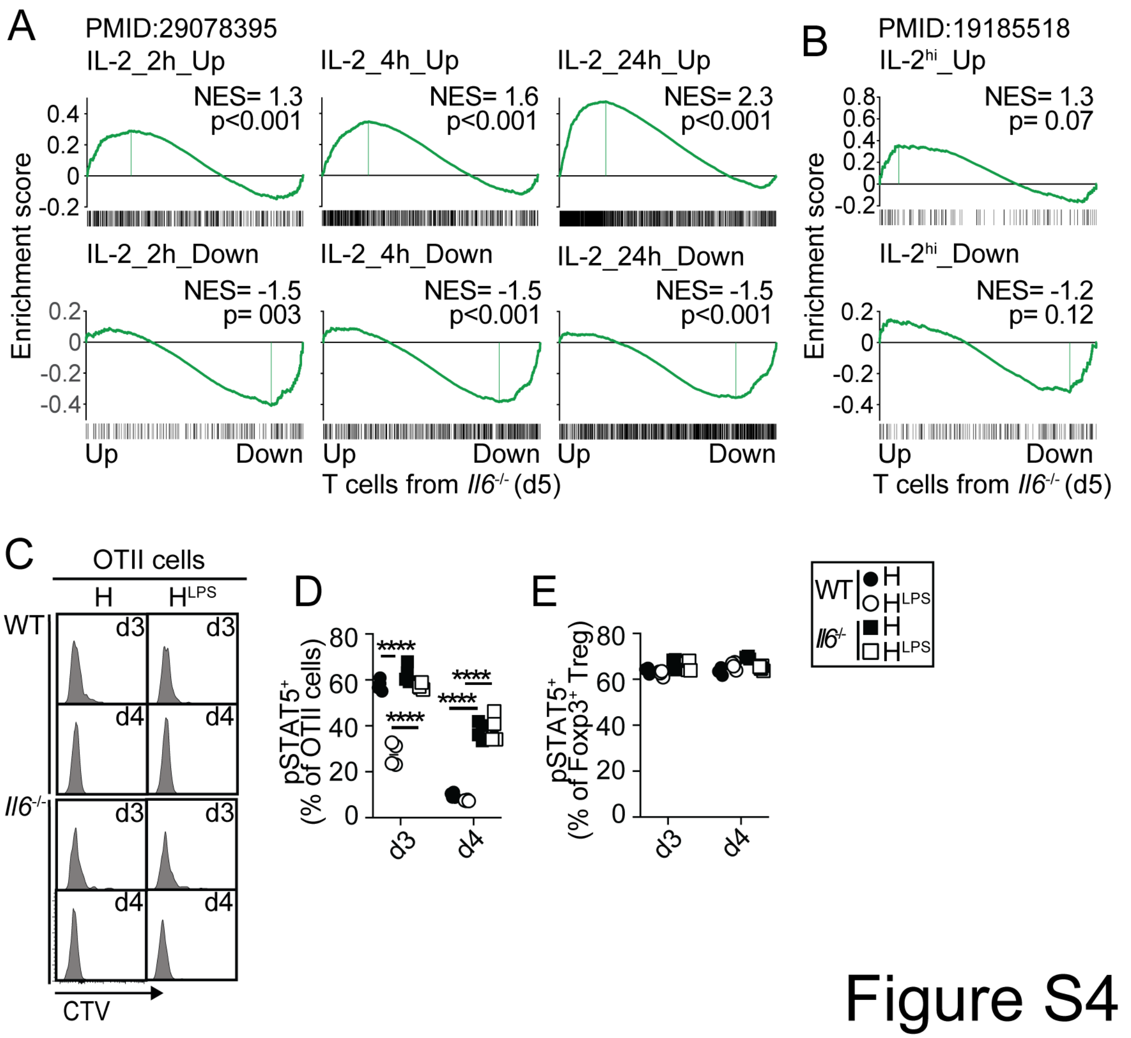


Figure S4

**Fig. S4. IL-6 signaling in responder T cells prevents prolonged IL-2 responsiveness.**

(**A-B**) WT and *Il6*$^{-/-}$ mice were transferred with OTII cells and i.n sensitized with HDM$^{LPS}$+OVA. On day 5, OTII cells were sorted from mLN and RNA-seq was performed (three replicates). 156 differentially expressed genes, with 43 up- and 113 down-regulated in OTII from *Il6*$^{-/-}$ mice, were identified (FDR <0.05, ≥2 FC. See **Table S1**). GSEA plots showing the enrichment of genes in OTII cells from *Il6*$^{-/-}$ versus WT mice for genes regulated by IL-2 (**A**) or by strong IL-2 signaling (**B**). (**C-E**) WT and *Il6*$^{-/-}$ mice were transferred with CTV-labeled OTII.4get cells and i.n sensitized with HDM or HDM$^{LPS}$ +

OVA. CTV profiles in donor OTII cells from mLNs (**C**). Cells from mLNs were stimulated with 1ug/ml rIL-2 for 15 min and STAT5 phosphorylation in OTII (**D**) and Foxp3$^{+}$ CD4$^{+}$ T cells (**E**) was determined. Data are representative of two independent experiments (mean±S.D., n=3-4, two-way Anova).

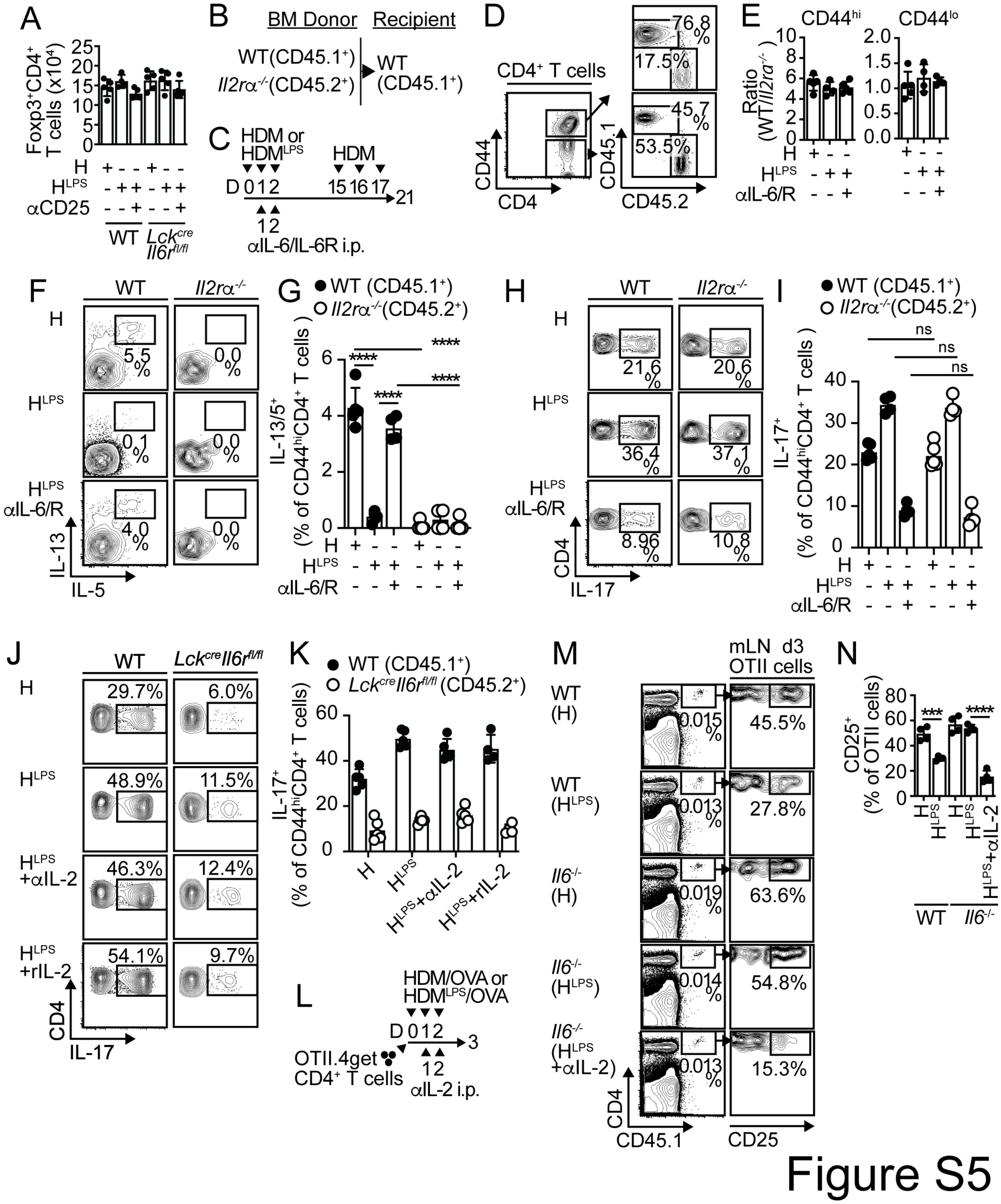


**Fig. S5. IL-2 signaling on allergen-specific T cells does not regulate polarization toward a Th17 profile.**

(**A**) WT:*Lck*$^{cre}$-*Il6r*$^{fl/fl}$ chimeras were i.n sensitized with HDM or HDM$^{LPS}$, i.p treated with anti-CD25 or PBS, and i.n challenged with HDM. Numbers of Foxp3$^{+}$CD4$^{+}$ T cells in the lung. (**B-I**) Irradiated B6 (CD45.1$^{+}$) mice were reconstituted with 1:1 BM mix of B6 (CD45.1$^{+}$) and *Il2r*$\alpha^{-/-}$ (CD45.2$^{+}$) donors (**B**). Chimeras were i.n sensitized with HDM or HDM$^{LPS}$. Some mice also received 250µg anti-IL-6 and anti-IL-6R (i.p.). Mice were then challenged with HDM (**C**). Frequencies of CD45.1$^{+}$ and CD45.2$^{+}$ cells in CD44$^{hi}$ and CD44$^{lo}$ CD4$^{+}$ T cells from the lungs (**D**). Ratio of WT to *Il2r*$\alpha^{-/-}$ CD44$^{hi}$ and CD44$^{lo}$ CD4$^{+}$ T cells (**E**). Frequencies of IL-13$^{+}$ IL-5$^{+}$ (**F-G**) and IL-17$^{+}$ (**H-I**) cells within the WT and *Il2r*$\alpha^{-/-}$ CD44$^{hi}$CD4$^{+}$ T cell compartments in the lungs. (**J-K**) WT:*Lck*$^{cre}$-*Il6r*$^{fl/fl}$ chimeras were i.n sensitized with HDM or HDM$^{LPS}$, i.p treated with anti-IL-2 Abs (JES6-1A12 and S4B6; 250µg each), rIL-2 (60,000U) or PBS, and i.n challenged with HDM (**L**). Frequencies of IL-17$^{+}$ cells within the WT and *Lck*$^{cre}$-*Il6r*$^{fl/fl}$ CD44$^{hi}$CD4$^{+}$ T cell compartments in the lungs. (**L-N**) WT and *Il6*$^{-/-}$ mice were transferred with OTII.4get cells, i.n treated with HDM or HDM$^{LPS}$ + OVA, and i.p treated with anti-IL-2 Abs or PBS (**L**). Frequencies of total (**M**) and CD25$^{+}$ (**M-N**) OTII cells in the mLN on day 3. Data are representative of two independent experiments (mean±S.D., n=4-5, one-way Anova).
