## Supplementary material for "IL-6 prevents Th2 cell polarization by promoting SOCS3-dependent suppression of IL-2 signaling": Table S1

### DAY 3

| | gene | length | logFC | logCPM | F | PValue | FDR |
|---|---|---|---|---|---|---|---|
| **1574** | Cd163l1 | 4508 | -5.661271 | 0.07571365 | 31.1579518 | 6.18E-05 | 0.00179419 |
| **4996** | Klhl4 | 3884 | -4.7712577 | 0.83721086 | 15.9742835 | 0.00126676 | 0.01405047 |
| **437** | Adam12 | 7675 | -4.4645053 | 0.59733124 | 44.6902383 | 9.16E-06 | 0.00055157 |
| **4630** | Il17re | 3592 | -4.1845345 | 0.1107445 | 80.0594708 | 3.01E-07 | 7.53E-05 |
| **2087** | Cpeb1 | 8516 | -4.1540581 | 0.67860764 | 48.1702157 | 6.03E-06 | 0.00043241 |
| **1081** | B4galnt4 | 3682 | -4.0388041 | 1.20945191 | 36.1613833 | 2.87E-05 | 0.00111882 |
| **4635** | Il1r1 | 6138 | -3.3794005 | 4.20953397 | 75.7364843 | 4.23E-07 | 8.51E-05 |
| **8446** | Serpinb1a | 2428 | -3.0922393 | 0.35192038 | 34.3112402 | 3.78E-05 | 0.00133735 |
| **4779** | Itga7 | 7267 | -3.0444059 | 2.0906744 | 88.6111135 | 1.61E-07 | 4.52E-05 |
| **6470** | Ntng2 | 1924 | -2.8780591 | 0.55474277 | 27.1695549 | 0.00012167 | 0.00278975 |
| **10457** | Usp35 | 9685 | -2.8354766 | 0.94374408 | 19.366655 | 0.00057155 | 0.00816559 |
| **4641** | Il21 | 3178 | -2.8274679 | 4.68586637 | 38.0248745 | 2.20E-05 | 0.00094611 |
| **9546** | Tigit | 2137 | -2.8100842 | 5.48009788 | 31.5690386 | 5.79E-05 | 0.00174489 |
| **4958** | Kit | 5182 | -2.7610966 | 1.37422851 | 22.5120502 | 0.00029394 | 0.00526943 |
| **4660** | Iltifb | 1111 | -2.680275 | 0.06521613 | 28.4983102 | 9.64E-05 | 0.00235238 |
| **6136** | Nav2 | 114472 | -2.6144804 | 3.59616095 | 87.7597412 | 1.71E-07 | 4.69E-05 |
| **4234** | H2-Q2 | 2032 | -2.5493704 | 5.49332925 | 43.3681617 | 1.08E-05 | 0.000621 |
| **8201** | Rsad2 | 3785 | -2.4936196 | 1.83718815 | 27.6869687 | 0.00011103 | 0.00262595 |
| **6640** | P4ha2 | 2345 | -2.4722275 | -0.0569993 | 17.4478042 | 0.00088654 | 0.01098702 |
| **4114** | Gprc5b | 4525 | -2.4385363 | 0.4519597 | 10.2429382 | 0.00624984 | 0.04067094 |
| **4572** | Ifit1bl1 | 3186 | -2.3746672 | 2.24547322 | 27.0855427 | 0.00012351 | 0.00280895 |
| **8966** | Socs3 | 2732 | -2.347845 | 3.78761751 | 141.313748 | 8.10E-09 | 1.13E-05 |
| **10940** | Zfp365 | 4257 | -2.3173661 | 1.86334482 | 34.5530108 | 3.64E-05 | 0.00131012 |
| **1079** | B430306N03Rik | 4390 | -2.3127904 | 0.97366393 | 12.7080317 | 0.00300418 | 0.02516445 |
| **9490** | Tgfb3 | 3385 | -2.2585414 | 2.55909741 | 52.9921838 | 3.51E-06 | 0.00031591 |
| **6187** | Ndrg4 | 3374 | -2.2179349 | 0.6868044 | 20.5443477 | 0.00044228 | 0.00685843 |
| **9503** | Tha1 | 1840 | -2.2151195 | 3.46858898 | 75.7263769 | 4.23E-07 | 8.51E-05 |
| **9857** | Tox2 | 4623 | -2.213407 | 6.27037331 | 145.51437 | 6.69E-09 | 1.13E-05 |
| **6469** | Ntn1 | 5738 | -2.1937084 | 0.35336094 | 19.465718 | 0.00055915 | 0.00802921 |
| **4636** | Il1r2 | 2688 | -2.1743208 | 1.80012045 | 25.2642053 | 0.00017225 | 0.00360942 |
| **8042** | Rorc | 3564 | -2.169213 | 2.93368458 | 78.9032765 | 3.29E-07 | 7.72E-05 |
| **7958** | Rnase4 | 1996 | -2.1667137 | 1.6081197 | 48.3146208 | 5.93E-06 | 0.00042791 |
| **3712** | Gm30603 | 624 | -2.1421475 | 2.0959981 | 15.761176 | 0.00133594 | 0.01447543 |
| **3123** | Fam20a | 3505 | -2.1398229 | 1.83478414 | 28.5425038 | 9.57E-05 | 0.00235182 |
| **2310** | D930020B18Rik | 4453 | -2.1150197 | 0.57019626 | 11.3762993 | 0.00441887 | 0.03254868 |
| **3400** | Gabrr2 | 3437 | -2.1126658 | 2.13832447 | 37.2027232 | 2.47E-05 | 0.00102144 |
| **2905** | Emp1 | 3231 | -2.0883472 | 2.37858734 | 40.2776325 | 1.62E-05 | 0.00079855 |

| | | | | | | | |
|---|---|---|---|---|---|---|---|
| **4951** | Kif5c | 6868 | -2.0734254 | 0.82723066 | 23.8649309 | 0.00022489 | 0.0044032 |
| **4097** | Gpr146 | 7193 | -2.065553 | 3.52553162 | 45.0626152 | 8.75E-06 | 0.00053832 |
| **2942** | Epdr1 | 2387 | -2.0526742 | 0.7596498 | 9.5117908 | 0.00789408 | 0.04772907 |
| **3928** | Gm46941 | 398 | -2.0472368 | 0.5406836 | 11.7182699 | 0.00399357 | 0.03043979 |
| **3649** | Gm17455 | 1192 | -2.0168086 | 0.21481603 | 13.4939876 | 0.00241597 | 0.02156941 |
| **8263** | S100a4 | 597 | -2.009455 | 2.86703937 | 22.9614038 | 0.00026862 | 0.0049904 |
| **8256** | Ryr1 | 15442 | -1.9986048 | 0.44807634 | 21.8941364 | 0.00033334 | 0.0057293 |
| **7734** | Rasgef1a | 3582 | -1.9879294 | 1.73697407 | 19.3446487 | 0.00057434 | 0.00817443 |
| **3714** | Gm30712 | 4505 | -1.8654384 | 1.56375953 | 14.7945102 | 0.00170925 | 0.01702902 |
| **5447** | Man1c1 | 6382 | -1.8575415 | 0.27084489 | 10.6142495 | 0.0055678 | 0.03762441 |
| **3059** | F2rl1 | 2768 | -1.8386265 | 0.95918865 | 12.7787254 | 0.00294503 | 0.02487585 |
| **411** | Acss1 | 3613 | -1.8153154 | 4.26446686 | 75.3573639 | 4.36E-07 | 8.62E-05 |
| **8262** | S100a3 | 3206 | -1.80095 | 1.1215641 | 22.5991589 | 0.00028883 | 0.00524024 |
| **4002** | Gmfg-ps | 576 | -1.7996136 | 0.83783684 | 21.9109406 | 0.00033219 | 0.00571829 |
| **975** | Atp1a3 | 4092 | -1.7922975 | 0.66857656 | 24.3648794 | 0.00020422 | 0.00411379 |
| **6547** | Oasl2 | 3136 | -1.7913214 | 0.53006774 | 21.6527031 | 0.00035034 | 0.00593102 |
| **3573** | Gm11346 | 1116 | -1.7865478 | 3.79060377 | 55.1359082 | 2.79E-06 | 0.0002733 |
| **6589** | Orm3 | 2491 | -1.7754552 | 0.91250634 | 11.26562 | 0.00456749 | 0.03317468 |
| **1316** | C030034L19Rik | 5830 | -1.7717801 | 1.99935472 | 33.017252 | 4.61E-05 | 0.00148985 |
| **750** | Apold1 | 3284 | -1.7711508 | 0.74636318 | 19.0313858 | 0.0006159 | 0.00857454 |
| **9284** | Syt13 | 3751 | -1.7671274 | 0.26218411 | 16.9426178 | 0.00099988 | 0.01189917 |
| **9794** | Tnfrsf25 | 2541 | -1.7469706 | 5.17903892 | 129.281166 | 1.45E-08 | 1.13E-05 |
| **3914** | Gm46412 | 10914 | -1.7258023 | 1.74782835 | 14.2365298 | 0.00197863 | 0.01889631 |
| **8230** | Rtp4 | 2945 | -1.7219764 | 4.77103873 | 73.1530491 | 5.23E-07 | 0.00010142 |
| **9280** | Syp | 2482 | -1.711474 | 2.68458809 | 26.2466764 | 0.00014368 | 0.00316541 |
| **430** | Acvrl1 | 3788 | -1.7091907 | 3.29495343 | 24.5679433 | 0.00019645 | 0.00399928 |
| **8264** | S100a6 | 719 | -1.6976595 | 5.52582857 | 19.4183431 | 0.00056504 | 0.00809315 |
| **4872** | Kcna2 | 14395 | -1.6786723 | 3.15002883 | 77.8791434 | 3.57E-07 | 8.03E-05 |
| **2311** | D930028M14Rik | 1074 | -1.6742323 | 0.90449301 | 13.2817215 | 0.00256068 | 0.02252196 |
| **4574** | Ifit3 | 2678 | -1.6734759 | 1.84258041 | 11.8425661 | 0.00385077 | 0.02975426 |
| **8786** | Slfn5 | 6354 | -1.6674741 | 5.77590881 | 154.699993 | 4.47E-09 | 1.13E-05 |
| **4882** | Kcnmb3 | 8336 | -1.6605445 | 2.06866189 | 17.541145 | 0.00086724 | 0.01082418 |
| **7213** | Pou6f1 | 5353 | -1.6564623 | 1.97896887 | 20.0649472 | 0.00049039 | 0.00737085 |
| **3924** | Gm46781 | 2619 | -1.646769 | 1.33897876 | 17.2246457 | 0.00093469 | 0.01140528 |
| **759** | Aqp3 | 1763 | -1.638721 | 4.00874984 | 48.4594372 | 5.83E-06 | 0.00042348 |
| **8675** | Slc28a2b | 4043 | -1.6327692 | 5.70815583 | 34.9053545 | 3.46E-05 | 0.00126711 |
| **8744** | Slc43a1 | 3880 | -1.6149955 | 1.65743486 | 24.0871434 | 0.00021542 | 0.00426975 |
| **3953** | Gm51710 | 5981 | -1.6122547 | 1.19438738 | 23.5573819 | 0.0002388 | 0.00456427 |
| **10183** | Ttc39c | 5915 | -1.6014378 | 4.07387053 | 29.5521614 | 8.06E-05 | 0.00209035 |
| **8428** | Septin10 | 5152 | -1.5948776 | 0.5062806 | 18.7035194 | 0.00066312 | 0.00896209 |

| | | | | | | | |
|---|---|---|---|---|---|---|---|
| **6545** | Oas2 | 4293 | -1.5932931 | 0.54587375 | 21.3635203 | 0.00037203 | 0.00615765 |
| **3343** | Frat1 | 2614 | -1.5833934 | 2.59388758 | 14.7507928 | 0.00172877 | 0.01716273 |
| **4571** | Ifit1 | 2638 | -1.5814909 | 3.47887209 | 16.8913465 | 0.00101228 | 0.01202134 |
| **4972** | Klf8 | 8902 | -1.577395 | 0.79647138 | 11.9949128 | 0.00368367 | 0.02883917 |
| **3618** | Gm15411 | 18344 | -1.5720971 | 1.09349448 | 14.0713359 | 0.00206748 | 0.01951023 |
| **8040** | Ropn1l | 910 | -1.5683304 | 0.99173903 | 18.8988533 | 0.00063451 | 0.00866956 |
| **4563** | Ifi213 | 8794 | -1.5596245 | 4.10149281 | 41.3481648 | 1.40E-05 | 0.00075785 |
| **9459** | Tesc | 1036 | -1.5527255 | 6.11693926 | 100.655769 | 7.22E-08 | 2.46E-05 |
| **4753** | Irf7 | 1934 | -1.5442847 | 6.08312284 | 178.629692 | 1.72E-09 | 7.07E-06 |
| **5283** | Lncpint | 1808 | -1.5367064 | 1.81949572 | 18.1344046 | 0.00075526 | 0.00975084 |
| **9053** | Sqor | 2365 | -1.5235657 | 3.73389036 | 37.8325015 | 2.26E-05 | 0.00096098 |
| **243** | A430088P11Rik | 4502 | -1.5145397 | 2.72939122 | 31.463906 | 5.89E-05 | 0.00175771 |
| **3396** | Gabbr1 | 5848 | -1.5067645 | 2.9820105 | 38.1177336 | 2.17E-05 | 0.00094109 |
| **9793** | Tnfrsf23 | 4111 | -1.4972232 | 0.58815989 | 9.79581658 | 0.00720236 | 0.04479788 |
| **7010** | Pipox | 1775 | -1.4949353 | 1.77920151 | 11.6520773 | 0.00407209 | 0.03080883 |
| **10533** | Vipr1 | 4902 | -1.4913381 | 3.40743862 | 23.2256808 | 0.0002549 | 0.0047828 |
| **6037** | Mx1 | 2697 | -1.4765109 | 1.41431889 | 14.1855806 | 0.00200556 | 0.01905362 |
| **2244** | Cxcr5 | 2638 | -1.4579052 | 7.33192602 | 77.4820148 | 3.68E-07 | 8.13E-05 |
| **5099** | LOC118567527 | 5304 | -1.4402131 | 4.66308471 | 56.1205252 | 2.52E-06 | 0.00025557 |
| **5153** | LOC118568610 | 15356 | -1.4349359 | 1.4678403 | 9.98648053 | 0.0067771 | 0.0429115 |
| **250** | A630023P12Rik | 646 | -1.433737 | 4.47218157 | 35.6477617 | 3.10E-05 | 0.0011735 |
| **1965** | Cnga1 | 2818 | -1.4320455 | 1.23911948 | 13.6664912 | 0.00230525 | 0.02092944 |
| **7713** | Rap1gap2 | 9251 | -1.4186978 | 2.5957694 | 20.5856443 | 0.0004384 | 0.00681696 |
| **155** | 4930417O13Rik | 1093 | -1.4177669 | 2.14627812 | 15.3823751 | 0.00146984 | 0.01532175 |
| **868** | Armcx6 | 2863 | -1.4144557 | 2.29814362 | 30.8883896 | 6.46E-05 | 0.00183667 |
| **4463** | Hpgds | 4019 | -1.4109338 | 2.58877522 | 30.7609124 | 6.60E-05 | 0.0018565 |
| **459** | Adcy6 | 13136 | -1.4046924 | 1.4294861 | 18.9186661 | 0.00063169 | 0.00866956 |
| **9250** | Susd1 | 3964 | -1.4030303 | 1.79068874 | 29.1863492 | 8.57E-05 | 0.0021832 |
| **8616** | Slc16a5 | 4295 | -1.3944835 | 4.19882702 | 62.8429341 | 1.30E-06 | 0.00018134 |
| **3371** | Fut7 | 2965 | -1.3876793 | 2.42627185 | 19.5896252 | 0.00054407 | 0.0078794 |
| **334** | Abhd15 | 3249 | -1.373086 | 2.56652434 | 20.728825 | 0.00042523 | 0.00668058 |
| **8409** | Sell | 3206 | -1.3600023 | 7.82783504 | 134.67621 | 1.11E-08 | 1.13E-05 |
| **10211** | Tubb2b | 1922 | -1.353356 | 1.93558189 | 21.701956 | 0.00034679 | 0.0059138 |
| **10149** | Tspan5 | 15327 | -1.3496273 | 4.37340535 | 27.4914865 | 0.00011492 | 0.00268414 |
| **3657** | Gm19765 | 727 | -1.3489822 | 2.589609 | 10.8042365 | 0.00525213 | 0.03625291 |
| **4233** | H2-Ob | 3031 | -1.3484509 | 3.35296376 | 37.1031974 | 2.51E-05 | 0.00102673 |
| **9764** | Tmprss13 | 3754 | -1.3303084 | 0.98483461 | 9.79945415 | 0.00719396 | 0.0447937 |
| **8975** | Sorcs2 | 16057 | -1.3157149 | 0.3987211 | 9.93739865 | 0.00688375 | 0.04331876 |
| **5006** | Klrb1f | 4250 | -1.3131593 | 1.35354939 | 13.0306202 | 0.00274485 | 0.02359776 |
| **4129** | Gramd4 | 8624 | -1.3117103 | 4.74143552 | 108.760646 | 4.41E-08 | 1.91E-05 |

| | | | | | | | |
|---|---|---|---|---|---|---|---|
| **9252** | Susd3 | 3651 | -1.2988826 | 6.78327719 | 176.23771 | 1.88E-09 | 7.07E-06 |
| **3860** | Gm39876 | 2630 | -1.2985091 | 0.50229378 | 11.7069833 | 0.00400683 | 0.03046781 |
| **1832** | Chil5 | 5633 | -1.2983081 | 1.12907946 | 10.4031935 | 0.00594433 | 0.03935639 |
| **5040** | Krt83 | 1757 | -1.297246 | 2.77655783 | 17.0749005 | 0.00096867 | 0.01170099 |
| **7987** | Rnf144a | 5805 | -1.295266 | 4.25057911 | 57.5993075 | 2.17E-06 | 0.00022793 |
| **6822** | Pear1 | 4715 | -1.2950979 | 5.22913848 | 136.897922 | 9.97E-09 | 1.13E-05 |
| **4564** | Ifi214 | 1294 | -1.2905888 | 3.25160762 | 12.1834172 | 0.00348829 | 0.02779279 |
| **2440** | Dennd3 | 5647 | -1.2895906 | 2.11116815 | 20.0245912 | 0.0004947 | 0.00740604 |
| **8609** | Slc14a1 | 3995 | -1.2889912 | 3.62394908 | 44.5417664 | 9.33E-06 | 0.00055587 |
| **2269** | Cyp2s1 | 2623 | -1.2862483 | 2.80497555 | 19.0136346 | 0.00061836 | 0.00859442 |
| **8416** | Sema4f | 5004 | -1.2853837 | 5.30036935 | 76.9092012 | 3.85E-07 | 8.18E-05 |
| **8776** | Slc9a9 | 3492 | -1.279379 | 6.35309658 | 114.553684 | 3.16E-08 | 1.55E-05 |
| **6908** | Phf11a | 1549 | -1.2786308 | 1.82502396 | 11.5731234 | 0.00416807 | 0.03130362 |
| **5238** | Lgmn | 2008 | -1.2784289 | 4.15564937 | 66.1154764 | 9.60E-07 | 0.00014227 |
| **903** | Asb2 | 2899 | -1.276297 | 3.67057423 | 18.1440432 | 0.00075358 | 0.00974035 |
| **932** | Atcay | 4233 | -1.2745022 | 3.00041976 | 31.2786109 | 6.06E-05 | 0.00178259 |
| **3777** | Gm34702 | 1424 | -1.2704249 | 1.13377371 | 9.94166679 | 0.0068744 | 0.04331858 |
| **8907** | Snord118 | 137 | -1.262379 | 4.52247343 | 11.3728033 | 0.00442347 | 0.03254868 |
| **7210** | Pou2af1 | 2566 | -1.2571784 | 5.463052 | 47.4712452 | 6.55E-06 | 0.00045773 |
| **1351** | Cacna2d4 | 6480 | -1.2532665 | 3.14763316 | 24.4866215 | 0.00019952 | 0.00404716 |
| **1952** | Cmpk2 | 3439 | -1.2517301 | 3.52992537 | 38.8300567 | 1.97E-05 | 0.00087958 |
| **4759** | Irs2 | 6794 | -1.2426032 | 2.81431642 | 9.78775857 | 0.007221 | 0.04486423 |
| **404** | Acsbg1 | 3375 | -1.2320422 | 2.37167623 | 15.9440991 | 0.00127631 | 0.01407314 |
| **8607** | Slc12a8 | 4758 | -1.2301255 | 1.2604815 | 10.9443469 | 0.00503248 | 0.03527805 |
| **4561** | Ifi209 | 3497 | -1.2289127 | 6.92866446 | 39.111536 | 1.89E-05 | 0.00087159 |
| **8405** | Selenop | 2218 | -1.2265153 | 7.0429226 | 120.964141 | 2.23E-08 | 1.25E-05 |
| **6542** | Oas1a | 2010 | -1.2239679 | 3.60886191 | 40.8254796 | 1.50E-05 | 0.00076811 |
| **5376** | Ly6a | 1002 | -1.2227304 | 8.1978266 | 27.3698834 | 0.00011742 | 0.00271993 |
| **9450** | Tent5c | 5680 | -1.2195052 | 4.73991352 | 62.479347 | 1.34E-06 | 0.00018134 |
| **10024** | Trib3 | 2051 | -1.2170866 | 1.2479051 | 17.1295508 | 0.00095611 | 0.011599 |
| **1077** | B3gnt5 | 5377 | -1.2064547 | 2.28907375 | 12.1593559 | 0.00351256 | 0.02792682 |
| **1585** | Cd27 | 1812 | -1.2052136 | 7.21666911 | 66.2646863 | 9.48E-07 | 0.00014224 |
| **3769** | Gm34150 | 1228 | -1.2050086 | 2.43735622 | 10.8857792 | 0.00512299 | 0.03560566 |
| **4001** | Gmfg | 2451 | -1.1992368 | 7.78882241 | 122.733361 | 2.03E-08 | 1.25E-05 |
| **3713** | Gm30693 | 3068 | -1.1977651 | 0.87718342 | 10.0858596 | 0.00656694 | 0.0420299 |
| **1062** | B230118I11Rik | 7105 | -1.19742 | 4.22538866 | 27.5226549 | 0.00011429 | 0.0026838 |
| **10023** | Trib2 | 4448 | -1.1959471 | 7.46199316 | 95.7586264 | 9.89E-08 | 3.01E-05 |
| **4385** | Hid1 | 6349 | -1.1947529 | 3.16333534 | 18.9892395 | 0.00062175 | 0.00859912 |
| **1125** | Baiap3 | 6167 | -1.1848971 | 1.80433672 | 12.6398893 | 0.00306248 | 0.02553885 |
| **9283** | Syt11 | 5783 | -1.1765393 | 6.80594671 | 222.19742 | 3.97E-10 | 4.47E-06 |

| | | | | | | | |
|---|---|---|---|---|---|---|---|
| **6024** | Mturn | 5307 | -1.1750349 | 3.38364715 | 23.6414256 | 0.0002349 | 0.00452054 |
| **6324** | Nipal1 | 4205 | -1.1718683 | 4.13826661 | 46.8497314 | 7.05E-06 | 0.00048376 |
| **3987** | Gm8369 | 2129 | -1.1680253 | 5.11615135 | 33.804262 | 4.08E-05 | 0.00137106 |
| **389** | Acot11 | 5876 | -1.1665666 | 3.46258183 | 45.3311475 | 8.47E-06 | 0.0005355 |
| **1128** | Bambi-ps1 | 1215 | -1.1647366 | 5.77044124 | 67.2291056 | 8.69E-07 | 0.00013382 |
| **4840** | Jun | 3187 | -1.1630265 | 6.60761427 | 12.0023727 | 0.0036757 | 0.02881688 |
| **1173** | Bcl6 | 5043 | -1.1625998 | 5.89035475 | 49.4538835 | 5.20E-06 | 0.00038507 |
| **8721** | Slc37a2 | 4436 | -1.1581557 | 1.50489611 | 12.7387593 | 0.0029783 | 0.02504587 |
| **7940** | Ripor2 | 10616 | -1.1548413 | 7.76155484 | 96.4871557 | 9.43E-08 | 2.95E-05 |
| **10121** | Trpv6 | 2928 | -1.1537691 | 1.25854111 | 11.8815899 | 0.00380715 | 0.0295856 |
| **8501** | Sgk1 | 4158 | -1.1529105 | 4.64431134 | 38.9793945 | 1.93E-05 | 0.00087262 |
| **9563** | Timp2 | 3622 | -1.1526623 | 4.75469804 | 62.7631062 | 1.31E-06 | 0.00018134 |
| **6896** | Phactr2 | 10540 | -1.1519434 | 4.32545034 | 45.0741997 | 8.74E-06 | 0.00053832 |
| **848** | Arl5c | 1673 | -1.1519388 | 6.01394814 | 69.887897 | 6.89E-07 | 0.00011746 |
| **5654** | Mettl27 | 3127 | -1.1491635 | 2.31071125 | 24.0540527 | 0.0002168 | 0.00428957 |
| **8669** | Slc26a11 | 3862 | -1.1407362 | 5.29576289 | 71.8005377 | 5.85E-07 | 0.00011163 |
| **3190** | Fbln1 | 3859 | -1.1383763 | 2.8501793 | 15.5309017 | 0.00141559 | 0.01498073 |
| **6801** | Pdlim4 | 1253 | -1.136014 | 2.25399169 | 10.7059937 | 0.00541272 | 0.03693115 |
| **3687** | Gm26740 | 12704 | -1.1254024 | 3.63131199 | 33.6050846 | 4.21E-05 | 0.0013968 |
| **5084** | LOC115490137 | 8110 | -1.1246311 | 1.10018828 | 9.87412299 | 0.0070241 | 0.04402966 |
| **3653** | Gm19585 | 1025 | -1.1233185 | 5.9329195 | 25.420912 | 0.00016729 | 0.00353346 |
| **1389** | Capn2 | 3318 | -1.1229496 | 5.75305884 | 79.4245317 | 3.16E-07 | 7.71E-05 |
| **6166** | Ncmap | 2445 | -1.1215394 | 1.72118192 | 16.4241238 | 0.00113378 | 0.01299805 |
| **4559** | Ifi206 | 3562 | -1.1146123 | 6.07755389 | 33.8340582 | 4.06E-05 | 0.00136892 |
| **10442** | Usp18 | 1771 | -1.1125008 | 3.76880542 | 14.8227957 | 0.00169676 | 0.01693451 |
| **3779** | Gm34771 | 5091 | -1.1116944 | 1.12071721 | 10.417285 | 0.0059183 | 0.03921614 |
| **3909** | Gm46289 | 4985 | -1.1027218 | 3.09573966 | 23.5315539 | 0.00024001 | 0.00457191 |
| **8268** | S1pr1 | 3029 | -1.096021 | 7.84554204 | 119.215956 | 2.45E-08 | 1.25E-05 |
| **7543** | Ptrh1 | 1079 | -1.0914599 | 3.4039839 | 20.4867628 | 0.00044777 | 0.00691527 |
| **4133** | Grb7 | 4243 | -1.0870159 | 1.92422992 | 15.279215 | 0.00150893 | 0.01559918 |
| **3843** | Gm38906 | 1321 | -1.0833104 | 2.18996347 | 11.3550067 | 0.00444702 | 0.03261535 |
| **2136** | Crmp1 | 3436 | -1.0813761 | 3.18042304 | 15.3234508 | 0.00149202 | 0.015469 |
| **9424** | Tcp11l2 | 2400 | -1.0751392 | 6.60946122 | 27.9962848 | 0.00010517 | 0.00251927 |
| **1172** | Bcl3 | 1846 | -1.0746841 | 5.0410426 | 18.8890707 | 0.00063591 | 0.00867767 |
| **4432** | Hmox1 | 1576 | -1.074216 | 2.38977594 | 12.641391 | 0.00306119 | 0.02553885 |
| **3955** | Gm51770 | 18656 | -1.0734471 | 2.8463422 | 13.9436694 | 0.00213929 | 0.01999442 |
| **831** | Arid5b | 9976 | -1.072777 | 5.56697333 | 39.4526028 | 1.81E-05 | 0.00085162 |
| **1431** | Castor2 | 4769 | -1.0692889 | 2.22151082 | 15.7366092 | 0.00134419 | 0.01452288 |
| **3814** | Gm36681 | 4004 | -1.0673245 | 1.60767056 | 11.8809007 | 0.00380792 | 0.0295856 |
| **1857** | Chst10 | 3248 | -1.0636351 | 5.38117609 | 101.321315 | 6.92E-08 | 2.46E-05 |

| | | | | | | | |
|---|---|---|---|---|---|---|---|
| **1630** | Cdc14b | 11141 | -1.0633632 | 3.38428602 | 28.4780829 | 9.67E-05 | 0.00235238 |
| **8309** | Satb1 | 13246 | -1.0576746 | 7.44349064 | 52.6573439 | 3.64E-06 | 0.00032244 |
| **4013** | Gna15 | 2261 | -1.0538309 | 3.72609965 | 19.5860356 | 0.0005445 | 0.0078794 |
| **2714** | E230032D23Rik | 1195 | -1.0505171 | 1.99298814 | 9.42564104 | 0.00811879 | 0.04880097 |
| **8229** | Rtn4rl1 | 3502 | -1.0481663 | 2.33019283 | 15.3766183 | 0.00147199 | 0.01532997 |
| **7730** | Rasa3 | 4205 | -1.0468864 | 7.05868497 | 56.5442418 | 2.41E-06 | 0.00025145 |
| **4538** | Icos | 4325 | -1.0460841 | 9.17990026 | 126.188932 | 1.69E-08 | 1.13E-05 |
| **3612** | Gm14718 | 758 | -1.045442 | 3.59901095 | 20.9119689 | 0.00040903 | 0.00655966 |
| **5049** | L1cam | 5693 | -1.0375567 | 2.47748828 | 9.5437811 | 0.00781246 | 0.04736281 |
| **9205** | Stx1a | 3425 | -1.03659 | 1.13751281 | 10.2193736 | 0.00629627 | 0.04083146 |
| **7822** | Rd3 | 9842 | -1.0361388 | 1.64488488 | 12.2568769 | 0.0034154 | 0.02746208 |
| **7049** | Plcb4 | 12740 | -1.0180632 | 3.13813863 | 21.0362702 | 0.00039844 | 0.00644492 |
| **8823** | Smco4 | 1545 | -1.0175266 | 5.32301358 | 34.1528563 | 3.87E-05 | 0.00134583 |
| **1606** | Cd53 | 2802 | -1.0163095 | 9.43415535 | 46.6007431 | 7.26E-06 | 0.00049522 |
| **2499** | Dhx58 | 2480 | -1.0120946 | 4.24545542 | 29.9953921 | 7.48E-05 | 0.00202638 |
| **7873** | Rflnb | 3512 | -1.0101695 | 6.13011742 | 36.0208446 | 2.93E-05 | 0.00112634 |
| **9026** | Spo11 | 3310 | -1.008342 | 2.61889336 | 12.1094074 | 0.00356355 | 0.02821918 |
| **3455** | Gbp7 | 5622 | 1.00423161 | 4.10912547 | 16.6037481 | 0.00108519 | 0.012582 |
| **25** | 1500009L16Rik | 1363 | 1.01253053 | 3.59344994 | 11.4663097 | 0.00430208 | 0.03202502 |
| **9797** | Tnfrsf8 | 3496 | 1.01472442 | 4.69416292 | 9.94092387 | 0.00687603 | 0.04331858 |
| **10402** | Unc93b1 | 2281 | 1.01635615 | 5.57581808 | 119.514639 | 2.41E-08 | 1.25E-05 |
| **2732** | Ebi3 | 1171 | 1.02766134 | 5.2490729 | 25.4824995 | 0.00016538 | 0.00350635 |
| **6066** | Myo1e | 4998 | 1.04251792 | 3.58045142 | 13.4536255 | 0.00244275 | 0.0217052 |
| **5365** | Lta | 1627 | 1.05164485 | 6.08441298 | 111.017824 | 3.87E-08 | 1.81E-05 |
| **1863** | Chsy1 | 4446 | 1.06062596 | 2.98389145 | 17.8033522 | 0.00081556 | 0.01036295 |
| **6940** | Phtf2 | 5149 | 1.06280717 | 4.25561472 | 39.0081576 | 1.92E-05 | 0.00087262 |
| **9070** | Srm | 1328 | 1.06949868 | 6.21045074 | 25.5178822 | 0.0001643 | 0.00350316 |
| **526** | Ahr | 5494 | 1.07205387 | 5.46506401 | 31.6319867 | 5.73E-05 | 0.00173862 |
| **7336** | Prelid2 | 724 | 1.07672914 | 2.69517863 | 25.5073156 | 0.00016462 | 0.00350341 |
| **1197** | Bhlhe40 | 3113 | 1.09435021 | 6.96744321 | 11.3564142 | 0.00444515 | 0.03261535 |
| **6432** | Nrp1 | 9735 | 1.09494865 | 6.43102403 | 26.9349992 | 0.00012688 | 0.00287396 |
| **5984** | Mtfp1 | 1396 | 1.10807771 | 3.10096307 | 19.5683283 | 0.00054663 | 0.00789981 |
| **8885** | Snhg15 | 592 | 1.12018743 | 4.19415084 | 49.5823144 | 5.12E-06 | 0.00038452 |
| **9487** | Tg | 8695 | 1.12535186 | 3.2563837 | 9.65896513 | 0.00752655 | 0.04602695 |
| **161** | 4930481B07Rik | 7228 | 1.12854527 | 1.70000258 | 17.7078884 | 0.00083396 | 0.01051365 |
| **9347** | Tasl | 4367 | 1.12955953 | 1.98177493 | 16.9888445 | 0.00098884 | 0.01183185 |
| **3802** | Gm3636 | 5353 | 1.13305087 | 3.44482034 | 21.249085 | 0.00038103 | 0.00624402 |
| **5737** | Mir17hg | 2339 | 1.16215701 | 1.35651746 | 10.3893434 | 0.00597005 | 0.03944296 |
| **2242** | Cxcr3 | 1609 | 1.16232062 | 5.0310838 | 18.4196721 | 0.00070736 | 0.00932495 |
| **1167** | Bcl2l11 | 9973 | 1.17181637 | 5.57760232 | 55.5665718 | 2.67E-06 | 0.00026356 |

| | | | | | | | |
|---|---|---|---|---|---|---|---|
| **8185** | Rreb1 | 14627 | 1.17312734 | 3.3502781 | 26.0859406 | 0.00014796 | 0.00322814 |
| **4754** | Irf8 | 3435 | 1.18174776 | 6.07930101 | 79.2009346 | 3.22E-07 | 7.71E-05 |
| **4454** | Homer1 | 18067 | 1.18266538 | 3.0146741 | 37.2954236 | 2.44E-05 | 0.00101377 |
| **1618** | Cd83 | 2141 | 1.18573178 | 7.67802311 | 51.9930008 | 3.91E-06 | 0.00033614 |
| **2359** | Dctd | 3046 | 1.18808002 | 3.48979799 | 26.3464896 | 0.00014109 | 0.00313293 |
| **1861** | Chst15 | 5922 | 1.19930067 | 3.94293535 | 41.2749457 | 1.42E-05 | 0.00075785 |
| **793** | Arhgap24 | 14407 | 1.21338661 | 1.58737206 | 17.2570489 | 0.00092751 | 0.01134993 |
| **1103** | BC049352 | 5050 | 1.22034071 | 3.30870112 | 18.9900348 | 0.00062164 | 0.00859912 |
| **4575** | Ifitm2 | 1666 | 1.2223931 | 2.64590749 | 18.6404666 | 0.00067267 | 0.00905175 |
| **4396** | Hip1 | 25858 | 1.22296324 | 0.67776803 | 9.92459971 | 0.00691188 | 0.04347145 |
| **3923** | Gm46762 | 4775 | 1.22951097 | 2.34106246 | 24.9919457 | 0.00018128 | 0.00375166 |
| **745** | Apol10b | 2744 | 1.23026509 | 1.30154248 | 12.4565269 | 0.00322594 | 0.02643202 |
| **3730** | Gm31813 | 2673 | 1.233118 | 2.3381595 | 21.4076622 | 0.00036862 | 0.00613285 |
| **5736** | Mir155hg | 1264 | 1.25909444 | 2.54073862 | 36.7762347 | 2.63E-05 | 0.00106046 |
| **7138** | Pogk | 10686 | 1.26345155 | 1.5100971 | 10.4366319 | 0.00588277 | 0.03907271 |
| **8909** | Snord14c | 87 | 1.26709354 | 0.45958547 | 10.322223 | 0.00609649 | 0.04005348 |
| **3179** | Fasl | 1935 | 1.27135368 | 3.50292396 | 13.4801177 | 0.00242514 | 0.02158276 |
| **1613** | Cd74 | 1415 | 1.28397652 | 5.00194685 | 59.6416475 | 1.77E-06 | 0.00020723 |
| **3453** | Gbp5 | 3021 | 1.28977302 | 2.69976272 | 27.599639 | 0.00011275 | 0.00266103 |
| **1285** | Bspry | 2211 | 1.2979994 | 2.27375115 | 18.9948377 | 0.00062097 | 0.00859912 |
| **2676** | Dusp22 | 3058 | 1.30565331 | 3.00324204 | 35.9399681 | 2.97E-05 | 0.00113583 |
| **1807** | Chchd10 | 1555 | 1.31522781 | 4.78456835 | 133.227581 | 1.19E-08 | 1.13E-05 |
| **1873** | Ciart | 2394 | 1.31878378 | 2.48132315 | 11.9637224 | 0.00371719 | 0.02904103 |
| **3689** | Gm26885 | 2969 | 1.34127413 | 0.45290632 | 12.6315443 | 0.00306971 | 0.02554238 |
| **7609** | Rab11fip5 | 6171 | 1.34943285 | 2.02460778 | 18.6384909 | 0.00067297 | 0.00905175 |
| **10215** | Tubb6 | 1744 | 1.35909406 | 1.54357534 | 11.8026376 | 0.00389599 | 0.02988435 |
| **8963** | Soat2 | 2283 | 1.37104287 | 1.6505686 | 11.1972596 | 0.00466214 | 0.03364514 |
| **6835** | Per2 | 6106 | 1.39334152 | 2.04039323 | 21.0005591 | 0.00040145 | 0.00646573 |
| **3268** | Fgr | 4030 | 1.41079799 | 3.60657815 | 42.2726143 | 1.24E-05 | 0.0006938 |
| **7091** | Plpp1 | 1765 | 1.43622891 | 2.9665773 | 38.1880277 | 2.15E-05 | 0.00093549 |
| **1148** | Bcam | 2407 | 1.4674915 | 1.67531789 | 15.5660223 | 0.00140309 | 0.01490592 |
| **1181** | Bco2 | 2565 | 1.47367563 | 0.19728098 | 9.99067049 | 0.00676809 | 0.04287853 |
| **9417** | Tcf4 | 44311 | 1.50968963 | 2.59448568 | 48.1086897 | 6.07E-06 | 0.00043277 |
| **7540** | Ptprk | 7097 | 1.51644089 | 3.35364267 | 50.3116268 | 4.72E-06 | 0.0003662 |
| **4644** | Il2ra | 4428 | 1.57572612 | 4.84822893 | 36.2859423 | 2.82E-05 | 0.0011103 |
| **6832** | Penk | 1593 | 1.58598732 | 4.03502224 | 23.6472989 | 0.00023463 | 0.00452054 |
| **8252** | Rxra | 5703 | 1.60426073 | 6.15208053 | 126.059769 | 1.71E-08 | 1.13E-05 |
| **7895** | Rgs16 | 2335 | 1.60801844 | 4.67857459 | 21.2131384 | 0.00038391 | 0.00627293 |
| **8241** | Runx2 | 8554 | 1.63226989 | 3.27897522 | 50.7675516 | 4.48E-06 | 0.00036263 |
| **6156** | Ncf1 | 2969 | 1.63677638 | 2.84469277 | 39.4963297 | 1.80E-05 | 0.00085012 |

| | | | | | | | |
|---|---|---|---|---|---|---|---|
| **10516** | Vcam1 | 3398 | 1.65011123 | 2.43170814 | 18.0360884 | 0.00077262 | 0.0099294 |
| **7903** | Rhbdf1 | 4991 | 1.65023518 | 0.64664623 | 13.3810135 | 0.00249179 | 0.02204561 |
| **1886** | Cish | 4293 | 1.69719141 | 4.6926159 | 98.5098629 | 8.27E-08 | 2.66E-05 |
| **4369** | Hes5 | 4386 | 1.71563798 | 1.41650982 | 15.0826629 | 0.00158674 | 0.01613725 |
| **4106** | Gpr25 | 2013 | 1.74308958 | 1.4501127 | 15.4884548 | 0.00143086 | 0.01506882 |
| **3534** | Gjb2 | 2404 | 1.74368284 | 2.23954158 | 32.6651219 | 4.87E-05 | 0.00153854 |
| **8833** | Smim10l2a | 3133 | 1.76972358 | 2.61477056 | 40.4119235 | 1.59E-05 | 0.00079477 |
| **227** | 9530036M11Rik | 12763 | 1.78439699 | 1.08152321 | 22.5881254 | 0.00028947 | 0.00524024 |
| **6243** | Nedd4 | 7090 | 1.82473325 | 1.16049383 | 21.1383221 | 0.00038998 | 0.00633537 |
| **3932** | Gm4956 | 1148 | 1.88297472 | 2.30027169 | 11.2968673 | 0.00452495 | 0.03299348 |
| **6430** | Nrn1 | 1769 | 1.93139087 | 3.485174 | 57.7441118 | 2.13E-06 | 0.00022673 |
| **2216** | Ctsw | 1498 | 1.99721329 | 2.78389752 | 36.0243602 | 2.93E-05 | 0.00112634 |
| **8561** | Sik1 | 4553 | 2.00235374 | 3.81307212 | 80.6959443 | 2.87E-07 | 7.34E-05 |
| **4579** | Ifng | 1210 | 2.0334309 | 1.10096174 | 9.35941078 | 0.00829655 | 0.04947169 |
| **5005** | Klrb1c | 2632 | 2.09446233 | 4.3482647 | 131.523648 | 1.29E-08 | 1.13E-05 |
| **4631** | Il18 | 2593 | 2.09489074 | 1.87278836 | 62.0710307 | 1.40E-06 | 0.00018289 |
| **9188** | Stra6 | 4384 | 2.09909335 | 2.24908988 | 37.6046669 | 2.34E-05 | 0.00098858 |
| **3747** | Gm32643 | 14800 | 2.21412635 | 1.31830002 | 9.42922634 | 0.00810929 | 0.04879443 |
| **1474** | Ccdc148 | 25202 | 2.22568437 | 0.61584419 | 19.1653374 | 0.00059773 | 0.00839594 |
| **1590** | Cd300lf | 1908 | 2.2459784 | 1.19569703 | 37.4668924 | 2.38E-05 | 0.0010005 |
| **5411** | Macc1 | 4301 | 2.25886106 | 0.74822596 | 18.4753838 | 0.00069842 | 0.0092722 |
| **3825** | Gm3739 | 5766 | 2.26776736 | 3.08361781 | 14.9177804 | 0.00165556 | 0.0165969 |
| **2062** | Coro2a | 5326 | 2.32184323 | 3.08386702 | 39.101404 | 1.90E-05 | 0.00087159 |
| **536** | Ajuba | 3503 | 2.32802921 | 1.23541131 | 34.0179054 | 3.95E-05 | 0.00135137 |
| **8686** | Slc30a2 | 3483 | 2.36181583 | 0.5632663 | 18.9106193 | 0.00063283 | 0.00866956 |
| **3783** | Gm35113 | 9767 | 2.37228914 | 0.93209053 | 39.015695 | 1.92E-05 | 0.00087262 |
| **7035** | Pla1a | 3899 | 2.47305598 | 0.53796396 | 21.2843533 | 0.00037823 | 0.00620715 |
| **9401** | Tbx21 | 2552 | 2.47481029 | 2.66108488 | 70.5672723 | 6.50E-07 | 0.00011608 |
| **1401** | Car2 | 1947 | 2.57683089 | 1.8437799 | 54.2748008 | 3.06E-06 | 0.00029167 |
| **4562** | Ifi211 | 2007 | 2.59817233 | 3.05452748 | 42.013613 | 1.29E-05 | 0.00070983 |
| **6336** | Nkg7 | 813 | 2.62761408 | 3.65527764 | 85.7295913 | 1.97E-07 | 5.17E-05 |
| **1620** | Cd86 | 3513 | 2.6757483 | 4.22996908 | 142.584475 | 7.64E-09 | 1.13E-05 |
| **3980** | Gm6637 | 1502 | 2.73644803 | 1.47491422 | 39.6769142 | 1.75E-05 | 0.0008437 |
| **7043** | Plac8 | 3910 | 2.87202474 | 3.2412477 | 34.7480787 | 3.54E-05 | 0.00128481 |
| **1581** | Cd22 | 4689 | 3.0155816 | 1.82758598 | 45.2240762 | 8.58E-06 | 0.00053832 |
| **8450** | Serpinc1 | 2088 | 3.09324759 | 3.07357395 | 42.8972147 | 1.15E-05 | 0.000656 |
| **8270** | S1pr3 | 4484 | 3.80487422 | 1.37521688 | 71.2137065 | 6.15E-07 | 0.00011346 |
| **10673** | Wnt10a | 1962 | 3.9747882 | 2.94282867 | 129.885483 | 1.40E-08 | 1.13E-05 |
| **5003** | Klk1b27 | 889 | 4.92340395 | 0.96326896 | 13.0346845 | 0.00342718 | 0.02748088 |
| **8493** | Sftpc | 804 | 5.57877491 | 1.9255896 | 10.2116378 | 0.0063116 | 0.0409073 |

**DAY 5**

| | gene | length | logFC | logCPM | F | PValue | FDR |
|---|---|---|---|---|---|---|---|
| 1574 | Cd163l1 | 4508 | -4.047131 | 0.07571365 | 20.198662 | 0.00047639 | 0.01746981 |
| 8042 | Rorc | 3564 | -3.0226864 | 2.93368458 | 104.786506 | 5.59E-08 | 0.00029218 |
| 4630 | Il17re | 3592 | -2.4474947 | 0.1107445 | 29.9357855 | 7.56E-05 | 0.00583482 |
| 1624 | Cd99 | 206 | -2.2282677 | 2.17640408 | 19.4491137 | 0.00056121 | 0.01932127 |
| 4635 | Il1r1 | 6138 | -2.0489737 | 4.20953397 | 28.8184968 | 9.13E-05 | 0.00654349 |
| 5487 | Map4k5 | 4454 | -1.8994198 | 0.57951951 | 16.4629382 | 0.00112307 | 0.02929792 |
| 7355 | Prkar2b | 3811 | -1.8366448 | 0.59588694 | 13.1894271 | 0.00262668 | 0.04641801 |
| 7934 | Riox1 | 2346 | -1.8176956 | 4.00809597 | 25.3320404 | 0.00017008 | 0.00962213 |
| 9655 | Tmem160 | 885 | -1.7679753 | 5.66662032 | 39.8923238 | 1.70E-05 | 0.00317746 |
| 2911 | Endog | 1531 | -1.701302 | 2.42686326 | 28.3945339 | 9.81E-05 | 0.00690607 |
| 7248 | Ppm1m | 2070 | -1.6229214 | 5.15379098 | 43.395905 | 1.08E-05 | 0.00258061 |
| 10724 | Yars2 | 1587 | -1.5857204 | 4.18697129 | 35.5508796 | 3.14E-05 | 0.00413677 |
| 1216 | Bloc1s3 | 2030 | -1.5834265 | 2.65584996 | 15.0353763 | 0.00160613 | 0.0351787 |
| 2949 | Epop | 2330 | -1.5674026 | 2.6246978 | 31.8424681 | 5.54E-05 | 0.00511231 |
| 11077 | Zfp771 | 2970 | -1.5466677 | 3.69331316 | 33.7458435 | 4.12E-05 | 0.0045471 |
| 5398 | Lysmd2 | 2296 | -1.5450332 | 2.92136504 | 36.6244464 | 2.69E-05 | 0.00398767 |
| 4290 | Habp4 | 2567 | -1.5059361 | 3.91025373 | 26.9691018 | 0.0001261 | 0.0080207 |
| 9459 | Tesc | 1036 | -1.4930647 | 6.11693926 | 99.4532076 | 7.79E-08 | 0.00029218 |
| 7620 | Rab24 | 1901 | -1.4737313 | 5.58134871 | 33.4202497 | 4.33E-05 | 0.00460771 |
| 1217 | Bloc1s4 | 1306 | -1.4476587 | 3.85629351 | 40.1435698 | 1.65E-05 | 0.00314194 |
| 3422 | Gan | 3454 | -1.4412414 | 1.52733351 | 17.1482508 | 0.00095185 | 0.02607287 |
| 2544 | Dnaja4 | 5996 | -1.4172057 | 2.53162255 | 17.4589907 | 0.0008842 | 0.02506768 |
| 10267 | Tysnd1 | 2349 | -1.3963395 | 4.57186135 | 13.6648715 | 0.00230626 | 0.04320116 |
| 8404 | Selenoo | 2428 | -1.3844575 | 3.22597838 | 32.158112 | 5.27E-05 | 0.00494341 |
| 7884 | Rfxap | 2162 | -1.3656336 | 4.69241141 | 38.6548627 | 2.02E-05 | 0.00348377 |
| 10533 | Vipr1 | 4902 | -1.3608897 | 3.40743862 | 31.4805371 | 5.87E-05 | 0.00520385 |
| 9206 | Stx2 | 5027 | -1.3540861 | 1.52709976 | 14.9704591 | 0.00163321 | 0.03556413 |
| 2719 | E2f5 | 7181 | -1.3520942 | 2.13439184 | 26.2127998 | 0.00014457 | 0.00865715 |
| 9083 | Srrd | 1455 | -1.3466924 | 3.43181662 | 34.1278572 | 3.88E-05 | 0.00443512 |
| 3553 | Glrx5 | 1002 | -1.3421808 | 5.59541065 | 26.5372958 | 0.00013629 | 0.00832734 |
| 7505 | Ptges2 | 1952 | -1.3224756 | 4.63297574 | 49.5399542 | 5.15E-06 | 0.0023563 |
| 8675 | Slc28a2b | 4043 | -1.3206661 | 5.70815583 | 27.5509128 | 0.00011372 | 0.00750134 |
| 1672 | Cdk16 | 3705 | -1.3123838 | 4.4215828 | 23.8839198 | 0.00022406 | 0.01121122 |
| 6827 | Peli1 | 3526 | -1.2976126 | 6.61842606 | 43.8370648 | 1.02E-05 | 0.00258061 |
| 9134 | St8sia6 | 3161 | -1.2886758 | 2.97598194 | 17.1938975 | 0.00094155 | 0.02600537 |
| 6737 | Pcgf6 | 4853 | -1.2871626 | 4.19466118 | 35.9732847 | 2.95E-05 | 0.0040038 |
| 7241 | Ppm1b | 6107 | -1.2731311 | 6.12903847 | 36.9070989 | 2.58E-05 | 0.00395384 |

| | | | | | | | |
|---|---|---|---|---|---|---|---|
| 5555 | Mcat | 2015 | -1.2679706 | 4.71108941 | 42.1018556 | 1.27E-05 | 0.0026948 |
| 2384 | Ddr1 | 11676 | -1.2659387 | 2.34654837 | 19.5100695 | 0.0005537 | 0.01924561 |
| 3746 | Gm32633 | 1340 | -1.2614167 | 2.28259181 | 14.7944056 | 0.0017093 | 0.03651482 |
| 1300 | Btg1 | 5010 | -1.2544779 | 8.50260197 | 30.3915082 | 7.01E-05 | 0.00563562 |
| 7141 | Poglut3 | 4806 | -1.2492478 | 2.46141655 | 17.39935 | 0.00089674 | 0.02517594 |
| 1172 | Bcl3 | 1846 | -1.2490126 | 5.0410426 | 23.9643006 | 0.0002206 | 0.01109097 |
| 9303 | Taf10 | 798 | -1.2439074 | 5.67991207 | 15.6297373 | 0.00138075 | 0.03263539 |
| 1117 | Bag1 | 1349 | -1.2421036 | 6.0052631 | 17.4486626 | 0.00088636 | 0.02506768 |
| 6366 | Noc2l | 3836 | -1.2368631 | 5.94179678 | 18.95212 | 0.00062696 | 0.02082086 |
| 3653 | Gm19585 | 1025 | -1.2265687 | 5.9329195 | 30.9746132 | 6.37E-05 | 0.00531252 |
| 2145 | Cry1 | 3977 | -1.2234167 | 4.9294029 | 47.0057562 | 6.92E-06 | 0.0025764 |
| 7886 | Rgl1 | 7507 | -1.2202131 | 0.78594939 | 13.8339312 | 0.0022033 | 0.0421905 |
| 3215 | Fbxo31 | 4626 | -1.2186609 | 3.54807083 | 19.1346487 | 0.00060184 | 0.02021677 |
| 8199 | Rrs1 | 2048 | -1.2179087 | 5.20027717 | 36.2541496 | 2.83E-05 | 0.00398767 |
| 4873 | Kcna3 | 1968 | -1.2176132 | 4.29932527 | 19.833408 | 0.00051574 | 0.01837412 |
| 10698 | Xpa | 1188 | -1.2172718 | 3.93444832 | 63.5810104 | 1.21E-06 | 0.00104948 |
| 4097 | Gpr146 | 7193 | -1.2052823 | 3.52553162 | 18.3856241 | 0.00071289 | 0.02241835 |
| 7992 | Rnf166 | 1875 | -1.2016976 | 5.76855319 | 50.1310153 | 4.81E-06 | 0.0023563 |
| 8595 | Slain2 | 4873 | -1.2008933 | 5.25452905 | 24.6815257 | 0.00019225 | 0.01025748 |
| 2710 | E130309D02Rik | 2424 | -1.1953146 | 5.0600617 | 44.9294844 | 8.90E-06 | 0.00258061 |
| 6675 | Pank2 | 4898 | -1.1932592 | 5.33743701 | 28.0311019 | 0.00010454 | 0.00713267 |
| 334 | Abhd15 | 3249 | -1.1924685 | 2.56652434 | 15.9606 | 0.00127108 | 0.03117609 |
| 8961 | Snx9 | 8321 | -1.1873782 | 1.52933604 | 14.7753479 | 0.00171778 | 0.03654753 |
| 5924 | Mrps30 | 1590 | -1.1868647 | 5.07813109 | 22.2863219 | 0.00030768 | 0.01357255 |
| 2513 | Dipk1b | 1622 | -1.1861412 | 1.69742195 | 18.8098334 | 0.00064737 | 0.02098521 |
| 10692 | Xkrx | 6775 | -1.1834801 | 2.26760969 | 17.2093232 | 0.0009381 | 0.02600537 |
| 183 | 4933439C10Rik | 2583 | -1.1691465 | 2.20456083 | 22.7824975 | 0.00027839 | 0.01275272 |
| 430 | Acvrl1 | 3788 | -1.1657654 | 3.29495343 | 14.7037604 | 0.00175006 | 0.03678605 |
| 9875 | Tprgl | 3408 | -1.1651323 | 6.34448029 | 31.392954 | 5.95E-05 | 0.00521395 |
| 7999 | Rnf187 | 1939 | -1.1598105 | 6.27643827 | 13.8949826 | 0.00216742 | 0.04194608 |
| 8905 | Snora81 | 164 | -1.155797 | 2.85167083 | 14.4369304 | 0.00187664 | 0.03855335 |
| 3400 | Gabrr2 | 3437 | -1.1458254 | 2.13832447 | 13.4501869 | 0.00244505 | 0.0447583 |
| 10452 | Usp30 | 4398 | -1.1458241 | 3.58110271 | 53.3440397 | 3.38E-06 | 0.00190088 |
| 10850 | Zdhhc7 | 3137 | -1.1444253 | 5.58734799 | 39.3422238 | 1.84E-05 | 0.00327971 |
| 7645 | Rab5if | 1051 | -1.1411832 | 6.59003567 | 31.2759415 | 6.07E-05 | 0.00521395 |
| 7006 | Pip4p1 | 1608 | -1.1280191 | 4.74828586 | 32.9985728 | 4.62E-05 | 0.00463657 |
| 4438 | Hnrnpab | 2545 | -1.1244686 | 7.69557935 | 18.8774606 | 0.00063757 | 0.02092658 |
| 7015 | Pitpna | 3795 | -1.1239966 | 7.94443378 | 48.7289277 | 5.65E-06 | 0.0023563 |
| 3221 | Fbxo4 | 3446 | -1.117073 | 3.69439116 | 32.6595305 | 4.87E-05 | 0.00472259 |
| 2141 | Crtap | 1714 | -1.1133369 | 3.3816031 | 22.5562576 | 0.00029133 | 0.01317207 |

| | | | | | | | |
|---|---|---|---|---|---|---|---|
| 7001 | Pink1 | 2429 | -1.1099029 | 5.69045373 | 39.7060274 | 1.75E-05 | 0.00317746 |
| 7181 | Polr2m | 2472 | -1.1067832 | 7.82828025 | 30.6222923 | 6.75E-05 | 0.00546498 |
| 3113 | Fam174a | 2107 | -1.1061374 | 5.13042151 | 40.7501009 | 1.52E-05 | 0.00299899 |
| 9280 | Syp | 2482 | -1.1030174 | 2.68458809 | 14.1997912 | 0.001998 | 0.04031094 |
| 7014 | Pithd1 | 1538 | -1.0980461 | 5.77735605 | 36.1409157 | 2.88E-05 | 0.00399659 |
| 7638 | Rab40c | 2466 | -1.0964501 | 3.51571241 | 32.9831684 | 4.63E-05 | 0.00463657 |
| 7361 | Prkci | 4492 | -1.0889088 | 3.32575506 | 13.0500305 | 0.00273009 | 0.04743104 |
| 4746 | Irf2bp1 | 2706 | -1.0865514 | 4.7957343 | 27.0289056 | 0.00012476 | 0.00798047 |
| 9556 | Timm29 | 3170 | -1.0856686 | 4.59207368 | 16.5337312 | 0.00110384 | 0.02906057 |
| 7253 | Ppp1cc | 2618 | -1.0775812 | 7.33637022 | 42.9283826 | 1.14E-05 | 0.00265948 |
| 1829 | Chic2 | 2550 | -1.0753569 | 5.29296991 | 20.3312729 | 0.00046296 | 0.01728858 |
| 3488 | Get4 | 3087 | -1.0648896 | 6.2187164 | 29.2306316 | 8.51E-05 | 0.00630108 |
| 9002 | Spata5l1 | 2583 | -1.062607 | 4.16839544 | 26.0578076 | 0.00014872 | 0.00871766 |
| 8797 | Smad1 | 3227 | -1.060996 | 3.59267692 | 26.2373881 | 0.00014392 | 0.00865715 |
| 2164 | Csnk1e | 7591 | -1.0524169 | 2.38562187 | 18.1437378 | 0.00075364 | 0.02279442 |
| 5530 | Matk | 3076 | -1.0502164 | 4.49657783 | 46.7937822 | 7.09E-06 | 0.0025764 |
| 7315 | Prag1 | 4800 | -1.0464119 | 3.15652725 | 29.9171614 | 7.58E-05 | 0.00583482 |
| 78 | 2300009A05Rik | 598 | -1.044374 | 3.92321652 | 36.8540747 | 2.60E-05 | 0.00395384 |
| 1688 | Cdk9 | 8915 | -1.0440708 | 6.10195406 | 38.5647907 | 2.04E-05 | 0.00348377 |
| 2096 | Cpox | 3186 | -1.0435665 | 4.90801179 | 15.1954845 | 0.00154153 | 0.03426981 |
| 4953 | Kifbp | 4512 | -1.0411635 | 4.71263658 | 18.139506 | 0.00075437 | 0.02279442 |
| 884 | Arrdc2 | 3350 | -1.0400946 | 2.9979345 | 17.1795868 | 0.00094477 | 0.02600537 |
| 4872 | Kcna2 | 14395 | -1.0381124 | 3.15002883 | 36.287546 | 2.82E-05 | 0.00398767 |
| 8964 | Socs1 | 1340 | -1.0378608 | 4.55315207 | 41.9757522 | 1.29E-05 | 0.0026948 |
| 10113 | Trp53rkb | 5238 | -1.0372893 | 2.68439603 | 16.0802027 | 0.00123391 | 0.03053056 |
| 1341 | Caap1 | 2789 | -1.0331969 | 3.62404815 | 26.0030027 | 0.00015022 | 0.00871766 |
| 2776 | Efhd2 | 2381 | -1.0315677 | 7.01252116 | 29.0925438 | 8.71E-05 | 0.00636659 |
| 3412 | Galnt10 | 4763 | -1.0265466 | 5.41738021 | 21.7851434 | 0.00034089 | 0.01449407 |
| 9162 | Stim2 | 4934 | -1.0259405 | 5.29285123 | 15.4249431 | 0.00145405 | 0.03375203 |
| 2596 | Dnttip1 | 1245 | -1.0219273 | 5.33193432 | 44.0488832 | 9.92E-06 | 0.00258061 |
| 10149 | Tspan5 | 15327 | -1.0194616 | 4.37340535 | 17.0230086 | 0.00098078 | 0.02659887 |
| 7712 | Rap1b | 1937 | -1.0054238 | 8.3736023 | 22.5917379 | 0.00028926 | 0.01313103 |
| 9501 | Tgtp1 | 2809 | 1.00735797 | 2.75166869 | 13.7719183 | 0.00224045 | 0.04246886 |
| 1103 | BC049352 | 5050 | 1.01032003 | 3.30870112 | 19.2421761 | 0.00058758 | 0.01980517 |
| 9009 | Spdl1 | 3254 | 1.01947569 | 4.72899647 | 18.8068811 | 0.0006478 | 0.02098521 |
| 5816 | Morrbid | 4492 | 1.02631108 | 4.21835941 | 24.0708992 | 0.0002161 | 0.01105251 |
| 1472 | Ccdc141 | 10566 | 1.0506747 | 1.83083758 | 13.9681221 | 0.00212532 | 0.04153961 |
| 9118 | St14 | 4454 | 1.06122286 | 4.738112 | 38.0118858 | 2.21E-05 | 0.00357941 |
| 1823 | Chdh | 5891 | 1.06740013 | 5.87254252 | 70.0965656 | 6.76E-07 | 0.00084601 |
| 1651 | Cdc6 | 5024 | 1.09952202 | 6.3059549 | 18.8509634 | 0.00064139 | 0.02098521 |

| | | | | | | | |
|---|---|---|---|---|---|---|---|
| 1327 | C1qtnf6 | 2793 | 1.11559197 | 1.71663243 | 17.108523 | 0.00096092 | 0.0262574 |
| 686 | Anxa2 | 1799 | 1.1158762 | 7.64044741 | 19.7581251 | 0.00052431 | 0.01860226 |
| 806 | Arhgap5 | 9948 | 1.11720994 | 2.65873167 | 13.1222346 | 0.00267595 | 0.04685236 |
| 9542 | Ticrr | 7199 | 1.14059346 | 5.17699305 | 38.1307315 | 2.17E-05 | 0.00357941 |
| 2925 | Entpd1 | 4966 | 1.19056254 | 2.09690811 | 13.1222101 | 0.00267597 | 0.04685236 |
| 4629 | Il17rb | 2061 | 1.20335721 | 5.90493725 | 160.24128 | 3.54E-09 | 3.99E-05 |
| 4654 | Ildr1 | 3147 | 1.22024126 | 2.24699833 | 13.7174719 | 0.00227365 | 0.04294754 |
| 5152 | LOC118568588 | 2354 | 1.25637979 | 0.43687769 | 13.348619 | 0.00251403 | 0.04551341 |
| 1620 | Cd86 | 3513 | 1.32045225 | 4.22996908 | 42.5317762 | 1.20E-05 | 0.0026948 |
| 9797 | Tnfrsf8 | 3496 | 1.36438886 | 4.69416292 | 13.3582825 | 0.00250737 | 0.04551341 |
| 5231 | Lgals3 | 1406 | 1.37075837 | 2.57119788 | 14.1665198 | 0.00201574 | 0.04037932 |
| 2260 | Cybb | 4802 | 1.45084817 | 2.86036568 | 14.1415678 | 0.00202916 | 0.04041712 |
| 6066 | Myo1e | 4998 | 1.51325069 | 3.58045142 | 27.8527664 | 0.00010785 | 0.00723272 |
| 6430 | Nrn1 | 1769 | 1.52290314 | 3.485174 | 42.008513 | 1.29E-05 | 0.0026948 |
| 9347 | Tasl | 4367 | 1.5290267 | 1.98177493 | 29.8467687 | 7.67E-05 | 0.00583482 |
| 8270 | S1pr3 | 4484 | 1.53184991 | 1.37521688 | 15.0432396 | 0.00160289 | 0.03517608 |
| 1886 | Cish | 4293 | 1.5732843 | 4.6926159 | 77.6866507 | 3.62E-07 | 0.00077696 |
| 3027 | Exo1 | 5875 | 1.6285428 | 4.67085931 | 20.2155833 | 0.00047465 | 0.01746296 |
| 10783 | Zbtb32 | 3864 | 1.80746866 | 5.90998983 | 35.1738879 | 3.32E-05 | 0.00421355 |
| 6336 | Nkg7 | 813 | 1.91810758 | 3.65527764 | 48.2329119 | 5.99E-06 | 0.0024069 |
| 1401 | Car2 | 1947 | 1.93437099 | 1.8437799 | 24.8799699 | 0.00018515 | 0.0101188 |
| 169 | 4930558J18Rik | 2052 | 1.94620379 | 1.32191784 | 19.4823181 | 0.0005571 | 0.01929801 |
| 25 | 1500009L16Rik | 1363 | 1.95491905 | 3.59344994 | 20.6112251 | 0.00043601 | 0.01698479 |
| 8861 | Smtn | 6119 | 1.99142057 | 7.25780297 | 65.0607598 | 1.06E-06 | 0.00099168 |
| 728 | Apc2 | 9678 | 2.11647656 | 0.13348117 | 13.6437025 | 0.00231954 | 0.04337768 |
| 5762 | Mlf1 | 1161 | 2.15101394 | 1.31257479 | 19.6122165 | 0.00054137 | 0.01898683 |
| 6072 | Myo6 | 8220 | 2.31941004 | 2.36673023 | 28.5843322 | 9.50E-05 | 0.00676843 |
| 1436 | Cbfa2t3 | 10458 | 2.35423709 | 0.98564224 | 14.5522288 | 0.0018207 | 0.03754111 |
| 9188 | Stra6 | 4384 | 2.42850254 | 2.24908988 | 22.5031987 | 0.00029447 | 0.01320765 |
| 3507 | Ggt1 | 2976 | 2.44095342 | 2.21607336 | 24.0501573 | 0.00021697 | 0.01105251 |
| 4106 | Gpr25 | 2013 | 2.772807 | 1.4501127 | 16.8375026 | 0.00102549 | 0.0274228 |
| 7043 | Plac8 | 3910 | 2.95718831 | 3.2412477 | 26.5394451 | 0.00013624 | 0.00832734 |
| 3583 | Gm11946 | 13145 | 3.04704806 | 0.309332 | 16.3360482 | 0.00115851 | 0.02977735 |
| 5410 | Maats1 | 2997 | 3.17107957 | 2.27028542 | 25.6779097 | 0.0001595 | 0.00906869 |
| 4497 | Hspa1a | 2798 | 3.46015544 | 5.58256726 | 12.9460878 | 0.00281024 | 0.04844969 |
