## Supplementary material for "IL-6 prevents Th2 cell polarization by promoting SOCS3-dependent suppression of IL-2 signaling": Table S2

| Regulator | Expression Log Ratio | Predicted on State | | Activation z-Score | -log10 B-H Corrected p-Value |
|---|---|---|---|---|---|
| Immunoglobulin | | complex | | -0.154 | 21.99 |
| IL2 | | cytokine | Activated | 2.488 | 20.96 |
| IL4 | | cytokine | | 1.385 | 19.26 |
| TNF | | cytokine | | 0.872 | 19.16 |
| Interferon alpha | | group | | -1.908 | 18.95 |
| IL1B | | cytokine | | -1.07 | 18.06 |
| IL21 | -2.827 | cytokine | | -1.378 | 17.84 |
| IFNG | 2.033 | cytokine | | 0.535 | 17.33 |
| STAT1 | | transcription regulator | | -1.132 | 16.51 |
| IL12 (complex) | | complex | | 0.498 | 16.10 |
| TGFB1 | | growth factor | | -1.03 | 15.51 |
| STAT3 | | transcription regulator | | -0.989 | 15.08 |
| NFATC2 | | transcription regulator | | -1.019 | 14.02 |
| Ige | | complex | | 1.65 | 13.98 |
| IFNB1 | | cytokine | | -0.182 | 13.28 |
| MAPK1 | | kinase | | 1.422 | 13.21 |
| STAT2 | | transcription regulator | | -1.563 | 13.21 |
| IFNAR1 | | transmembrane receptor | | -1.011 | 13.21 |
| SOCS1 | | other | | 0.958 | 13.19 |
| TBK1 | | kinase | | 0.25 | 13.12 |
| IL6 | | cytokine | Inhibited | -2.24 | 12.98 |
| DNASE2 | | enzyme | | 1.432 | 12.98 |
| IL12 (family) | | group | | -0.221 | 12.74 |
| Ifnar | | group | | -0.491 | 12.52 |
| CREBBP | | transcription regulator | | -1.342 | 11.97 |
| IRF7 | -1.544 | transcription regulator | Inhibited | -2.305 | 11.53 |
| MAVS | | other | Inhibited | -2.072 | 11.52 |
| TREX1 | | enzyme | Activated | 2.171 | 11.52 |
| TCF3 | | transcription regulator | | 0.311 | 10.91 |
| STING1 | | other | | -0.822 | 10.83 |
| IL7 | | cytokine | | 1.254 | 10.78 |
| TLR9 | | transmembrane receptor | | -0.087 | 10.74 |
| IFNA2 | | cytokine | | -1.834 | 10.72 |
| EP300 | | transcription regulator | | -0.64 | 10.64 |
| DUSP1 | | phosphatase | | 0.305 | 10.62 |

| | | | | | |
|---|---|---|---|---|---|
| Tcf7 | | transcription regulator | | 0.164 | 10.55 |
| ZBTB10 | | transcription regulator | | 0.453 | 10.36 |
| Irgm1 | | other | | 1.261 | 10.30 |
| PRDM1 | | transcription regulator | | 0.071 | 10.20 |
| CD3 | | complex | | 0.888 | 10.14 |
| DUSP11 | | phosphatase | Activated | 2.975 | 10.07 |
| IFN Beta | | group | | -1.642 | 9.81 |
| ID3 | | transcription regulator | | 1.117 | 9.70 |
| SMARCA5 | | transcription regulator | | -0.232 | 9.70 |
| ID2 | | transcription regulator | | 1.117 | 9.69 |
| SIRT1 | | transcription regulator | | 0.417 | 9.59 |
| TRIM24 | | transcription regulator | | 1.922 | 9.47 |
| IRF3 | | transcription regulator | Inhibited | -2.406 | 9.37 |
| IL15 | | cytokine | | 0.543 | 9.19 |
| ACKR2 | | G-protein coupled receptor | Activated | 2.111 | 9.19 |
| Ttc39aos1 | | other | Activated | 2.833 | 9.16 |
| IRF8 | 1.182 | transcription regulator | | -0.284 | 9.02 |
| KDM1A | | enzyme | | | 8.97 |
| TICAM1 | | other | | -0.393 | 8.94 |
| IFIH1 | | enzyme | Inhibited | -2.032 | 8.92 |
| STAT4 | | transcription regulator | | -1.145 | 8.86 |
| IRF9 | | transcription regulator | | -1.212 | 8.84 |
| IL27 | | cytokine | | -0.036 | 8.84 |
| IL18 | 2.095 | cytokine | | 1.311 | 8.82 |
| APP | | other | | -0.159 | 8.78 |
| mir-142 | | microRNA | | -1.012 | 8.75 |
| MAPT | | other | | -0.391 | 8.75 |
| STAT6 | | transcription regulator | | 0.48 | 8.54 |
| FOXP3 | | transcription regulator | | -0.27 | 8.54 |
| EGR2 | | transcription regulator | | -0.106 | 8.33 |
| RNASEH2B | | other | Activated | 3.219 | 8.31 |
| RUNX1 | | transcription regulator | | 0.684 | 8.30 |
| ETV6-RUNX1 | | fusion gene/product | | 1.976 | 8.03 |
| Ifn | | group | | -1.474 | 8.03 |
| LDB1 | | transcription regulator | | -1.606 | 7.99 |
| IFN alpha/beta | | group | | -0.274 | 7.97 |
| LMO2 | | transcription regulator | | -1.606 | 7.97 |
| CD28 | | transmembrane receptor | | 1.488 | 7.88 |

| | | | | | |
|---|---|---|---|---|---|
| MYD88 | | other | | 0.766 | 7.70 |
| IFNL1 | | cytokine | Inhibited | -2.099 | 7.70 |
| cytokine | | group | | -0.114 | 7.68 |
| KLF2 | | transcription regulator | | -1.956 | 7.49 |
| TCR | | complex | | 0.61 | 7.44 |
| TLR4 | | transmembrane receptor | | 1.178 | 7.44 |
| SLC15A4 | | transporter | | -1.586 | 7.44 |
| ZBTB16 | | transcription regulator | | 1.398 | 7.34 |
| IL27RA | | transmembrane receptor | | 1.01 | 7.30 |
| IL1 | | group | | -1.244 | 7.26 |
| IFNA1/IFNA13 | | cytokine | | -1.196 | 7.26 |
| IL33 | | cytokine | | 0.948 | 7.26 |
| DDX58 | | enzyme | | -1.736 | 7.18 |
| RARA | | ligand-dependent nuclear receptor | | 1.325 | 7.05 |
| TET2 | | enzyme | | 1.474 | 7.01 |
| MIF | | cytokine | | 1.665 | 6.98 |
| IL12A | | cytokine | | 0.409 | 6.98 |
| SENP3 | | peptidase | | -1.706 | 6.98 |
| RELA | | transcription regulator | | 0.52 | 6.97 |
| NFkB (complex) | | complex | | 1.507 | 6.92 |
| BCR (complex) | | complex | | 0.728 | 6.85 |
| NFKB1 | | transcription regulator | Activated | 2.245 | 6.85 |
| PARP9 | | enzyme | | -1.937 | 6.81 |
| IKZF1 | | transcription regulator | | -0.274 | 6.77 |
| EGR3 | | transcription regulator | | -1.284 | 6.75 |
| BACH2 | | transcription regulator | | -0.566 | 6.70 |
| PNPT1 | | enzyme | Activated | 2.03 | 6.68 |
| PRL | | cytokine | | -1.807 | 6.67 |
| IRGM | | enzyme | Activated | 2.132 | 6.60 |
| TBX21 | 2.475 | transcription regulator | Activated | 2.013 | 6.52 |
| SAMSN1 | | other | | -0.905 | 6.50 |
| CD40 | | transmembrane receptor | Activated | 2.392 | 6.49 |
| CITED2 | | transcription regulator | | 1.424 | 6.47 |
| TGM2 | | enzyme | | -1.807 | 6.37 |
| NFATC1 | | transcription regulator | | 0.044 | 6.37 |
| RC3H1 | | enzyme | | 1.567 | 6.31 |
| BTK | | kinase | | -1.164 | 6.29 |
| G protein alphai | | group | | -1.394 | 6.27 |

| | | | | | |
|---|---|---|---|---|---|
| mir-24 | | microRNA | | 0.68 | 6.27 |
| BTNL2 | | transmembrane receptor | | 0.632 | 6.27 |
| IgG | | complex | | 1.177 | 6.27 |
| CGAS | | enzyme | Inhibited | -2.795 | 6.26 |
| TLR3 | | transmembrane receptor | | -0.926 | 6.26 |
| KRAS | | enzyme | | 0.487 | 6.26 |
| SPI1 | | transcription regulator | | -1.397 | 6.25 |
| SOCS3 | -2.348 | phosphatase | | 0.853 | 6.21 |
| ELAVL1 | | other | | -0.846 | 6.18 |
| STAG2 | | other | Activated | 2.825 | 6.11 |
| EIF2AK2 | | kinase | Inhibited | -2.238 | 6.04 |
| IL17A | | cytokine | | -1.297 | 6.03 |
| CSF1 | | cytokine | | -0.581 | 5.97 |
| MEF2A | | transcription regulator | Inhibited | -3.156 | 5.94 |
| SOD1 | | enzyme | | | 5.93 |
| PTGER4 | | G-protein coupled receptor | | 1.132 | 5.93 |
| TPH1 | | enzyme | Inhibited | -2.003 | 5.90 |
| NOTCH1 | | transcription regulator | | 1.698 | 5.90 |
| BHLHE40 | 1.094 | transcription regulator | Activated | 2.559 | 5.85 |
| MAPK14 | | kinase | | 0.587 | 5.83 |
| CG | | complex | | -1.551 | 5.82 |
| FOXO1 | | transcription regulator | | -0.797 | 5.81 |
| STAT5B | | transcription regulator | | 0.419 | 5.81 |
| CREB1 | | transcription regulator | | -0.254 | 5.66 |
| CD38 | | enzyme | | -0.44 | 5.65 |
| SNCA | | enzyme | | 0.093 | 5.64 |
| IL23 | | complex | | -0.319 | 5.63 |
| SATB1 | -1.058 | transcription regulator | Activated | 2.634 | 5.63 |
| NR3C1 | | ligand-dependent nuclear receptor | | 0.324 | 5.63 |
| USP8 | | peptidase | Activated | 2.27 | 5.63 |
| PGR | | ligand-dependent nuclear receptor | | 0.255 | 5.54 |
| DOCK8 | | other | | -0.333 | 5.52 |
| CTLA4 | | transmembrane receptor | | -0.843 | 5.52 |
| SASH1 | | other | | -0.333 | 5.47 |
| BCL6 | -1.163 | transcription regulator | | -1.655 | 5.42 |
| TNFSF13B | | cytokine | | 0 | 5.35 |

| | | | | | |
|---|---|---|---|---|---|
| NFKB2 | | transcription regulator | | -0.24 | 5.35 |
| REL | | transcription regulator | | 0.31 | 5.32 |
| TGFBR2 | | kinase | | 0.153 | 5.31 |
| GFI1 | | transcription regulator | | 1.99 | 5.27 |
| CLEC12A | | other | | -1.98 | 5.20 |
| TAZ | | enzyme | | 0.226 | 5.19 |
| STAT5A | | transcription regulator | | 1.778 | 5.17 |
| Jnk | | group | | 0.232 | 5.15 |
| MTOR | | kinase | | 1.49 | 5.15 |
| IL5 | | cytokine | | -0.389 | 5.13 |
| GLI1 | | transcription regulator | | 1.487 | 5.09 |
| SMAD3 | | transcription regulator | Inhibited | -2.05 | 5.05 |
| CD274 | | enzyme | | -1.872 | 5.02 |
| RNASEH2A | | enzyme | | | 5.02 |
| NONO | | transcription regulator | Inhibited | -2.105 | 5.02 |
| PIK3CG | | kinase | | -1.014 | 5.00 |
| RNY3 | | other | Inhibited | -2.449 | 4.90 |
| TERT | | enzyme | Inhibited | -2.563 | 4.88 |
| TRIM14 | | other | | 0.308 | 4.87 |
| TNFSF10 | | cytokine | | 0.235 | 4.85 |
| OSMR | | transmembrane receptor | | | 4.84 |
| LYN | | kinase | | 0.301 | 4.82 |
| IL10 | | cytokine | | -1.447 | 4.81 |
| ITK | | kinase | | -0.289 | 4.78 |
| ERK1/2 | | group | | 0.666 | 4.77 |
| MAPKAP1 | | other | | -1.342 | 4.77 |
| IRF5 | | transcription regulator | Inhibited | -2.921 | 4.74 |
| CD276 | | other | | 1.164 | 4.74 |
| CBFB | | transcription regulator | | -0.378 | 4.71 |
| FOXO4 | | transcription regulator | | -0.366 | 4.71 |
| FOXO3 | | transcription regulator | | -1.47 | 4.68 |
| RPTOR | | other | | 0.045 | 4.66 |
| PPP2CA | | phosphatase | | -0.114 | 4.66 |
| TNFSF11 | | cytokine | | 0.222 | 4.63 |
| ETV6-NTRK3 | | fusion gene/product | Inhibited | -2 | 4.62 |
| ULBP1 | | transmembrane receptor | Activated | 2 | 4.62 |
| CSF2 | | cytokine | | 0.988 | 4.58 |
| POLG2 | | enzyme | | | 4.52 |
| Ap1 | | complex | | 0.475 | 4.50 |

| | | | | | |
|---|---|---|---|---|---|
| ICOS | -1.046 | transmembrane receptor | | -0.39 | 4.49 |
| TLR7 | | transmembrane receptor | | -0.53 | 4.48 |
| IL1A | | cytokine | | -0.072 | 4.47 |
| SMARCA4 | | transcription regulator | | -0.176 | 4.44 |
| IFNA4 | | cytokine | | -0.568 | 4.44 |
| IFN type 1 | | group | | -0.754 | 4.44 |
| P38 MAPK | | group | | 0.464 | 4.39 |
| CSF3 | | cytokine | | -0.43 | 4.38 |
| IRF4 | | transcription regulator | | 0.529 | 4.38 |
| JAK1 | | kinase | | -1.471 | 4.37 |
| IPMK | | kinase | | -0.032 | 4.33 |
| PTPN11 | | phosphatase | | -0.005 | 4.31 |
| PTEN | | phosphatase | | 1.612 | 4.30 |
| TSC22D3 | | transcription regulator | | -0.886 | 4.28 |
| Tgf beta | | group | | -0.384 | 4.24 |
| NGLY1 | | enzyme | Activated | 2.386 | 4.22 |
| IKZF2 | | transcription regulator | | 1.3 | 4.15 |
| ARNT | | transcription regulator | | 0.555 | 4.15 |
| IRF1 | | transcription regulator | | -0.888 | 4.15 |
| Vegf | | group | | -0.693 | 4.14 |
| MYC | | transcription regulator | Activated | 2.935 | 4.14 |
| IL10RA | | transmembrane receptor | | 0.742 | 4.14 |
| KLF6 | | transcription regulator | | 0.322 | 4.14 |
| CD40LG | | cytokine | | 1.919 | 4.10 |
| EBI3 | 1.028 | cytokine | | 0.734 | 4.10 |
| CD14 | | transmembrane receptor | | 0.218 | 4.10 |
| PARP1 | | enzyme | Activated | 2.594 | 4.05 |
| NSD2 | | enzyme | | -1.342 | 4.04 |
| Histone h3 | | group | | | 4.04 |
| TCF12 | | transcription regulator | | 0.356 | 4.03 |
| ESR2 | | ligand-dependent nuclear receptor | Activated | 2.224 | 4.02 |
| IL23A | | cytokine | | | 3.99 |
| SOX4 | 1.006 | transcription regulator | | -0.543 | 3.98 |
| PRKAA1 | | kinase | | 0.24 | 3.93 |
| CAMP | | other | | -1.372 | 3.93 |
| CD80 | | transmembrane receptor | | -0.096 | 3.93 |
| KIT | -2.761 | transmembrane receptor | | | 3.90 |
| NFKBIA | | transcription regulator | | 0.506 | 3.89 |

| | | | | | |
|---|---|---|---|---|---|
| EGF | | growth factor | | -1.426 | 3.88 |
| TASL | 1.13 | enzyme | | -0.447 | 3.86 |
| Tlr | | group | | 0.977 | 3.85 |
| miR-182-5p (and other miRNAs w/seed UUGGCAA) | | mature microRNA | Activated | 2.621 | 3.85 |
| OGA | | enzyme | | 0.728 | 3.83 |
| IL2RG | | transmembrane receptor | | -0.258 | 3.81 |
| IL13 | | cytokine | | -0.858 | 3.80 |
| mir-96 | | microRNA | Inhibited | -2.433 | 3.77 |
| PI3K (complex) | | complex | | -0.236 | 3.76 |
| IFNL4 | | cytokine | | -1.977 | 3.75 |
| HTT | | transcription regulator | | | 3.75 |
| MAF | | transcription regulator | | -0.399 | 3.69 |
| IFNAR2 | | transmembrane receptor | | | 3.67 |
| NFAT5 | | transcription regulator | | 1.127 | 3.67 |
| TP63 | | transcription regulator | | 0.849 | 3.66 |
| TNFSF4 | | cytokine | | -0.829 | 3.64 |
| ACTL6A | | other | | | 3.64 |
| RAG1 | | enzyme | | | 3.62 |
| mir-21 | | microRNA | | -0.638 | 3.61 |
| BCL11B | | transcription regulator | | 1.115 | 3.59 |
| IFNL3 | | cytokine | | -0.665 | 3.57 |
| EPOR | | transmembrane receptor | | | 3.57 |
| DNMT3A | | enzyme | | 0.412 | 3.53 |
| SPIB | | transcription regulator | | -1.134 | 3.52 |
| ZBTB7B | | transcription regulator | | 0.849 | 3.51 |
| KLF3 | | transcription regulator | Activated | 2.537 | 3.51 |
| F2 | | peptidase | | -1.093 | 3.50 |
| POMC | | other | | -1.099 | 3.49 |
| Akt | | group | | 0.352 | 3.47 |
| CTNNB1 | | transcription regulator | | -0.12 | 3.47 |
| IL1RN | | cytokine | Activated | 2.155 | 3.46 |
| VCAN | | other | | 0.664 | 3.45 |
| TNFRSF1A | | transmembrane receptor | | -0.447 | 3.45 |
| IGF1 | | growth factor | | 0.726 | 3.45 |
| USP18 | -1.113 | peptidase | | 1.387 | 3.45 |
| ITGB2 | | transmembrane receptor | | 1.664 | 3.45 |
| ATN1 | | transcription regulator | | | 3.44 |

| | | | | | |
|---|---|---|---|---|---|
| TSC2 | | other | Activated | 2.535 | 3.41 |
| IL12RB1 | | transmembrane receptor | | | 3.40 |
| mir-181 | | microRNA | | -1.054 | 3.39 |
| BAK1 | | other | | -1.432 | 3.39 |
| IL21R | | transmembrane receptor | | -0.314 | 3.39 |
| S100A9 | | other | | 0.508 | 3.38 |
| RICTOR | | other | | -0.736 | 3.35 |
| EZH2 | | transcription regulator | | 0.228 | 3.34 |
| PRKAA2 | | kinase | | 1.042 | 3.33 |
| CD44 | | other | | 0.251 | 3.33 |
| ANXA1 | | enzyme | | 0.186 | 3.33 |
| FZD9 | | G-protein coupled receptor | | -1.131 | 3.33 |
| TNK1 | | kinase | | -1 | 3.33 |
| mir-183 | | microRNA | Inhibited | -2.449 | 3.33 |
| PPP2R5C | | other | | -1 | 3.33 |
| NFKBIB | | transcription regulator | | | 3.33 |
| FAS | | transmembrane receptor | | 0.555 | 3.33 |
| CXCL8 | | cytokine | | 0.851 | 3.33 |
| HAVCR1 | | other | | -0.555 | 3.33 |
| CX3CL1 | | cytokine | | -1.242 | 3.30 |
| ATG7 | | enzyme | | 0.194 | 3.30 |
| RHOA | | enzyme | | 0.311 | 3.30 |
| SP110 | | transcription regulator | | 1 | 3.30 |
| FCGR2A | -1.103 | transmembrane receptor | | -0.557 | 3.29 |
| Secretase gamma | | complex | | 0.247 | 3.28 |
| mir-155 | | microRNA | Activated | 2.25 | 3.25 |
| CD70 | | cytokine | | | 3.24 |
| FABP5 | | transporter | | -0.339 | 3.24 |
| TP53 | | transcription regulator | | -0.681 | 3.23 |
| CD5 | | transmembrane receptor | | -0.97 | 3.21 |
| JUND | | transcription regulator | | -0.221 | 3.18 |
| RGS10 | | enzyme | | | 3.18 |
| PML-RARA | | fusion gene/product | | | 3.17 |
| Klrk1 | | transmembrane receptor | | 1.257 | 3.17 |
| IL3 | | cytokine | | 1.136 | 3.17 |
| IFNA14 | | cytokine | | -0.882 | 3.16 |
| IKBKB | | kinase | | 0.252 | 3.15 |
| POU2AF1 | -1.257 | transcription regulator | | -1.944 | 3.13 |

| | | | | | |
|---|---|---|---|---|---|
| CD4 | | transmembrane receptor | | 0.349 | 3.13 |
| IL12B | | cytokine | | 1.512 | 3.11 |
| GIP | | other | | -1.664 | 3.11 |
| FOXC1 | | transcription regulator | | -1.238 | 3.10 |
| PIM2 | | kinase | | 1.131 | 3.08 |
| MGA | | transcription regulator | | 1.633 | 3.08 |
| TNFRSF8 | 1.015 | transmembrane receptor | | -1.633 | 3.08 |
| SPP1 | | cytokine | | -0.304 | 3.06 |
| NOS2 | | enzyme | | -1.287 | 3.06 |
| immune complex | | complex | | | 3.04 |
| Duxbl1 | | other | | | 3.04 |
| SPIC | | transcription regulator | | | 3.04 |
| GAPDH | | enzyme | | 0.447 | 3.04 |
| TCF4 | 1.51 | transcription regulator | | -1.667 | 3.02 |
| MAP3K8 | | kinase | | 0.757 | 3.01 |
| HNRNPU | | transporter | | 1.664 | 3.01 |
| MVP | | other | | | 3.01 |
| AGT | | growth factor | | -0.76 | 3.00 |
| JUN | -1.163 | transcription regulator | | 1.849 | 3.00 |
| TNFSF15 | | cytokine | Activated | 2.188 | 2.99 |
| LDL | | complex | | 0.45 | 2.98 |
| HGF | | growth factor | Inhibited | -2.458 | 2.95 |
| TLR2 | | transmembrane receptor | | 1.645 | 2.95 |
| KEAP1 | | transcription regulator | | 1.253 | 2.94 |
| CREM | | transcription regulator | | -0.633 | 2.94 |
| HIF1A | | transcription regulator | | -0.039 | 2.94 |
| BMP10 | | growth factor | | 1.25 | 2.93 |
| PDLIM2 | | other | | 1.134 | 2.93 |
| HMGB1 | | transcription regulator | | 0.422 | 2.93 |
| OSM | | cytokine | | -0.184 | 2.92 |
| Tnf (family) | | group | | 0.046 | 2.92 |
| CEBPB | | transcription regulator | Inhibited | -2.764 | 2.92 |
| CD3E | | transmembrane receptor | | | 2.92 |
| STAT5a/b | | group | | 0.876 | 2.91 |
| GSE1 | | other | | | 2.91 |
| DUSP16 | | phosphatase | | | 2.91 |
| CYLD | | transcription regulator | | -0.52 | 2.91 |
| ZBED2 | | transcription regulator | | | 2.91 |
| FCGR3A/FCGR3B | | transmembrane receptor | | | 2.91 |

| | | | | | |
|---|---|---|---|---|---|
| IL12RB2 | | transmembrane receptor | | | 2.91 |
| EGLN1 | | enzyme | | 1.633 | 2.90 |
| DYSF | | other | | | 2.89 |
| NADPH oxidase | | complex | | | 2.89 |
| IFNK | | cytokine | | 0.943 | 2.89 |
| SUPT20H | | other | | 1 | 2.89 |
| STAT4 dimer | | complex | | | 2.88 |
| ANO6 | | ion channel | | | 2.88 |
| IFIT1B | -1.581 | other | | | 2.88 |
| Raet1a | | other | | | 2.88 |
| GAB3 | | other | | | 2.88 |
| RNA polymerase II | | complex | | | 2.88 |
| JAK | | group | Inhibited | -2 | 2.88 |
| ARHGAP21 | | other | | 1.342 | 2.88 |
| NFATC3 | | transcription regulator | | | 2.88 |
| IL4R | | transmembrane receptor | | 1.039 | 2.88 |
| IL6ST | | transmembrane receptor | Inhibited | -2.2 | 2.88 |
| NCOA2 | | transcription regulator | | -0.547 | 2.84 |
| IGHM | | transmembrane receptor | | | 2.84 |
| IL36B | | cytokine | | | 2.83 |
| CD81 | | other | | | 2.83 |
| NCOR1 | | transcription regulator | Activated | 2.621 | 2.83 |
| TOX | | transcription regulator | | 1 | 2.83 |
| TSLP | | cytokine | | 1.452 | 2.82 |
| CXCL12 | | cytokine | | -0.736 | 2.81 |
| FLT3 | | kinase | | | 2.80 |
| ERK | | group | | 0.562 | 2.79 |
| Stat1-Stat2 | | complex | | | 2.79 |
| RC3H2 | | enzyme | | | 2.79 |
| NR4A1 | | ligand-dependent nuclear receptor | | 1.077 | 2.77 |
| TRA | | transmembrane receptor | | | 2.77 |
| PRKACA | | kinase | | 0 | 2.76 |
| IFNLR1 | | transmembrane receptor | | | 2.76 |
| EPAS1 | | transcription regulator | | -0.426 | 2.73 |
| ISG15 | | other | Activated | 2.207 | 2.72 |
| ZC3H12A | | enzyme | | -0.867 | 2.71 |
| HRAS | | enzyme | | -1.757 | 2.71 |
| FLT3LG | | cytokine | Activated | 2.421 | 2.71 |

| | | | | | |
|---|---|---|---|---|---|
| CHD4 | | enzyme | Activated | 2.236 | 2.71 |
| Map3k7 | | kinase | | 0.168 | 2.71 |
| MAPK9 | | kinase | | 0.459 | 2.70 |
| MET | | kinase | | -0.378 | 2.70 |
| FYN | | kinase | | 0.555 | 2.69 |
| ADAM10 | | peptidase | Inhibited | -2 | 2.68 |
| IFN Lambda | | group | | | 2.68 |
| TXK | | kinase | | | 2.68 |
| MAPK3 | | kinase | | -0.928 | 2.68 |
| APOH | | transporter | | | 2.68 |
| IKBKG | | kinase | | -0.989 | 2.67 |
| ECSIT | | transcription regulator | | -1.109 | 2.66 |
| TXN | | enzyme | Inhibited | -2 | 2.66 |
| CALCA | | other | | -1.633 | 2.66 |
| Srgn | | other | | 0.447 | 2.66 |
| CISH | 1.697 | other | | -1.342 | 2.66 |
| EPO | | cytokine | | -1.367 | 2.66 |
| Mapk | | group | | 0.218 | 2.64 |
| LEP | | growth factor | | -0.177 | 2.64 |
| Pka | | complex | | 0 | 2.63 |
| NKX2-3 | | transcription regulator | | 0.681 | 2.63 |
| NEIL2 | | enzyme | | | 2.63 |
| AHR | 1.072 | ligand-dependent nuclear receptor | | -0.623 | 2.63 |
| PPARGC1A | | transcription regulator | | 0.397 | 2.63 |
| JAK3 | | kinase | | 1.342 | 2.63 |
| WNT3A | | cytokine | | -0.488 | 2.62 |
| S100A8 | | other | | 0.707 | 2.62 |
| GATA4 | | transcription regulator | | 0.673 | 2.62 |
| LIF | | cytokine | | -0.72 | 2.61 |
| ERBB2 | | kinase | | 0.163 | 2.61 |
| LRP5 | | transmembrane receptor | | 0 | 2.59 |
| STAT1/3/5 dimer | | complex | | | 2.59 |
| CCL5 | | cytokine | | -0.447 | 2.59 |
| TCIRG1 | | enzyme | | | 2.59 |
| PRKCE | | kinase | | 0.128 | 2.59 |
| HLA-G | | other | | | 2.59 |
| HMG20A | | transcription regulator | | -0.378 | 2.58 |
| CCL2 | | cytokine | | 0.218 | 2.56 |

| | | | | | |
|---|---|---|---|---|---|
| RAC2 | | enzyme | | | 2.56 |
| PRKDC | | kinase | | | 2.56 |
| IRAK4 | | kinase | | 1.078 | 2.56 |
| MAP3K7 | | kinase | | -0.218 | 2.56 |
| BATF | | transcription regulator | | | 2.56 |
| SQSTM1 | | transcription regulator | | -1 | 2.56 |
| SPHK1 | | kinase | | -0.803 | 2.53 |
| CD86 | 2.676 | transmembrane receptor | | 0.494 | 2.53 |
| PPARG | | ligand-dependent nuclear receptor | Inhibited | -2.597 | 2.51 |
| CD69 | | transmembrane receptor | | 1 | 2.51 |
| PSMD14 | | peptidase | | | 2.50 |
| IgG2a | | complex | | | 2.49 |
| PIM3 | | kinase | | | 2.49 |
| TIGIT | -2.81 | other | | | 2.49 |
| BAX | | transporter | | -1.432 | 2.49 |
| PF4 | | cytokine | | 0.603 | 2.49 |
| Focal adhesion kinase | | group | | 1.964 | 2.47 |
| TGFB2 | | growth factor | | -1.026 | 2.47 |
| TG | 1.125 | other | | | 2.47 |
| WAS | | other | | | 2.47 |
| BRD4 | | kinase | Activated | 2.2 | 2.44 |
| RUNX1-RUNX1T1 | | fusion gene/product | | | 2.42 |
| HSPD1 | | enzyme | | -0.246 | 2.42 |
| ZC3H12C | | other | | -0.152 | 2.42 |
| TRB | | transmembrane receptor | | | 2.42 |
| THRB | | ligand-dependent nuclear receptor | | -0.277 | 2.42 |
| miR-30c-5p (and other miRNAs w/seed GUAAACA) | | mature microRNA | | 1.181 | 2.41 |
| AIRE | | transcription regulator | | | 2.41 |
| SCAVENGER receptor CLASS A | | group | | | 2.41 |
| LTA | 1.052 | cytokine | | 1.474 | 2.40 |
| FOS | | transcription regulator | Activated | 2.415 | 2.40 |
| AMPK | | complex | | -0.342 | 2.39 |
| HFE | | transmembrane receptor | | -0.452 | 2.39 |
| TREM1 | | transmembrane receptor | | 1.154 | 2.38 |
| HAVCR2 | | other | | -1.103 | 2.38 |

| | | | | | |
|---|---|---|---|---|---|
| CNOT7 | | transcription regulator | | | 2.38 |
| PIK3R1 | | kinase | | 0.128 | 2.37 |
| FADD | | other | | -1.673 | 2.37 |
| HNF1B | | transcription regulator | | 1.387 | 2.35 |
| TFRC | | transporter | | 1.673 | 2.35 |
| IFNL2 | | other | | | 2.33 |
| TAP1 | | transporter | | | 2.33 |
| BDNF | | growth factor | | -0.803 | 2.33 |
| PRKCA | | kinase | | -1.117 | 2.32 |
| ROCK2 | | kinase | | | 2.32 |
| Insulin | | group | Inhibited | -2.024 | 2.31 |
| UCP1 | | transporter | | 1.463 | 2.31 |
| PRKCZ | | kinase | | 0.152 | 2.29 |
| MARK2 | | kinase | | 0 | 2.29 |
| IKBKE | | kinase | | -0.655 | 2.29 |
| PML | | transcription regulator | | -1.348 | 2.27 |
| Hdac | | group | | 1.722 | 2.26 |
| TYROBP | | transmembrane receptor | | 1.982 | 2.26 |
| Stat3-Stat3 | | complex | | -1.177 | 2.26 |
| TNFSF14 | | cytokine | | 0.851 | 2.26 |
| DLL1 | | enzyme | | 1.103 | 2.26 |
| CDC42 | | enzyme | | | 2.26 |
| MUC13 | | other | | | 2.26 |
| NLRP12 | | other | | -0.283 | 2.26 |
| AIM2 | | other | | 1.98 | 2.26 |
| Il12 receptor | | complex | | | 2.25 |
| NLRP6 | | G-protein coupled receptor | | | 2.25 |
| RAB11FIP3 | | other | | | 2.25 |
| TPP2 | | peptidase | | | 2.25 |
| PSEN1 | | peptidase | | -0.063 | 2.25 |
| RARG | -1.375 | ligand-dependent nuclear receptor | | 0.277 | 2.25 |
| ATF2 | | transcription regulator | | | 2.25 |
| FGF1 | | growth factor | | -0.705 | 2.24 |
| NR4A2 | | ligand-dependent nuclear receptor | | -0.487 | 2.23 |
| RNASEL | | enzyme | | | 2.23 |
| PRKCQ | | kinase | | 1.929 | 2.23 |

| | | | | | |
|---|---|---|---|---|---|
| MSC | | transcription regulator | | -1.342 | 2.22 |
| TAF4 | | transcription regulator | | | 2.21 |
| Integrin | | complex | | | 2.20 |
| Foxo | | group | | | 2.20 |
| USP1 | | peptidase | | | 2.20 |
| TNFRSF6B | | transmembrane receptor | | | 2.20 |
| RAS | | group | | -0.208 | 2.20 |
| NLRP3 | | other | | 0.543 | 2.20 |
| POU2F2 | | transcription regulator | | -1.633 | 2.20 |
| CD200R1 | | transmembrane receptor | | 1.342 | 2.20 |
| CNTF | | cytokine | | -1.77 | 2.19 |
| CHUK | | kinase | | 0.386 | 2.19 |
| CASZ1 | | enzyme | | 1.091 | 2.19 |
| SMARCB1 | | transcription regulator | | -0.447 | 2.16 |
| NMDA Receptor | | complex | | | 2.14 |
| XRCC6 | | enzyme | | | 2.14 |
| TRAF5 | | transporter | | | 2.14 |
| RARB | | ligand-dependent nuclear receptor | | 0.478 | 2.14 |
| IRF2 | | transcription regulator | | 1.414 | 2.14 |
| CDKN2A | | transcription regulator | | -0.421 | 2.13 |
| Pdgf (complex) | | complex | | -1.202 | 2.12 |
| F2RL1 | -1.839 | G-protein coupled receptor | | | 2.12 |
| PRKCB | | kinase | | 0.447 | 2.12 |
| ATF3 | | transcription regulator | | 0.061 | 2.12 |
| C10orf99 | | cytokine | | 0 | 2.12 |
| PRKCI | | kinase | | -0.218 | 2.08 |
| Dexamethasone-GR | | complex | | | 2.08 |
| p38 Sapk | | group | | | 2.08 |
| CYP | | group | | | 2.08 |
| Mucin | | group | | | 2.08 |
| ESR1 | 1.451 | ligand-dependent nuclear receptor | | -0.66 | 2.08 |
| EGOT | | other | | | 2.08 |
| PAEP | | other | | | 2.08 |
| ICAM5 | | other | | | 2.08 |
| LAMP1 | | other | | | 2.08 |
| IL31RA | | transmembrane receptor | | | 2.08 |

| | | | | | |
|---|---|---|---|---|---|
| HLA-B | | transmembrane receptor | | | 2.08 |
| MAPK8 | | kinase | | 0.7 | 2.08 |
| USF2 | | transcription regulator | | -0.254 | 2.08 |
| ADRB | | group | | 1.89 | 2.07 |
| ETS1 | | transcription regulator | | 0.277 | 2.06 |
| AGTR1 | | G-protein coupled receptor | | | 2.06 |
| CIITA | | transcription regulator | | -1.086 | 2.06 |
| ATF1 | | transcription regulator | | | 2.06 |
| RHO | | G-protein coupled receptor | | 0.447 | 2.06 |
| JAK1/2 | | group | | 0.447 | 2.06 |
| IKZF3 | | transcription regulator | Activated | 2.219 | 2.06 |
| RAG2 | | enzyme | | | 2.04 |
| TNFRSF1B | | transmembrane receptor | | -1.342 | 2.04 |
| IFNW1 | | cytokine | | | 2.03 |
| ORAI1 | | ion channel | | | 2.03 |
| ZBP1 | | other | | | 2.03 |
| KLRC4-KLRK1/KLRK1 | | transmembrane receptor | | | 2.03 |
| Klra7 (includes others) | | transmembrane receptor | | | 2.03 |
| IFNE | | cytokine | | 1.091 | 2.03 |
| CD2 | | transmembrane receptor | | | 2.03 |
| CYP19A1 | | enzyme | | -1.616 | 2.02 |
| LCK | | kinase | | 1.461 | 2.02 |
| PRKD1 | | kinase | | 1.387 | 2.02 |
| TARDBP | | transcription regulator | | -0.152 | 2.02 |
| PAX7 | | transcription regulator | | | 2.02 |
| EPHA2 | | kinase | | -1.091 | 2.00 |
| mir-223 | | microRNA | | -1.134 | 2.00 |
| HNRNPA2B1 | | other | | | 2.00 |
| PTPRC | | phosphatase | | | 2.00 |
| CRH | | cytokine | | -1.066 | 2.00 |
| JAG1 | | growth factor | | 0.254 | 2.00 |
| PDCD1 | | transmembrane receptor | | -1.998 | 2.00 |
| SREBF1 | | transcription regulator | | -0.653 | 1.99 |
| Stat5 dimer | | complex | | | 1.98 |
| PER1 | | transcription regulator | | | 1.98 |
| Histone h4 | | group | | | 1.95 |
| PIEZO1 | | ion channel | | | 1.95 |

| | | | | | |
|---|---|---|---|---|---|
| SH2B3 | | other | | | 1.95 |
| ZNF395 | | transcription regulator | | | 1.95 |
| AATF | | transcription regulator | | | 1.95 |
| BTG1 | | transcription regulator | | | 1.95 |
| Hbb-b2 | | other | | | 1.94 |
| DPP4 | | peptidase | | -0.686 | 1.94 |
| TNFSF9 | | cytokine | | | 1.94 |
| PLCG2 | | enzyme | | 0.492 | 1.94 |
| MAP3K5 | | kinase | | | 1.94 |
| MAP2K6 | | kinase | | | 1.94 |
| MAPK8IP1 | | other | | | 1.94 |
| GNAQ | | enzyme | | -0.698 | 1.92 |
| ZFTA-RELA | | fusion gene/product | Inhibited | -2 | 1.92 |
| EOMES | 1.788 | transcription regulator | | -0.6 | 1.91 |
| Creb | | group | | -0.555 | 1.91 |
| MAP2K1/2 | | group | | -0.555 | 1.90 |
| RORC | -2.169 | ligand-dependent nuclear receptor | | -0.93 | 1.90 |
| EHMT1 | | transcription regulator | | 0 | 1.90 |
| JUNB | | transcription regulator | | 0.099 | 1.90 |
| SCGB1A1 | | cytokine | | | 1.89 |
| Tnfsf9 | | other | | | 1.89 |
| IL2RA | 1.576 | transmembrane receptor | | | 1.89 |
| OLR1 | | transmembrane receptor | | -0.145 | 1.89 |
| INSR | | kinase | | -0.239 | 1.87 |
| AR | | ligand-dependent nuclear receptor | | -0.567 | 1.87 |
| CAV1 | | transmembrane receptor | | -0.412 | 1.87 |
| GATA1 | | transcription regulator | | -0.196 | 1.87 |
| IL24 | | cytokine | | 1.96 | 1.86 |
| SMAD2 | | transcription regulator | | -1.751 | 1.85 |
| TRIM21 | | enzyme | | | 1.84 |
| BMPER | | other | | | 1.84 |
| LGALS9 | | other | | | 1.84 |
| SOCS2 | | other | | | 1.84 |
| FASLG | 1.271 | cytokine | | -1 | 1.84 |
| CCL18 | | cytokine | | | 1.84 |
| RAC1 | | enzyme | | 0.246 | 1.84 |
| ENPP1 | | enzyme | | | 1.84 |

| | | | | | |
|---|---|---|---|---|---|
| SH3RF1 | | enzyme | | | 1.84 |
| PDE4D | | enzyme | | | 1.84 |
| Calcineurin B | | group | | | 1.84 |
| MAP3K14 | | kinase | | 0.537 | 1.84 |
| miR-494-3p (miRNAs w/seed GAAACAU) | | mature microRNA | | | 1.84 |
| NLRX1 | | other | | 0.728 | 1.84 |
| LIMIT | | other | | | 1.84 |
| SERPINA4 | | other | | | 1.84 |
| CASP4 | | peptidase | | | 1.84 |
| CTSE | | peptidase | | | 1.84 |
| POU5F1 | | transcription regulator | | 0.486 | 1.84 |
| ULBP2 | | transmembrane receptor | | | 1.84 |
| STRA6 | 2.099 | transporter | | | 1.84 |
| GJA8 | | transporter | | | 1.84 |
| TGFB3 | -2.259 | growth factor | | -1.293 | 1.83 |
| ATF4 | | transcription regulator | | -0.33 | 1.83 |
| Hsp70 | | group | | | 1.82 |
| H2AB3 (includes others) | | other | | 0 | 1.82 |
| POM121/POM121C | | other | | | 1.81 |
| TNFRSF10A | | transmembrane receptor | | | 1.81 |
| EDN1 | | cytokine | | -0.051 | 1.81 |
| PSMB11 | | peptidase | | 0.816 | 1.79 |
| POU2F1 | | transcription regulator | | | 1.79 |
| PARP2 | | enzyme | | | 1.77 |
| MAP3K3 | | kinase | | | 1.77 |
| BRD2 | | kinase | | | 1.77 |
| mir-154 | | microRNA | | | 1.77 |
| LGALS1 | | other | | 0.365 | 1.77 |
| ATG5 | | other | | 0.447 | 1.77 |
| IL31 | | other | | | 1.77 |
| TGIF1 | | transcription regulator | | | 1.77 |
| TAF4B | | transcription regulator | | | 1.77 |
| FHL2 | | transcription regulator | | -0.762 | 1.77 |
| SIGIRR | | transmembrane receptor | | | 1.77 |
| CCN1 | | other | | 0.277 | 1.76 |
| MUC1 | | other | | -1.109 | 1.76 |
| BLM | | enzyme | | | 1.75 |
| PCM1-JAK2 | | fusion gene/product | | | 1.75 |

| | | | | | |
|---|---|---|---|---|---|
| ADRB2 | | G-protein coupled receptor | | | 1.75 |
| VIPR2 | | G-protein coupled receptor | | | 1.75 |
| ROCK | | group | | 1.067 | 1.75 |
| CD3 group | | group | | 1.985 | 1.75 |
| mir-30 | | microRNA | | -0.351 | 1.75 |
| MIR17HG | 1.162 | other | | -0.254 | 1.75 |
| VTCN1 | | other | | | 1.75 |
| CLTC | | other | | | 1.75 |
| Gm20703 | | other | | | 1.75 |
| SOX18 | | transcription regulator | | | 1.75 |
| TBL1X | | transcription regulator | | | 1.75 |
| IL23R | | transmembrane receptor | | | 1.75 |
| NFAT (complex) | | complex | | | 1.74 |
| CHI3L1 | | enzyme | | | 1.74 |
| P2RY6 | | G-protein coupled receptor | | | 1.74 |
| JUN/JUNB/JUND | | group | | | 1.74 |
| SLC9A3 | | ion channel | | | 1.74 |
| SBDS | | other | | | 1.74 |
| PROCR | | other | | | 1.74 |
| ICOSLG/LOC102723996 | | other | | | 1.74 |
| CTSS | | peptidase | | | 1.74 |
| MTA2 | | transcription regulator | | | 1.74 |
| FN1 | | enzyme | | 1.414 | 1.74 |
| MTORC1 | | complex | | 1.091 | 1.73 |
| TNFAIP3 | | enzyme | | -1.067 | 1.73 |
| NOX4 | | enzyme | | 0.762 | 1.73 |
| Hsp90 | | group | | -0.147 | 1.73 |
| SAFB | | other | | 1 | 1.73 |
| DCN | | other | | 0.152 | 1.73 |
| NEDD9 | | other | | 1.342 | 1.73 |
| EIF2S1 | | translation regulator | | | 1.73 |
| ICAM1 | | transmembrane receptor | | 1.154 | 1.73 |
| MED1 | | transcription regulator | | | 1.73 |
| SOX9 | | transcription regulator | | -1.633 | 1.73 |
| DICER1 | | enzyme | | -0.846 | 1.71 |
| CAT | | enzyme | | 0.218 | 1.71 |

| | | | | | |
|---|---|---|---|---|---|
| BIRC5 | | other | | | 1.71 |
| PTPRJ | | phosphatase | Activated | 2 | 1.71 |
| CEBPE | | transcription regulator | | | 1.71 |
| AIP | | transcription regulator | | 0.218 | 1.71 |
| CBL | | transcription regulator | | | 1.71 |
| miR-203a-3p (and other miRNAs w/seed UGAAAUG) | | mature microRNA | | | 1.71 |
| SFN | | other | | | 1.71 |
| VDR | | transcription regulator | | -0.936 | 1.71 |
| RAF1 | | kinase | | -0.57 | 1.70 |
| NKX3-1 | | transcription regulator | | | 1.69 |
| NFE2L2 | | transcription regulator | | -0.958 | 1.68 |
| XCL1 | | cytokine | | | 1.68 |
| IRS1 | | enzyme | | -0.329 | 1.68 |
| BLVRA | | enzyme | | | 1.68 |
| UBA7 | | enzyme | | | 1.68 |
| DRD5 | | G-protein coupled receptor | | | 1.68 |
| GPR183 | | G-protein coupled receptor | | | 1.68 |
| HVCN1 | | ion channel | | | 1.68 |
| MAP3K12 | | kinase | | | 1.68 |
| PIM1 | | kinase | | 1.131 | 1.68 |
| mir-338 | | microRNA | | | 1.68 |
| MEMO1 | | other | | | 1.68 |
| SHANK3 | | other | | | 1.68 |
| CTSZ | | peptidase | | | 1.68 |
| HP | | peptidase | | | 1.68 |
| CAPN1 | | peptidase | | | 1.68 |
| PPP3CA | | phosphatase | | | 1.68 |
| ARID5A | | transcription regulator | | | 1.68 |
| SMARCC2 | | transcription regulator | | | 1.68 |
| MNT | | transcription regulator | | 0 | 1.68 |
| IL22RA2 | | transmembrane receptor | | | 1.68 |
| CD160 | | transmembrane receptor | | | 1.68 |
| PDYN | | transporter | | | 1.68 |
| Hbb-b1 | | transporter | | -0.218 | 1.68 |
| RASSF1 | | other | | 0.447 | 1.66 |

| | | | | | |
|---|---|---|---|---|---|
| MAPKAPK2 | | kinase | | 1.103 | 1.65 |
| GNAI2 | | enzyme | | | 1.64 |
| USP22 | | peptidase | | 0.762 | 1.63 |
| Ifn gamma | | complex | | -0.497 | 1.61 |
| CXCR3 | 1.162 | G-protein coupled receptor | | | 1.61 |
| Ubiquitin | | group | | | 1.61 |
| TRPV4 | | ion channel | | | 1.61 |
| SPHK2 | | kinase | | | 1.61 |
| mir-29 | | microRNA | | 0.356 | 1.61 |
| VTN | | other | | | 1.61 |
| CFLAR | | other | | | 1.61 |
| SLC22A5 | | transporter | | | 1.61 |
| ARID1A | | transcription regulator | | | 1.60 |
| CXCL13 | | cytokine | | | 1.60 |
| TLR2/3/4/9 | | group | | | 1.60 |
| KCNA3 | | ion channel | | | 1.60 |
| mir-217 | | microRNA | | | 1.60 |
| LRRC32 | | other | | | 1.60 |
| PHF21A | | other | | | 1.60 |
| RANBP9 | | other | | | 1.60 |
| ATAD3A | | other | | | 1.60 |
| IL9R | | transmembrane receptor | | | 1.60 |
| SLC6A1 | | transporter | | | 1.60 |
| IL7R | | transmembrane receptor | | -1.109 | 1.60 |
| BRCA1 | | transcription regulator | | -1.423 | 1.59 |
| ASPSCR1-TFE3 | | fusion gene/product | | -0.447 | 1.58 |
| collagenase | | group | | | 1.58 |
| Fc gamma receptor | | group | | | 1.58 |
| MAPK11 | | kinase | | | 1.58 |
| TIRAP | | other | | | 1.58 |
| ZEB1 | | transcription regulator | | 0.152 | 1.58 |
| MYCN | | transcription regulator | | 1.795 | 1.57 |
| LAMA4 | | enzyme | | 1.342 | 1.56 |
| TAB1 | | enzyme | | | 1.55 |
| Fgf | | group | | | 1.55 |
| LEF1 | | transcription regulator | | 1.342 | 1.55 |
| STAR | | transporter | | 0 | 1.55 |
| SCD | | enzyme | | 0.447 | 1.54 |

| | | | | | | |
|---|---|---|---|---|---|---|
| Stat1 dimer | | complex | | | | 1.54 |
| Adaptor protein 2 | | complex | | | | 1.54 |
| IL25 | | cytokine | | | 0.577 | 1.54 |
| TNFSF18 | | cytokine | | | | 1.54 |
| Ifnz (includes others) | | cytokine | | | | 1.54 |
| Fcor | | enzyme | | | | 1.54 |
| PDE3B | | enzyme | | | | 1.54 |
| ADAR | | enzyme | | | | 1.54 |
| ENTPD1 | | enzyme | | | | 1.54 |
| SLC9A1 | | ion channel | | | | 1.54 |
| LAMTOR1 | | other | | | | 1.54 |
| Traj18 | | other | | | | 1.54 |
| LNX2 | | other | | | | 1.54 |
| OSTM1 | | other | | | | 1.54 |
| MMP7 | | peptidase | | | | 1.54 |
| TRIM28 | | transcription regulator | | | | 1.54 |
| NEUROG1 | | transcription regulator | | | 0 | 1.54 |
| KLF7 | | transcription regulator | | | | 1.54 |
| HIVEP3 | | transcription regulator | | | | 1.54 |
| ELF2 | | transcription regulator | | | | 1.54 |
| KLRD1 | | transmembrane receptor | | | | 1.54 |
| SLC7A5 | | transporter | | | | 1.54 |
| SYVN1 | | transporter | | | 0.816 | 1.53 |
| THPO | | cytokine | | | -0.152 | 1.53 |
| Nfat (family) | | group | | | 1 | 1.53 |
| TYK2 | | kinase | | | | 1.53 |
| OSCAR | | other | | | | 1.53 |
| MALT1 | | peptidase | | | | 1.53 |
| NOTCH4 | | transcription regulator | | | | 1.53 |
| ZBTB7A | | transcription regulator | | | | 1.53 |
| DAXX | | transcription regulator | | | | 1.53 |
| SGK1 | -1.153 | kinase | | | -1.982 | 1.52 |
| CEBPA | -1.947 | transcription regulator | | | 1.446 | 1.52 |
| TRAF3 | | enzyme | | | -0.2 | 1.52 |
| LIPE | | enzyme | | | | 1.51 |
| IKK (complex) | | complex | | | | 1.50 |
| Notch | | group | | | 0.339 | 1.50 |
| mir-126 | | microRNA | | | | 1.50 |
| XBP1 | | transcription regulator | Inhibited | | -2.176 | 1.50 |

| | | | | | |
|---|---|---|---|---|---|
| PHB | | transcription regulator | | | 1.50 |
| PROX1 | | transcription regulator | | | 1.50 |
| EGR1 | | transcription regulator | | 0.629 | 1.49 |
| Growth hormone | | group | | -0.57 | 1.48 |
| PDGF BB | | complex | | -1.018 | 1.48 |
| ALOX12 | | enzyme | | | 1.48 |
| DOK1 | | kinase | | | 1.48 |
| SERPINA3 | | other | | | 1.48 |
| FYB1 | | other | | | 1.48 |
| ELN | | other | | | 1.48 |
| MAP2K1 | | kinase | | -0.928 | 1.47 |
| TRIB3 | -1.217 | kinase | | | 1.47 |
| miR-221-3p (and other miRNAs w/seed GCUACAU) | | mature microRNA | | | 1.47 |
| NCSTN | | peptidase | | | 1.47 |
| RBCK1 | | transcription regulator | | | 1.47 |
| 26s Proteasome | | complex | | 0.634 | 1.45 |
| GATA3 | | transcription regulator | Inhibited | -2.406 | 1.45 |
| IL20 | | cytokine | | | 1.45 |
| RNASE2 | | enzyme | | | 1.45 |
| GNB2 | | enzyme | | | 1.45 |
| AKT1 | | kinase | | 0.966 | 1.45 |
| VIP | | other | | -1.117 | 1.45 |
| TANK | | other | | | 1.45 |
| ZEB2 | | transcription regulator | | 0 | 1.45 |
| EBF3 | | transcription regulator | | | 1.45 |
| SS18 | | transcription regulator | | | 1.45 |
| ELF4 | | transcription regulator | | | 1.45 |
| TREM2 | | transmembrane receptor | | -1 | 1.45 |
| EBF1 | | transcription regulator | | -0.689 | 1.45 |
| SP1 | | transcription regulator | | -0.823 | 1.45 |
| WNT5A | | cytokine | | -0.218 | 1.44 |
| SPRY2 | | other | | 1.342 | 1.44 |
| ISGF3 | | complex | | | 1.43 |
| DHCR24 | | enzyme | | | 1.43 |
| CRY2 | | enzyme | | | 1.43 |
| CRY1 | | enzyme | | | 1.43 |
| CYBB | | enzyme | | | 1.43 |

| | | | | | |
|---|---|---|---|---|---|
| Sos | | group | | | 1.43 |
| STAT | | group | | | 1.43 |
| RIPK2 | | kinase | | 0.991 | 1.43 |
| PIK3R2 | | kinase | | | 1.43 |
| RPS6KA4 | | kinase | | | 1.43 |
| mir-130 | | microRNA | | | 1.43 |
| ADAP1 | | other | | | 1.43 |
| DANCR | | other | | | 1.43 |
| HLA-A | -2.549 | other | | | 1.43 |
| CDC37 | | other | | | 1.43 |
| KPNA2 | | other | | | 1.43 |
| ATXN3 | | peptidase | | | 1.43 |
| PON1 | | phosphatase | | | 1.43 |
| PPP5C | | phosphatase | | | 1.43 |
| DUSP14 | | phosphatase | | | 1.43 |
| SMARCA2 | | transcription regulator | | | 1.43 |
| SMAD7 | | transcription regulator | Activated | 2.408 | 1.43 |
| SOX5 | | transcription regulator | | | 1.43 |
| PA2G4 | | transcription regulator | | | 1.43 |
| HOXA1 | | transcription regulator | | | 1.43 |
| PRDM16 | | transcription regulator | | | 1.43 |
| RBP1 | | transporter | | | 1.43 |
| MTTP | | transporter | | | 1.43 |
| VEGFA | | growth factor | | 0.471 | 1.43 |
| Smad2/3 | | group | | -1.067 | 1.43 |
| INSIG1 | | other | Inhibited | -2 | 1.42 |
